# Tocilizumab treatment leads to early resolution of lymphopenia and myeloid dysregulation in patients hospitalized with COVID-19

**DOI:** 10.1101/2022.10.27.514096

**Authors:** Haridha Shivram, Jason A. Hackney, Carrie M Rosenberger, Anastasia Teterina, Aditi Qamra, Olusegun Onabajo, Jacqueline McBride, Fang Cai, Min Bao, Larry Tsai, Aviv Regev, Ivan O. Rosas, Rebecca N. Bauer

## Abstract

High interleukin (IL)-6 levels are associated with more severe clinical manifestations in patients hospitalized with COVID-19, but the complex role of IL-6 in antiviral and inflammatory processes has made it difficult to decipher its involvement in the disease. IL-6 receptor blockade by tocilizumab (anti-IL6R; Actemra) is used globally for the treatment of severe COVID-19, yet a molecular understanding of the therapeutic benefit remains unclear. We characterized the immune profile and identified cellular and molecular pathways directly modified by tocilizumab in peripheral blood samples collected from patients enrolled in the COVACTA study, a phase 3, randomized, double-blind, placebo-controlled trial that assessed the efficacy and safety of tocilizumab in hospitalized patients with severe COVID-19 pneumonia. We identified factors predicting disease severity and clinical outcomes, including markers of inflammation, lymphopenia, myeloid dysregulation, and organ injury. Proteomic analysis confirmed a pharmacodynamic effect for tocilizumab in addition to identifying novel pharmacodynamic biomarkers. Transcriptomic analysis revealed that tocilizumab treatment leads to faster resolution of lymphopenia and myeloid dysregulation associated with severe COVID-19, indicating greater anti-inflammatory activity relative to standard of care and potentially leading to faster recovery in patients hospitalized with COVID-19.

**One sentence summary:** Interleukin-6 receptor blockade with tocilizumab accelerated resolution of myeloid dysfunction and lymphopenia in patients hospitalized with COVID-19

## Introduction

Coronavirus disease 2019 (COVID-19) has caused millions of deaths globally. Case numbers continue to rise despite increasing availability of vaccines as more infectious severe acute respiratory syndrome coronavirus 2 (SARS-CoV-2) variants emerge. The course of COVID-19 is heterogeneous, ranging from asymptomatic to severe illness leading to acute respiratory distress syndrome (ARDS), multiorgan system failure and death (*1*). Moreover, some COVID-19 survivors continue to display postacute COVID-19 cardiovascular and pulmonary complications (*2–3*).

Patients with severe COVID-19 exhibit immune dysregulation characterized by hyperinflammation and lymphopenia, with evidence of myeloid cell activation and impaired T cell function (*4*). Similar to other causes of ARDS, high levels of the pro-inflammatory cytokine interleukin-6 (IL-6) have been associated with more severe clinical manifestations in patients hospitalized with COVID-19, and elevated IL-6 is prognostic for worse clinical outcomes (*5–8*). IL-6 plays a critical role in driving the pro-inflammatory acute phase of antiviral responses, as well as in switching the inflammatory response from an acute, neutrophil-driven response to an inflammatory response characterized by recruitment of monocytes and differentiation of macrophages (*9*). The complex role of IL-6 has made it difficult to decipher the involvement of IL-6 in the pathology of COVID-19, and spurred initial concerns that anti-inflammatory treatments such as IL-6 receptor (IL-6R) blockade could exacerbate disease or increase risk for secondary infections, as was observed for corticosteroid treatment in critically ill patients with influenza (*10*).

Since the beginning of the pandemic, more than one million people hospitalized with COVID-19 have been treated with tocilizumab, an IL-6R alpha antibody. Multiple studies have assessed the benefit of tocilizumab in patients hospitalized with COVID-19 (*11–20*), and have individually shown mixed results for efficacy, likely due to heterogeneity of the patient population, timing of treatment, shifting standard of care (eg, systemic corticosteroids), and insufficient sample size. A meta-analysis conducted by the World Health Organization (WHO) demonstrated reduced mortality, shortened time to hospital discharge, and reduced mechanical ventilation (MV) in patients treated with tocilizumab across the 19 studies considered (*2*). Based on these findings, tocilizumab was included in the WHO treatment recommendations for severe COVID-19 and subsequently tocilizumab has been approved in more than 30 countries for the treatment of patients hospitalized with severe COVID-19. As of December 2022, tocilizumab is the first FDA-approved monoclonal antibody to treat patients hospitalized with severe COVID-19 who are receiving systemic corticosteroids and require supplemental oxygen, noninvasive or invasive MV, or extracorporeal membrane oxygenation.

To support the clinical use of tocilizumab for COVID-19 and advance understanding of IL-6R blockade for treatment of severe pneumonia and ARDS, we studied the effects of tocilizumab treatment in patients with COVID-19 by analyzing proteomic and transcriptomic data from peripheral blood samples in a post hoc analysis of the COVACTA study. COVACTA was a global, double-blind, randomized, placebo-controlled, phase 3 trial of 452 patients hospitalized with COVID-19 randomized to tocilizumab or placebo treatment that enrolled patients early in the pandemic (April 3, 2020 through May 28, 2020) before widespread use of corticosteroids or effective antivirals for the treatment of COVID-19 (13). The study did not meet its primary end point of improved clinical status on a 7-category ordinal scale at day 28, and no difference was observed in mortality at day 28 with tocilizumab versus placebo. However, patients treated with tocilizumab showed a nominal reduction in median time to hospital discharge or ready for discharge compared with placebo (20 vs 28 days, respectively). This study provides a unique opportunity to evaluate the mechanism of action for IL-6R blockade with tocilizumab in patients with COVID-19 and determine the basis for its benefit on improved time to clinical recovery independent of a combined effect with corticosteroids or antivirals.

Here, we characterized the effect of tocilizumab on immune aberrations associated with COVID-19 disease progression, pathways downstream of interleukin-6 signaling, and organ injury protein signatures in patients hospitalized with COVID-19 pneumonia. First, we defined biomarkers associated with severity and disease outcome, recapitulating previously described mechanisms of COVID-19 pathogenesis in this large clinical cohort despite potential confounders (eg, inclusion/exclusion criteria). Second, we confirm tocilizumab pharmacodynamic activity and identified novel pharmacodynamic biomarkers that could be monitored in future trials of IL-6R blockade for new indications such as ARDS. Third, we evaluated effects of tocilizumab on the pathways associated with COVID-19 severity or worse clinical outcome and those that were not affected by IL-6R blockade. Our results show that tocilizumab treatment leads to faster resolution of lymphopenia and myeloid dysregulation associated with severe COVID-19. We also identified cell subsets and pathways that remain dysregulated in patients who died despite treatment that may be targets for novel treatments. In totality, these findings from a randomized placebo-controlled trial describe the molecular underpinnings supporting tocilizumab efficacy in patients with COVID-19 and delineate the mechanism of IL-6R blockade from its combinatorial effect with corticosteroids. Further, these findings yield impactful hypotheses for future clinical trials to evaluate the efficacy of IL-6R blockade in patients with severe pneumonia or ARDS irrespective of etiology.

## Results

### Design of serum protein and whole blood RNA profiling at baseline and during treatment with tocilizumab in COVACTA

COVACTA (ClinicalTrials.gov, NCT04320615) was a randomized, placebo-controlled, double-blind, global, multicenter, phase 3 trial investigating the efficacy and safety of tocilizumab versus placebo in patients hospitalized with severe COVID-19 pneumonia (**Fig. 1**) (15). Patients were stratified by geographic region (North America/Europe) and MV (yes/no) and randomly assigned (2:1) to receive a single intravenous tocilizumab infusion (8 mg/kg body weight) or placebo in addition to standard care. Patients who showed disease worsening or no clinical improvement were allowed to receive a second tocilizumab/placebo infusion 8 to 24 hours after the first dose. Disease severity was assessed at baseline and after dosing throughout the study using a 7-category ordinal scale with increasing severity from 1 to 7 as follows: 1, discharged or ready for discharge; 2, non–intensive care unit (ICU) hospital ward, not requiring supplemental oxygen; 3, non–ICU hospital ward, requiring supplemental oxygen; 4, ICU or non–ICU hospital ward, requiring noninvasive ventilation or high-flow oxygen; 5, ICU, requiring intubation and mechanical ventilation; 6, ICU, requiring extracorporeal membrane oxygenation or mechanical ventilation and additional organ support; and 7, death (15). At baseline, “Moderate” COVID-19 was defined as an ordinal scale score <4 and “severe” COVID-19 was defined as ordinal scale score ≥4. Refer to the methods of that paper for more details.

**Fig 1.**
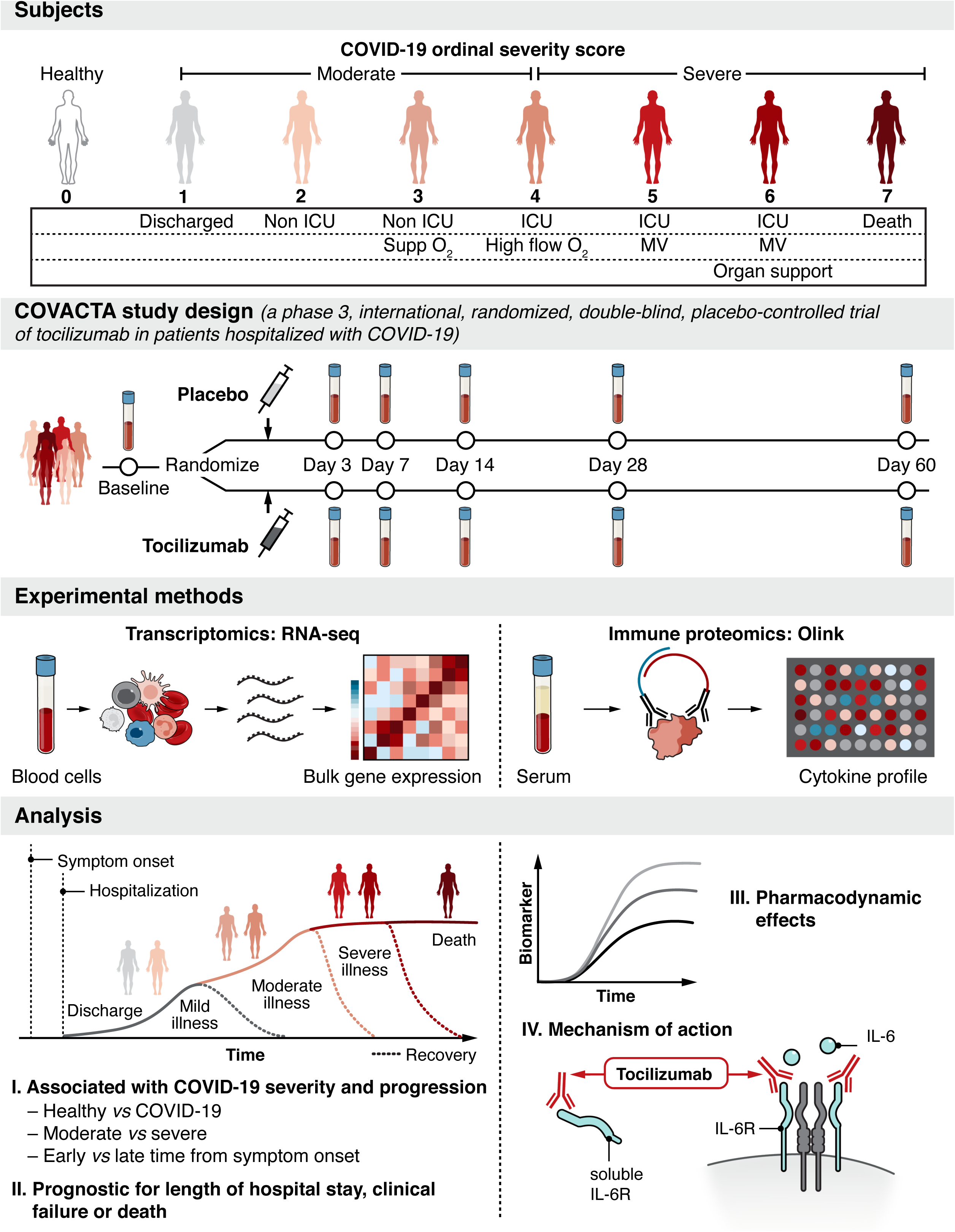
Serum protein and whole blood RNA profiling from hospitalized patients with COVID-19 enrolled in the COVACTA study at baseline and during treatment with tocilizumab. Disease severity was assessed at baseline and after dosing using a 7-category ordinal scale. At baseline, “moderate” COVID-19 was defined as an ordinal scale score <4 and “severe” COVID-19 was defined as ordinal scale score ≥4. The present analysis included an external control group using peripheral blood from healthy adults. Patients enrolled in COVACTA received a single intravenous tocilizumab infusion (8 mg/kg) or placebo in addition to standard care. Serum and peripheral whole blood collected from COVACTA patients at baseline (before dosing) and up to 60 days after dosing were profiled by a proteomics (Olink) and transcriptomics (RNA-Seq) assay. Key immune aberrations associated with COVID-19, time from symptom onset, and disease severity were identified at baseline and assessed for prognostic association with time to hospital discharge/ready for discharge, mortality, and time to clinical failure. To understand mechanism of action, the pharmacodynamic effect of tocilizumab and the response of severity associated biomarkers to tocilizumab treatment were assessed in longitudinal samples. ICU, intensive care unit; IL-6R, interleukin 6 receptor; MV, mechanical ventilation; supp, supplemental.

For the present analysis, serum and peripheral whole blood collected from COVACTA patients at baseline (before dose) and up to 60 days after dosing were profiled by a proteomics (Olink) and transcriptomics (RNA-Seq) assay, along with an external control group comprising peripheral blood samples from healthy adult volunteers obtained from the Genentech Samples for Science program (**Fig. 1**, **Table 1**). First, we identified key immune aberrations associated with COVID-19 severity and disease progression at baseline, relationship with time from symptom onset, and associations of genes and protein biomarkers with measures of clinical outcomes – time to hospital discharge/ready for discharge, mortality, and time to clinical failure (defined as any one of the following: death, discontinuation from trial participation during hospitalization, initiation of mechanical ventilation, or ICU transfer or a 1-category worsening of clinical status in patients who were receiving mechanical ventilation or who were in the ICU at baseline; the initiation of mechanical ventilation among patients who were not receiving mechanical ventilation at randomization; the incidence of ICU transfer among patients who were not in an ICU at baseline; and the duration of ICU stay) (*15*). Second, to understand mechanism of action, we assessed the pharmacodynamic effect of tocilizumab treatment and evaluated the response of the aforementioned severity-associated biomarkers to tocilizumab treatment in longitudinal samples and compared responses in patients who survived to those who died during the course of the study.

**Table 1.** Baseline characteristics of patients included in the analyses.

| Characteristic | RNA Sequencing Analysis |  | Olink Proteomic Analysis |  |
| --- | --- | --- | --- | --- |
|  | Tocilizumab<br>*N* = 271 | Placebo<br>*N* = 133 | Tocilizumab<br>*N* = 266 | Placebo<br>*N* = 122 |
| Sex |  |  |  |  |
| Male | 189 (69.7) | 95 (71.4) | 187 (70.3) | 88 (72.1) |
| Female | 82 (30.3) | 38 (30.3) | 79 (29.7) | 34 (27.9) |
| Age (years) |  |  |  |  |
| 18-64 | 148 (54.6) | 75 (56.4) | 146 (54.9) | 67 (54.9) |
| 65-84 | 110 (40.6) | 55 (41.4) | 107 (40.2) | 53 (40.2) |
| ≥85 | 13 (4.8) | 3 (2.3) | 13 (4.9) | 2 (1.6) |
| Corticosteroid treatment<sup>a</sup> | 54 (19.9) | 39 (29.3) | 51 (19.2) | 33 (27.0) |
| Antiviral treatment<sup>a</sup> | 65 (24.0) | 39 (29.3) | 65 (24.4) | 33 (27.0) |
| Days from symptom onset |  |  |  |  |
| ≤10 | 116 (43.3) | 70 (53) | 117 (44.5) | 65 (53.7) |
|  | 152 (56.7) | 62 (47) | 146 (55.5) | 56 (46.3) |
| >10 |  |  |  |  |
| Ordinal scale score |  |  |  |  |
| 2 |  |  |  |  |
| 3 | 9 (3.3) | 6 (4.5) | 9 (3.4) | 5 (4.1) |
| 4 | 69 (25.5) | 40 (30.1) | 70 (26.3) | 38 (31.1) |
| 5 | 81 (29.9) | 33 (24.8) | 79 (29.7) | 33 (27.0) |
| 6 | 43 (15.9) | 15 (11.3) | 43 (16.2) | 13 (10.7) |
| 7 | 69 (25.5) | 38 (28.6) | 65 (24.4) | 32 (26.2) |
|  | 0 | 1 (0.8) | 0 | 1 (0.8) |

All data are presented as n (%).
<sup>a</sup>Corticosteroid and/or antiviral use at baseline (study day –7 to first dose of study treatment) or at any time during the study. Corticosteroid use included only systemic use, and antiviral treatment included lopinavir-ritonavir, remdesivir, lopinavir, ritonavir, chloroquine, hydroxychloroquine, and hydroxychloroquine sulfate.

**Table 2.**
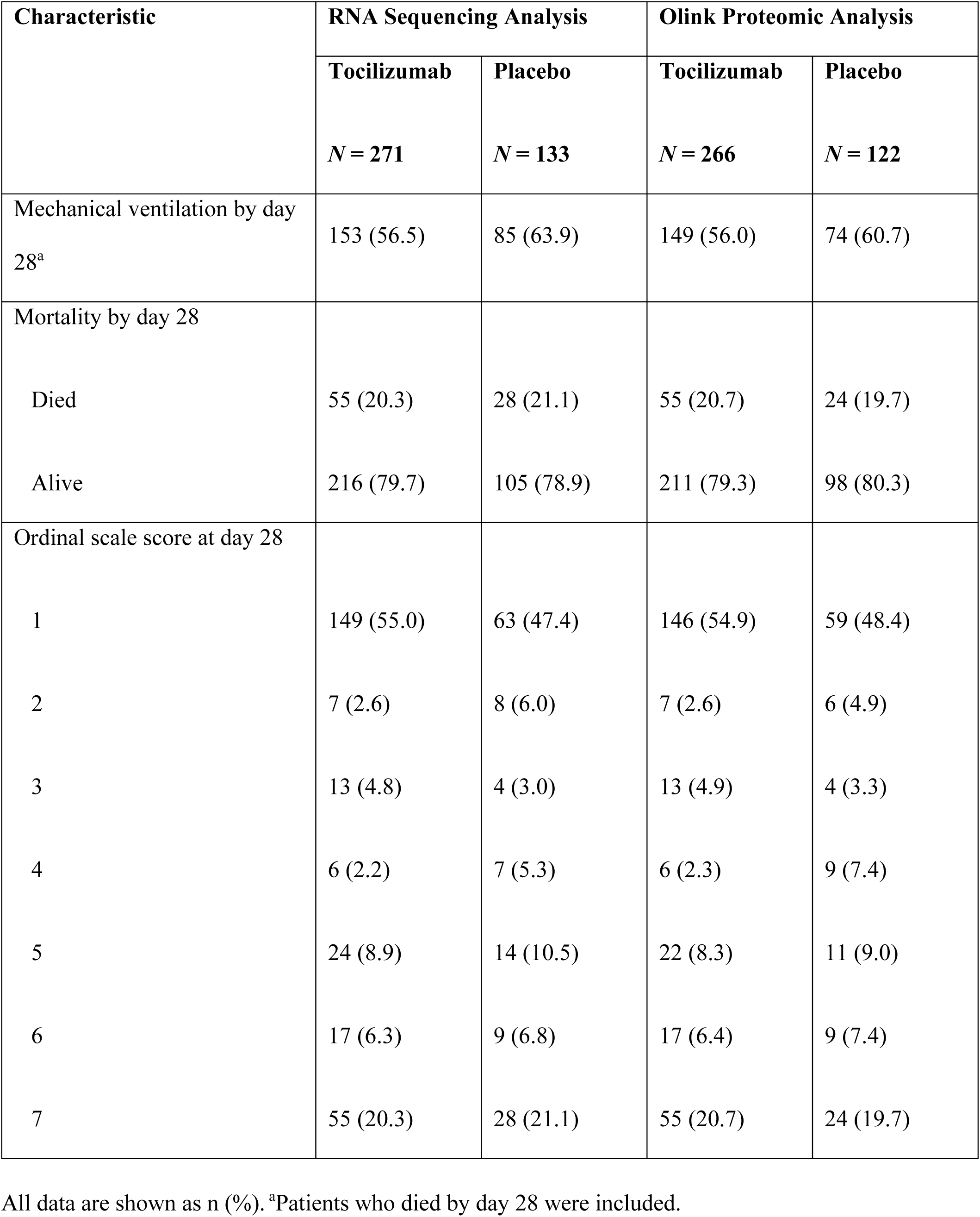
Patient characteristics at day 28 of the study.

| Characteristic | RNA Sequencing Analysis |  | Olink Proteomic Analysis |  |
| --- | --- | --- | --- | --- |
|  | Tocilizumab<br>*N* = 271 | Placebo<br>*N* = 133 | Tocilizumab<br>*N* = 266 | Placebo<br>*N* = 122 |
| Mechanical ventilation by day 28<sup>a</sup> | 153 (56.5) | 85 (63.9) | 149 (56.0) | 74 (60.7) |
| Mortality by day 28 |  |  |  |  |
| Died | 55 (20.3) | 28 (21.1) | 55 (20.7) | 24 (19.7) |
| Alive | 216 (79.7) | 105 (78.9) | 211 (79.3) | 98 (80.3) |
| Ordinal scale score at day 28 |  |  |  |  |
| 1 | 149 (55.0) | 63 (47.4) | 146 (54.9) | 59 (48.4) |
| 2 | 7 (2.6) | 8 (6.0) | 7 (2.6) | 6 (4.9) |
| 3 | 13 (4.8) | 4 (3.0) | 13 (4.9) | 4 (3.3) |
| 4 | 6 (2.2) | 7 (5.3) | 6 (2.3) | 9 (7.4) |
| 5 | 24 (8.9) | 14 (10.5) | 22 (8.3) | 11 (9.0) |
| 6 | 17 (6.3) | 9 (6.8) | 17 (6.4) | 9 (7.4) |
| 7 | 55 (20.3) | 28 (21.1) | 55 (20.7) | 24 (19.7) |

All data are shown as n (%). <sup>a</sup>Patients who died by day 28 were included.

### Elevated serum levels of proteins indicative of inflammation, myeloid dysfunction, and organ injury in patients with severe COVID-19

To identify molecular changes associated with COVID-19 severity at baseline, we first compared the levels of serum proteins in patients enrolled in the COVACTA study classified at baseline as “moderate” (ordinal scale <4) vs “severe” (ordinal scale ≥4). Of 1472 measured proteins, 230 (224 high confidence measurements based on our quality control [QC] filter) had higher levels and 21 had lower serum levels in patients with severe vs moderate COVID-19 at baseline (Benjamini-Hochberg false discovery rate (FDR) <0.05, Limma; logFC ≥0.5 or logFC ≤−0.5, **Fig. 2A-B, Table S1**). Proteins having higher levels in patients with severe COVID-19 included several inflammatory cytokines and other proteins involved in IL6-Janus kinase-signal transducer and activator of transcription (JAK/STAT) and tumor necrosis factor (TNF) signaling (**Fig. 2A-B**); multiple markers of inflammatory myeloid cells (eg, OLR1, OSM, and CLEC5A) (**Fig. 2A-B**); and proteins involved in edema, coagulation, and tissue repair (**Fig. 2A-B**).

**Fig 2.**
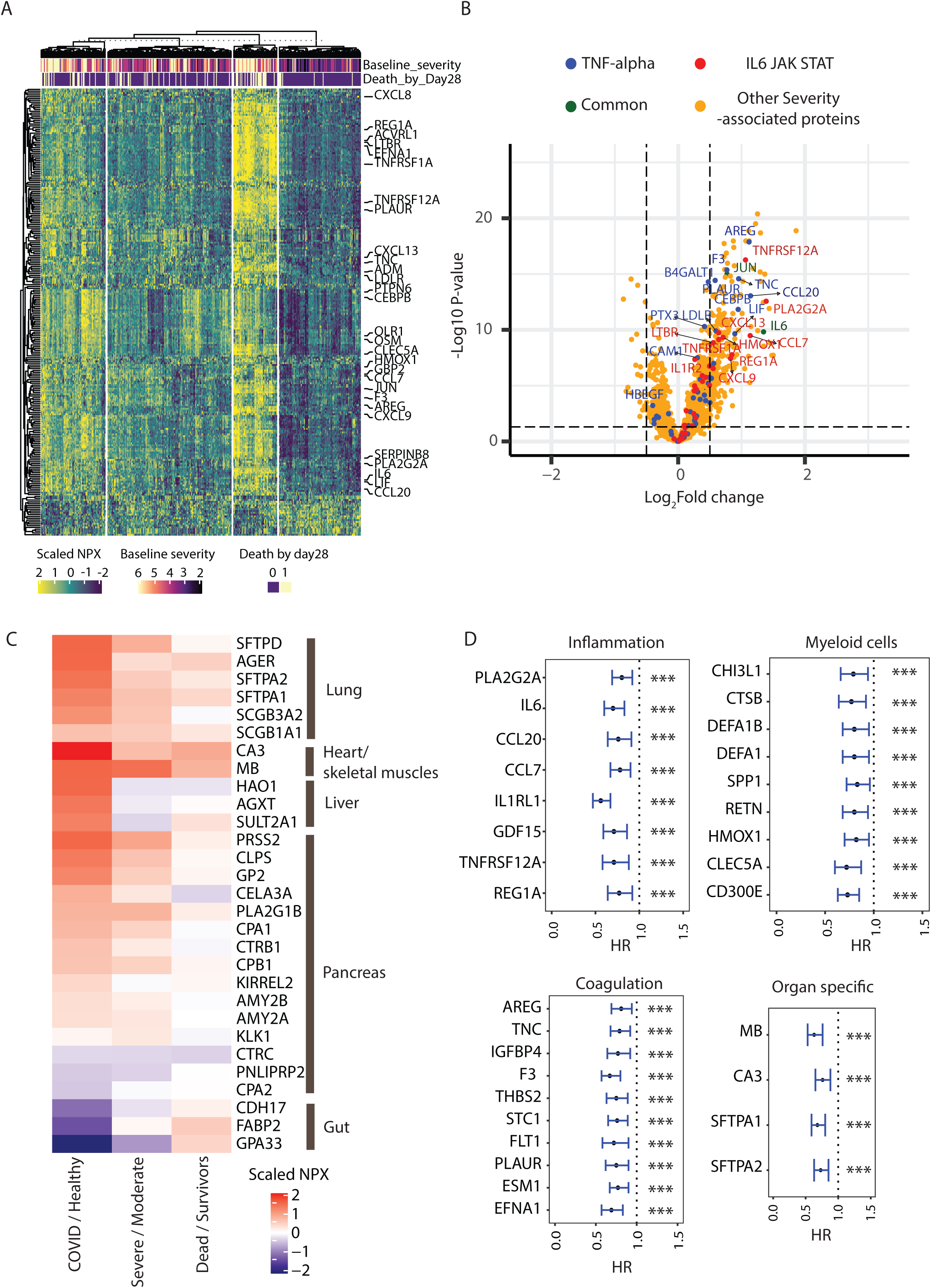
Protein biomarkers associated with COVID-19 severity and disease progression. **(A)** Heatmap showing serum levels in NPX units of 245 proteins differentially expressed based on COVID-19 baseline severity (224 proteins with higher and 21 with lower levels in severe patients respectively). 6 proteins were filtered out before plotting due to lower measurement quality. Rows represent proteins and columns represent samples measured at baseline. Columns (*n* = 391) are clustered using K-means clustering. Specific proteins involved in inflammatory pathways are highlighted by name. **(B)** Volcano plot showing differentially expressed proteins between severe and moderate COVID-19. Proteins involved in the specified pathways are highlighted by colors. **(C)** Heatmap showing differential expression of organ-specific proteins between COVID (*n* = 388) vs healthy (*n* = 59) (Column 1), baseline severity ≥4 (Severe, *n* = 266) vs <4 (Moderate, *n* = 122) (Column 2), and mortality at day 28 yes (Dead, *n* = 79) vs no (Survivors, *n* = 309) (Column 3). **(D)** Proteins prognostic for worse clinical outcome (time to hospital discharge) identified using Cox proportional hazard model. Proteins are arranged in decreasing order of fold change between severe vs moderate cases. Error bars represent 95% confidence intervals. IL, interleukin; JAK STAT, Janus kinase-signal transducer and activator of transcription; TNF, tumor necrosis factor.

Moreover, organ-specific proteins were also present in serum at levels different from patients with severe COVID-19, moderate COVID-19, or healthy volunteers, which suggests COVID-19-associated organ injury or decline in organ function (*21*). To this end, we annotated 29 of the 1472 measured proteins as being restricted to specific organ systems, including lung, liver, heart/skeletal muscle, gut, and pancreas (see methods for details). Proteins specific to lung, cardiac/skeletal muscle, and liver were elevated in patients with COVID-19 vs healthy controls and patients with severe vs moderate COVID-19 (**Fig. 2C; Fig. S1,** Benjamini-Hochberg FDR <0.05). These included severity-associated increases in the levels of lung-related surfactant proteins (SFTPA1/2 and D) and receptors for advanced glycation end-products (RAGE/AGER), associated, respectively, with a decline in pulmonary function and lung epithelial cell injury (*22, 23*); increases in cardiac-related proteins in patients with severe COVID-19 vs moderate or healthy volunteers, including carbonic anhydrase III (CA3), a cardiac/skeletal muscle-specific protein, and myoglobin (MB), expressed in skeletal, smooth, and cardiac muscle (**Fig. 2C; Fig. S1**); increases in liver-specific protein markers in patients with COVID-19 vs healthy volunteers (but not severe vs moderate), which were also positively correlated with established clinical liver damage markers (aspartate aminotransferase [AST] and alanine aminotransferase [ALT]), consistent with previous reports of liver injury in patients with severe COVID-19 (*23*) (**Fig. S1; Fig. S2**); and significant increases in pancreas markers CLPS, PRSS2, PLA2G1B, and GP2 in COVID-19 vs healthy controls and in severe vs moderate COVID-19. Conversely, there were lower levels of gut-associated proteins in patients with severe vs moderate COVID-19 or COVID-19 vs healthy volunteers (**Fig. 2C**).

Notably, 515 proteins remained elevated in COVID-19 survivors at day 60 compared to healthy volunteers, including 119 (51%) of the 230 severity-associated proteins (Benjamini-Hochberg FDR <0.05; logFC ≥0.5). These include proteins associated with heart (CA3), liver (HAO1), and lung injury (SFTPD) or pulmonary fibrosis (CCL18 and OPN), inflammatory markers of myeloid origin (eg, DEFA1, PADI4, AZU1, OSM) and tissue repair proteins (eg, PLAUR, IGFBP4, AREG, TNC) (**Table S1**).

### Serum proteins associated with baseline severity are prognostic for worse clinical progression

We next asked whether proteins associated with baseline severity were also prognostic for worse clinical outcomes. Comparison of protein levels at baseline between patients progressing to death vs survivors by day 28 identified 27 proteins that are both correlated with baseline disease severity and further elevated in patients who died (Limma model controlled for baseline severity and age; Benjamini-Hochberg FDR <0.05; logFC ≥0.5, **Fig. S3A, labelled red dots**). Moreover, hazard modeling identified 533 proteins associated with increased (hazard ratio >1) likelihood of clinical failure and 261 proteins associated with lower (hazard ratio <1) likelihood of hospital discharge (model controlled for baseline severity, age and treatment arm; Benjamini-Hochberg FDR <0.05, **Table S2**). Of these, 246 proteins were associated with both increased likelihood of clinical failure and lower likelihood of hospital discharge. The prognostic proteins included severity-associated proteins related to inflammation, myeloid cells, edema, coagulation and tissue repair, and proteins specific to heart and lung (Benjamini-Hochberg FDR <0.05) associated with either lower likelihood of hospital discharge (**Fig. 2D**) or increased clinical failure (**Fig. S3B**).

Since the time of hospitalization of patients varied with respect to time from symptom onset (median = 11 days [range 1-50]), the observed prognostic signals could be impacted by the stage of disease and/or timing of intervention. We thus repeated our hazard modeling adjusting for time from symptom onset at baseline and found most proteins remained prognostic for worse clinical outcomes (**Table S2**). Since ∼25% of the enrolled patients were also treated with corticosteroids, we reassessed the prognostic properties of the identified proteins and found the prognostic effect of the proteins to be maintained even after controlling for corticosteroid treatment (data not shown). Taken together, these data suggest that biomarkers indicative of a heightened inflammatory state and consequential organ injury are not only associated with severe COVID-19, but are also prognostic for worse clinical progression.

### Increased expression signatures of lymphopenia and myeloid dysregulation in whole blood from patients with severe COVID-19

To define molecular pathways modulated in patients with COVID-19, we first identified RNA expression signatures measured at baseline that are correlated with COVID-19 disease severity. In baseline RNA-seq data, 4755 genes were differentially expressed in severe vs moderate COVID-19 (1900 upregulated; 2855 downregulated; Benjamini-Hochberg FDR <0.05, DESeq2 and log_2_FoldChange ≥0.5 or log_2_FoldChange ≤−0.5). Gene set enrichment analysis (GSEA) on genes ranked by difference in expression (DESeq2 derived Log_2_fold change using all genes) showed increased expression of inflammation, monocyte, heme metabolism, coagulation, and platelet activation genes and decreased expression of lymphocyte–biology-related gene signatures in severe patients (**Fig. 3A**). Blood collected at baseline from patients progressing to death by day 28 had increased expression of antiviral and inflammatory pathways (DESeq2 model controlled for baseline severity and age, **Fig. 3B**). Concordantly, higher expression of antiviral and inflammatory pathways at baseline was significantly (Cox proportional hazards model controlled for baseline severity, age, and treatment arm; Benjamini-Hochberg FDR <0.05, **Table S3**) prognostic for slower recovery (increased time to hospital discharge/ready for discharge) and faster time to clinical failure (**Fig. 3C**), even when adjusting for time from symptom onset at baseline (**Table S3**).

**Fig 3.**
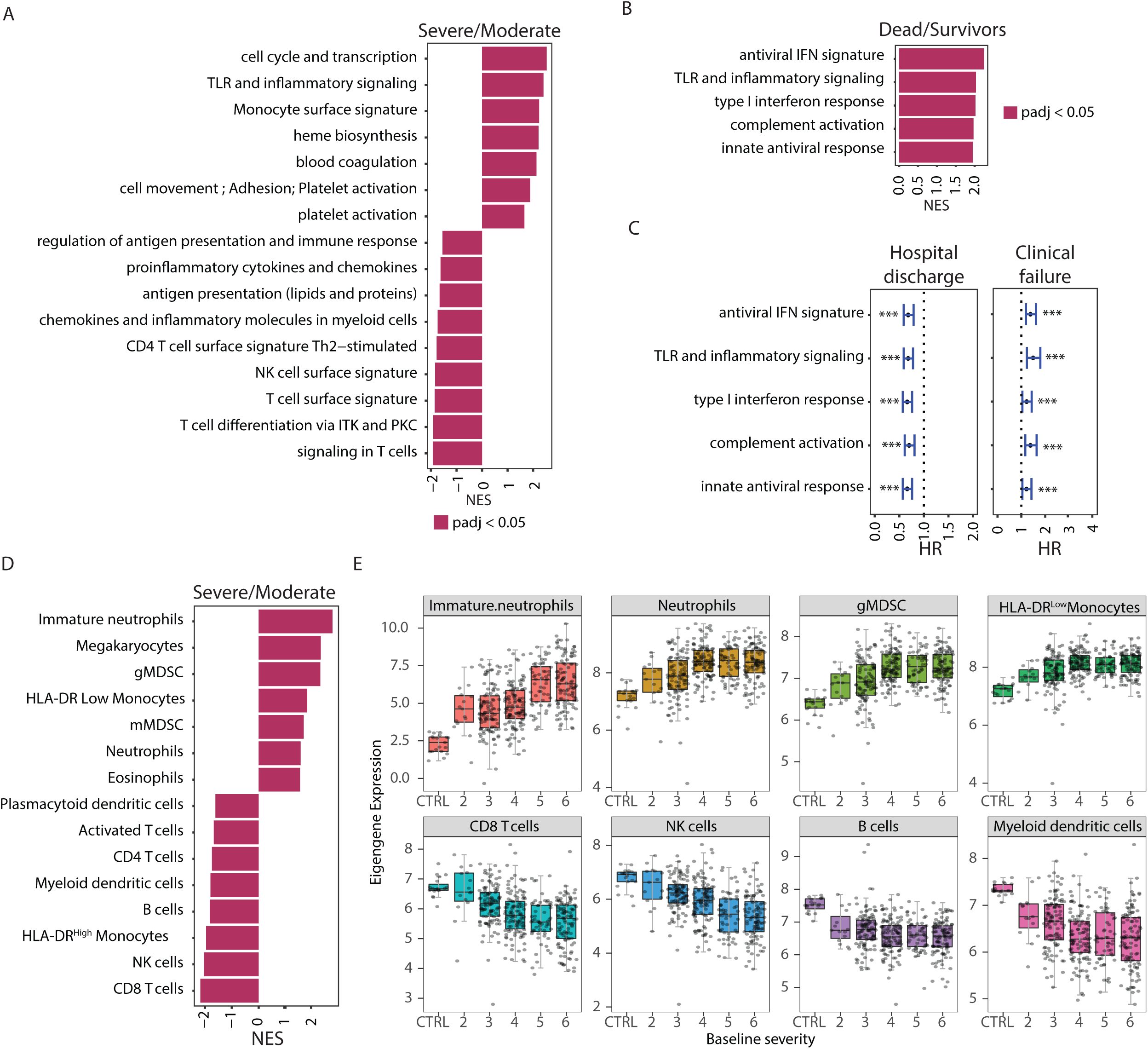
Transcriptomic signatures associated with COVID-19 severity and disease progression. **(A)** Fast gene set enrichment analysis of differentially expressed genes between cases with baseline severity ≥4 (Severe, *n* = 280) and <4 (Moderate, *n* = 124) at baseline. Higher NES represents upregulation of the indicated immune pathways. **(B)** FGSEA analysis of differentially expressed genes between cases associated with higher mortality (death by day 28, *n* = 83) and survivors (*n* = 321). Higher NES represents upregulation of the indicated immune pathway. **(C)** Forest plot showing hazard ratios with 95% confidence ratios identified using Cox proportional hazard model depicting time to hospital discharge (left) and clinical failure (right) from COVID-19 for immune pathways shown in Fig 3B. ***Represents Benjamini-Hochberg adjusted *P* value <0.05. Error bars represent 95% confidence intervals. **(D)** Same as 3A where higher NES represents up regulation of blood cellularity signatures. **(E)** Box plots showing changes in eigengene values of blood cell type signatures across different baseline severity scales of COVID-19 patients. The X-axis represents the baseline severity scales: 2 (*n* = 15), 3, (*n* = 109), 4 (*n* = 114), 5 (*n* = 58), 6 (*n* = 107) and healthy controls (CTRL, *n* = 19). CD, cluster of differentiation; gMDSC, granulocytic myeloid-derived suppressor cells; HLA, human leukocyte antigen; HR, hazard ratio; IFN, interferon; ITK, IL-2 inducible T cell kinase; mMDSC, monocytic myeloid-derived suppressor cells; NES, normalized enrichment score; NK, natural killer; PKC, protein kinase C; PBO, placebo; TCZ, tocilizumab; Th, T helper cell; TLR, toll-like receptors.

Blood composition was also altered in patients with severe vs moderate COVID-19 at baseline by hematological measurements and by inference from bulk RNA-seq. There was a relative decrease in the hematological measurements of total leukocyte fraction (%) of monocytes and lymphocytes and a concomitant increase in neutrophils (**Fig. S4A**). Furthermore, analysis of cell-type–related signatures revealed that patients with severe COVID-19 had lymphopenia, indicated by lower expression of genes associated with T, natural killer (NK), and B cells, and a significant increase in expression of genes associated with mature and immature neutrophils, relative to patients with moderate COVID-19 (**Fig. 3D-E**). Moreover, patients with severe vs moderate COVID-19 had higher expression of an immunosuppressive human leukocyte antigen (HLA)-DR^low^ signature and decreased expression of an HLA-DR^high^-activated classical monocyte subset signature (**Fig. 3D-E**). Several severity-associated cell type signatures (granulocytic myeloid-derived suppressor cells [gMDSC], neutrophils, and HLA-DR^low^ monocytes) were also prognostic for worse clinical outcome by day 28 (**Fig. S4B, Table S3**) even when adjusting for time from symptom onset.

### Elevated expression of antiviral pathway genes in the first 10 days of COVID-19 symptom onset

Since COVID-19 progression can involve time-dependent molecular changes, we sought to understand the temporal patterns of COVID-19 severity and progression-associated biomarkers with respect to time from symptom onset at baseline. We leveraged the fact that the time from symptom onset to baseline measurement varied among patients (median = 11 days [range 1-50]) and compared protein abundance in baseline samples collected from patients sampled ≤10 days from symptom onset (early) to those who were sampled >10 days after symptom onset (late) (**Table 1)**. Results were adjusted for baseline severity since the baseline severity scores of patients sampled late were significantly higher than those sampled earlier (**Fig. S5A**, *P* < 0.05, Fisher’s exact test). Several proteins involved in antiviral response and immune signaling (IFNL1, IFNG, DDX58, RAGE [AGER], CCL8, CXCL10 and IL-12B) displayed higher serum levels in patients sampled early during infection, when adjusting for baseline clinical severity, consistent with an earlier stage of innate immunity (**Fig. S5B-C**). Multiple (6 of 13; DDX58, IL12B, ADH4, GSTA, ITGAI1, SMPDL3A) proteins with greater serum levels in patients sampled early during infection were lower in patients with severe vs moderate COVID-19, but two, RAGE (AGER) and CXCL10 were significantly *higher* in patients with severe vs moderate COVID-19 (Benjamini-Hochberg FDR <0.05, logFC 0.48 and 0.36 for CXCL10 and RAGE, respectively) and higher levels were prognostic for worse clinical outcomes (**Fig. S5B-C, Table S1 and S2**). Most proteins elevated in patients sampled late were also elevated in patients with severe vs moderate COVID-19 (BAIAP2, CLC, SDC1, NEFL, MUC16, FGF23, PRSS2, WFDC2, EDA2R, NMNAT1, SFTPD, PFKFB2, PADI4, CEBPB, TNC, CD177, ACE2, REG3A) and several were also prognostic for worse clinical outcomes (FGF23, EDA2R, EGLN1, TNC, COL6A3, TNFRS10B) (Benjamini-Hochberg FDR <0.05) (**Fig. S5B-C, Table S1 and S2**).

Consistent with the trends observed with proteomics, RNA-seq analysis also showed an increase in expression levels of antiviral response signatures in patients sampled early vs late (**Fig. S5D**). Pathways elevated in patients sampled late, such as heme metabolism and platelet activation, were also elevated in patients with severe COVID-19 (**Fig. S5D**), suggesting these pathways are associated with COVID-19 progression.

### Novel pharmacodynamic biomarkers of tocilizumab treatment identified by serum proteomics

Clinical analysis of the COVACTA trial indicated that tocilizumab treatment improved the time to discharge or ready for discharge of patients hospitalized with COVID-19. To understand the mechanism of action of tocilizumab in these patients, we evaluated tocilizumab’s pharmacodynamic response and its ability to resolve elevated serum protein levels in patients with COVID-19 by comparing whole blood samples collected before treatment and through 60 days after treatment from tocilizumab-treated and placebo-treated patients hospitalized with COVID-19.

Consistent with the known pharmacodynamic effects of tocilizumab and engagement of its target IL-6R, treatment led to initial increased levels of circulating IL-6 and soluble IL-6R, followed by a decrease at later timepoints compared with placebo-treated patients (**Fig. 4A-B**), whereas patients treated with tocilizumab had significantly reduced levels of C-reactive protein (CRP), a downstream target of the IL-6 pathway, at 3 days after treatment vs baseline, whereas CRP did not start to decline in patients who received placebo until 7 days after treatment. This pattern was observed for patients who survived as well as those who died by day 28, suggesting that the changes were a pharmacodynamic effect of tocilizumab treatment, although the levels of CRP did not decrease as substantially in patients who died (**Fig. 4A-B**).

**Fig 4.**
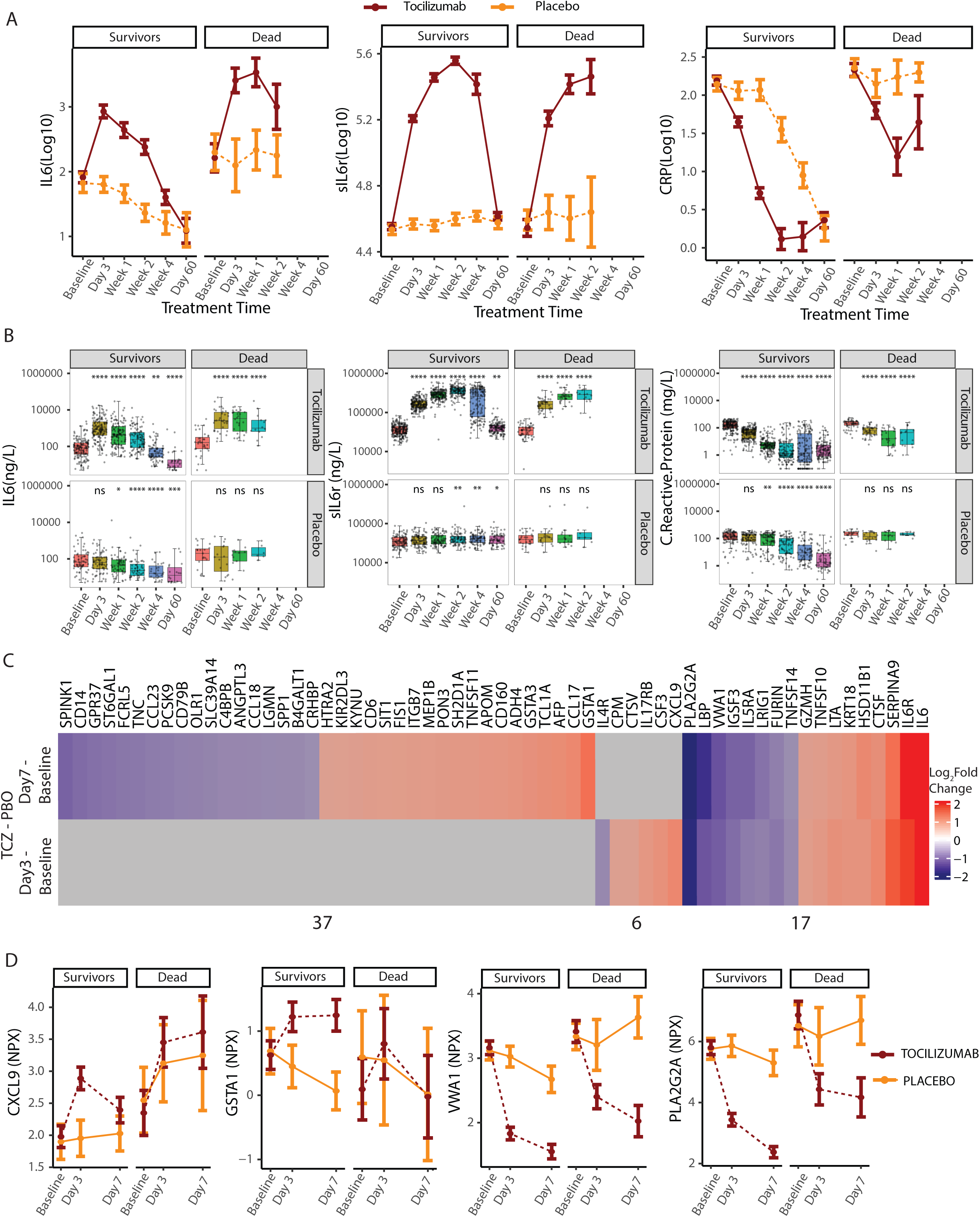
Serum proteins responsive to tocilizumab treatment in hospitalized COVID-19 patients. **(A)** Line plots and **(B)** box plots showing changes in serum levels measured by ELISA of known pharmacodynamic biomarkers of TCZ comparing response to treatment with TCZ and placebo longitudinally for cases showing clinical improvement (survivors) vs those who die by day 28 (dead). Sample numbers plotted are provided in Supplementary Table 5. *Represents statistical significance using Wilcoxon tests at each timepoint using baseline as the reference group. *≤0.05, **≤0.01, ***≤0.001 and ****≤0.0001 **(C)** Heatmap showing proteins differentially regulated between TCZ and placebo treatment by day 3 and day 7 determined by Olink. Proteins highlighted represent those that change on TCZ treatment by at least ±0.5 log_2_fold over placebo. (TCZ Day 3/7 − baseline) – (PBO Day 3/7 − Baseline). Gray cells represent proteins that did not meet the differential cut-off. The indicated numbers represent the number of proteins showing response to TCZ only at day 7 (37), only at day 3 (6) and at both timepoints (17). Proteins colored green represent those that are severity-associated and/or prognostic for worse clinical outcomes and show a significant increase or decrease with tocilizumab treatment relative to placebo. **(D)** Line plots showing examples of treatment response of selected proteins to tocilizumab and placebo faceted by clinical status by day 28. ELISA, enzyme-linked immunosorbent assay; IL-6, interleukin 6; PBO, placebo; TCZ, tocilizumab.

To identify additional pharmacodynamic biomarkers, we compared serum levels of 1472 proteins in blood samples drawn at 3 and 7 days after treatment with tocilizumab or placebo vs baseline. There were 23 and 54 proteins (17 shared) with a significantly greater change (increase or decrease) from baseline in tocilizumab- vs placebo-treated patients at day 3 and 7, respectively (Benjamini-Hochberg FDR < 0.05 using Limma; logFC ≥0.5 or ≤−0.5) (**Fig. 4C-D**). Consistent with our qualified enzyme-linked immunosorbent assay (ELISA) assays, there was also a significant increase in IL-6 and IL-6R after treatment with tocilizumab (**Fig. 4C**). Several of the tocilizumab-responsive proteins showed similar pharmacodynamic trends as IL-6, IL-6R, or CRP at day 7 after treatment (43 proteins by Limma; logFC ≥0.5 or ≤−0.5, no *P* value cut-off, for example, PLA2G2A and VWA1, **Fig. 4D**), whereas tocilizumab had an effect both in patients who survived and those who died by day 28, suggesting these proteins could serve as additional pharmacodynamic biomarkers.

### Tocilizumab-treated patients showed a more rapid normalization of COVID-19 severity-associated protein signatures and blood cellular composition

To assess the basis for the benefit of tocilizumab treatment in patients hospitalized with COVID-19, we next evaluated both serum protein biomarkers and RNA profiles associated with COVID-19 clinical severity for normalization by tocilizumab treatment. Of the proteins associated with severity and that were prognostic for worse clinical outcomes (clinical failure or hospital discharge; adjusted for age, treatment arm, baseline severity, and time from symptom onset), 18 proteins (CD14, GPR37, LBP, OLR1, OPN, PLA2G2A, ST6GAL1, TNC, B4GALT1, CCL23, IL4R, CD79B, IL5RA, FCRL5, SPINK1, FURIN, VWA1 and IGSF3) showed a significantly greater reduction and 7 proteins (IL6, KRT18, CXCL9, CSF3, KYNU, SH2D1A and HTRA2) had a greater increase in response to tocilizumab compared to placebo by day 7 (Benjamini-Hochberg FDR <0.05 using Limma; logFC ≤−0.5 or ≥0.5). By day 28, there were significant reductions (Benjamini-Hochberg FDR <0.05 using Limma; logFC ≤−0.5) in serum levels of most severity-associated proteins and consequent normalization regardless of treatment, indicating clinical recovery for both treatment arms (**Table S1**). Concordant with the few severity-associated proteins normalized on tocilizumab treatment, principal component analysis (PCA) analysis revealed negligible difference in protein profiles from tocilizumab treatment and placebo (**Fig. S6A-B**).

In contrast to the protein profiles, RNA profiles of tocilizumab-treated patients who recovered (survivors) showed faster normalization of a much larger set of blood cell gene expression signatures by day 3 and 7 relative to placebo and patients who died by day 28 (**Fig. 5A-B, Fig. S6C**). Like the serum proteome analysis, there was significant normalization of blood cell gene expression by day 28 regardless of treatment, but at earlier timepoints (days 3 and 7) severity-associated gene expression normalized more rapidly in tocilizumab vs placebo-treated patients (**Fig. 5A-B, Fig. S6C**). Specifically, in patients treated with tocilizumab, 46% (1316 genes) of the 2855 genes initially downregulated in severe COVID-19 vs moderate were significantly upregulated by day 7 after treatment, relative to baseline, and 25% (472 genes) of the 1900 genes initially upregulated in severe COVID-19 vs moderate were significantly downregulated by day 7 after treatment, relative to baseline; this is compared with 1% and 3%, respectively, for placebo-treated patients (Benjamini-Hochberg FDR < 0.05; log2Fold change ≤−0.5 or ≥0.5; **Fig. 5B-C**). Notably, 464 genes that were elevated in severe vs moderate patients at baseline were further elevated following tocilizumab treatment (**Fig. 5C**) and were enriched for pathways such as myeloid cell homeostasis, erythrocyte development, and heme biosynthesis (**Fig. 5D**). These pathways tend to be associated with improved clinical outcomes based on hazard modeling for the end points time to hospital discharge/ready for discharge and time to clinical failure, but the association was not statistically significant (**Fig. 5E**). There was a stronger prognostic signal in the subset of patients with moderate vs severe COVID-19 (**Fig. S7**), suggesting that these pathways might be involved in recovery, especially among patients with moderate severity, that is further promoted by tocilizumab treatment.

**Fig 5.**
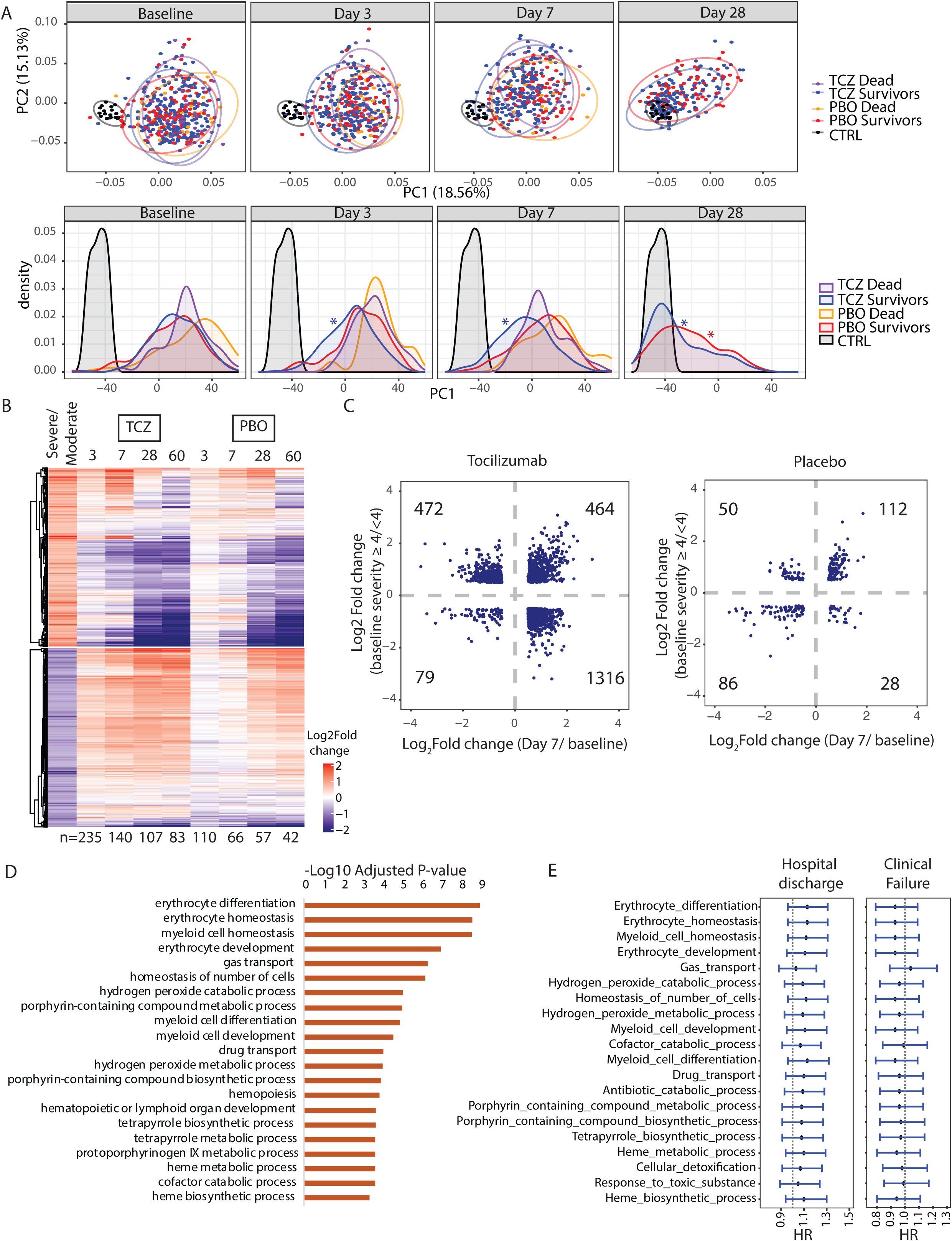
Differential gene expression in response to tocilizumab treatment in hospitalized COVID-19 patients. **(A)** PCA and corresponding density plots showing longitudinal changes in RNA-seq signal at baseline, day 3, day 7, and day 28. \**P* value <0.05, Kolmogorov-Smirnov test performed between the indicated timepoints relative to baseline. **(B)** Heatmap showing longitudinal response of the blood transcriptome to TCZ treatment. Rows represent severity-associated genes identified by DESeq2. For each timepoint only samples measured at baseline and at the indicated timepoint were used for analysis (n indicated below the heatmap). **(C)** Scatter plot of log_2_fold difference between severe vs moderate baseline cases and day 7 after treatment vs baseline for TCZ (left) and Placebo (right). Data points represent genes significantly different by ≥0.5 or ≥-0.5 log2fold for both comparisons. (**D)** Bar plot showing immune pathways enriched among genes expressed higher in severe vs moderate cases at baseline that also show greater increase on TCZ treatment vs placebo by day 7 ([TCZ Day 7 – TCZ Day 1] – [PBO Day 7 – PBO Day 1]). **(E)** Cox proportional hazard modeling for pathways shown in panel D. Forest plots generated as in Fig 3C. CTRL, control; HR, hazard ratio; PCA, principal component analysis; PBO, placebo; TCZ, tocilizumab.

The genes whose expression was reduced at day 3 and 7 following treatment more substantially in tocilizumab- vs placebo-treated patients were enriched for pathways upregulated in severe vs moderate COVID-19 at baseline, such as coagulation and complement. In parallel, genes with stronger upregulation in tocilizumab- vs placebo-treated patients at days 3 and 7 were enriched for pathways involved in lymphocyte biology (**Fig. 6A; Fig. S8A-B**). Conversely, expression of genes involved in antiviral signaling were enriched among genes differentially expressed at day 7 compared to baseline for both treatment arms (**Fig. S8B**).

**Fig 6.**
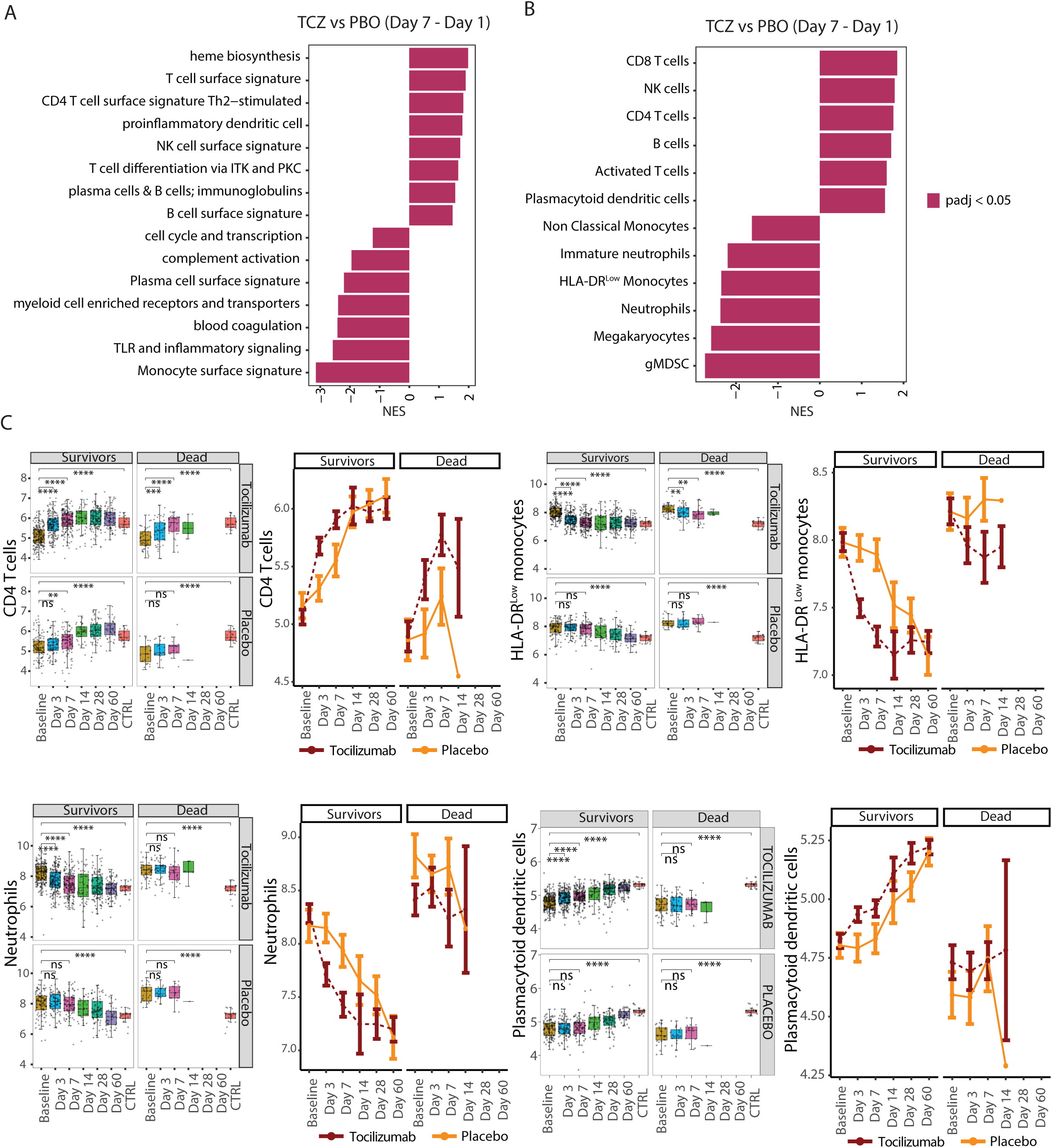
Immune pathways and cellularity signatures responsive to tocilizumab treatment in hospitalized COVID-19 patients. **(A)** Bar plot showing immune pathways enriched among genes showing a greater response to tocilizumab compared to placebo by day 7 (calculated as [TCZ Day 7 – TCZ Day 1] – [PBO Day 7 – PBO Day 1]). **(B)** Bar plot showing blood cell types enriched among genes showing greater response to TCZ compared to placebo by day 7 ([TCZ Day 7 – TCZ Day 1] – [PBO Day 7 – PBO Day 1]). **(C)** Box plot (left) and line plot (right) showing eigengene expression of gene sets corresponding to blood cell types for cases treated with tocilizumab and placebo across timepoints. *Represents statistical significance using pairwise T tests at each timepoint using baseline as reference the reference group. *≤0.05, **≤0.01, ***≤0.001 and ****≤0.0001. Sample numbers corresponding to the plotted categories are provided in Supplementary Table 5. CD, cluster of differentiation; gMDSC, granulocytic myeloid-derived suppressor cells; HLA, human leukocyte antigen; ITK, IL-2 inducible T cell kinase; IL-2, interleukin 2; NES, normalized enrichment score; NK, natural killer; PKC, protein kinase C; PBO, placebo; TCZ, tocilizumab; Th, T helper cell; TLR, toll-like receptors.

Hematology data indicated that tocilizumab treatment led to more rapid to changes in blood cell composition compared to placebo, rapidly normalizing hematologic blood cellularity, with lymphocytes and neutrophil frequency showing a greater change in response to tocilizumab compared with placebo starting at day 3 (**Fig. S9**). Similarly, there was a more rapid reduction of expression of gene signatures of megakaryocytes and immunosuppressive myeloid subsets, including HLA-DR^low^ monocytes and gMDSC, in response to tocilizumab vs placebo (**Fig. 6B-C; Fig. S10A-B**). Despite a rapid reduction in blood neutrophil numbers by tocilizumab, there were differential effects on expression signatures of neutrophil cell states. Consistent with neutrophil numbers, the expression of genes associated with mature neutrophils was reduced more rapidly following tocilizumab vs placebo treatment (**Fig. 6B-C; Fig. S10A-B**). Conversely, placebo-treated patients showed a significant increase in expression of immature neutrophil gene signatures at days 3 and 7, which was tempered in tocilizumab-treated patients (**Fig. 6B-C; Fig. S10A-B**). Gene signatures associated with cluster of differentiation 4 (CD4) T-cells, cluster of differentiation 8 (CD8) T-cells, NK cells, HLA-DR^low^ monocytes, gMDSCs and megakaryocytes were significantly responsive to tocilizumab both in patients showing clinical recovery and those progressing to death by day 28, suggesting an effect of tocilizumab treatment (**Fig. 6C and S10C-D**). Despite being responsive to tocilizumab treatment, patients who died by day 28 showed less resolution in gene expression of these gene sets at day 7 compared to survivors (**Fig. 6C and S10C-D**). In contrast, neutrophils, plasmacytoid dendritic cells, B cells, and nonclassical monocytes showed a significantly faster response to tocilizumab in patients who survived compared to those who died (**Fig. 6C and S10C-D**).

### Corticosteroid treatment dampens inflammatory response in COVID-19 patients

Meta-analysis of multiple clinical trials assessing tocilizumab efficacy indicates that tocilizumab has greater benefit for patients receiving concomitant corticosteroids than those not receiving corticosteroids. Based on this analysis, current guidelines recommend treatment with tocilizumab combined with corticosteroids for patients with severe COVID-19 (*2*). The COVACTA study enrolled patients before corticosteroids became the standard of care for treatment of COVID-19; therefore, only a low proportion of patients were on corticosteroids at baseline (eg, 19.2% in the TCZ arm; 27.0% in the placebo arm for the patients in the Olink proteomic analysis) (**Table 1**). Patients on corticosteroids at baseline had significantly lower serum levels of several cytokines, including IL-6, IL-12B, IFNG, and IL-1RN, and lower expression levels of genes associated with antiviral and inflammatory pathways at baseline compared to patients with no corticosteroid treatment (**Fig. S11**). The small number of patients treated with corticosteroids precludes further analysis of the combined effects of tocilizumab plus corticosteroids.

## Discussion

Tocilizumab, an IL-6R inhibitor, is approved globally for the treatment of COVID-19 in hospitalized adult patients who are receiving systemic corticosteroids and require supplemental oxygen, noninvasive or invasive mechanical ventilation, or extracorporeal membrane oxygenation (ECMO), yet the effects of IL-6R blockade on the pathogenesis of COVID-19 remains unclear. Our study identifies cellular and molecular pathways associated with COVID-19 severity modified by tocilizumab and provides additional mechanistic insight on the role of IL-6 signaling in COVID-19. Overall, tocilizumab treatment resulted in faster reversal of COVID-19-associated dysfunction in the myeloid and lymphoid compartments as well as faster downregulation of IL-6-associated inflammatory mediators and biomarkers compared with placebo treatment. This is consistent with reduced time to hospital discharge/ready for discharge as observed in the COVACTA study and the WHO meta-analysis in a larger number of patients that showed reduced mortality and faster recovery in patients treated with tocilizumab.

We corroborated previously identified biologic abnormalities in hospitalized patients with COVID-19, including excessive inflammatory cytokine production and dysregulated immune cell activity characterized by lymphopenia, high levels of immature neutrophils and low HLA-DR expressing monocytes, and disturbances in other biological pathways, such as coagulation and platelet activation, which together can cause multiorgan injury (*21, 24*). Using selected organ-specific markers from Olink proteomics data, we support published findings (*21*) that show evidence of injury of the lung, heart, and liver in patients with severe COVID-19. By contrast, the gut-specific markers GPA33, FABP2, and CDH17 were found to be less abundant in COVID-19 patients. This is consistent with a recent finding showing lower serum levels of I-FABP to be associated with COVID-19 severity (*25*). It is yet unclear whether and how lower serum levels of these markers relate to manifestations of COVID-19 on the gastrointestinal system. Taken together, these findings confirm that the patient population enrolled in COVACTA reflects observational cohorts of patients with COVID-19, despite potential confounders (eg, inclusion/exclusion criteria).

Analysis of proteomic and transcriptomic data indicate that inflammation, lymphopenia, and myeloid dysregulation resolved more rapidly following treatment with tocilizumab compared with placebo, which is consistent with faster time to hospital discharge of tocilizumab-treated patients in this study. Most transcriptional changes associated with disease severity were responsive to tocilizumab within a week of treatment, with more rapid reduction of gene sets for megakaryocytes, immunosuppressive myeloid subsets, and mature neutrophils, and increased expression of immature neutrophils compared to placebo. Tocilizumab also led to consistently faster normalization of blood counts, including reductions of neutrophils and increases in lymphocytes, in patients who showed clinical improvement within 28 days of treatment. Notably, a subset of genes induced by COVID-19 were further increased with tocilizumab treatment; however, increased expression was associated with improved clinical outcomes. Functional enrichment analysis of these genes implicated pathways involved in erythrocyte development and heme biosynthesis, suggesting that tocilizumab could have an indirect effect in reversing anemia associated with COVID-19, leading to improved clinical outcome (*26*).

Despite these observations, the COVACTA study did not meet its primary end point of improved clinical status on a 7-category ordinal scale at day 28, and no difference was observed in mortality at day 28 with tocilizumab versus placebo. Pharmacodynamic effects of tocilizumab on the biomarkers CRP, IL-6, and IL-6R were observed in patients regardless of their clinical progression by day 28, although the levels of CRP in patients who died decreased to a lesser extent and appeared to rebound in some patients. These findings could indicate insufficient target engagement or faster drug clearance in patients who died, or the late increase in CRP could reflect activation of IL-6 pathways by a new trigger that might have contributed to mortality (eg, secondary infection). Similarly, gene signatures for neutrophils, plasmacytoid dendritic cells, B cells and nonclassical monocytes that normalized more rapidly in patients treated with tocilizumab compared to placebo failed to normalize in patients who died by day 28 irrespective of tocilizumab or placebo treatment. Neutrophil gene signatures track temporally with respiratory dysfunction and disease resolution in patients with COVID-19 (*27*), suggesting resolution of neutrophilic inflammation is important for disease recovery. Tocilizumab treatment did not appear to alter antiviral or interferon pathways, consistent with similar viral load decline in patients who received tocilizumab or placebo treatment as previously reported (*28*). In addition, tocilizumab did not accelerate reversal of multiple biomarkers associated with tissue damage in this study, although analysis of alternate indicators might be necessary to evaluate recovery from organ injury. In totality, these findings suggest that therapies that further dampen neutrophilic inflammation, bolster early antiviral responses, reduce viral load, or promote tissue repair may be complementary to tocilizumab for the treatment of severe COVID-19. It is, however, notable that tocilizumab dosed in combination with the antiviral remdesivir did not show additional benefit compared with placebo plus remdesivir in the REMDACTA study (*16*).

Meta-analysis indicated larger benefits of tocilizumab in patients who also received corticosteroids (*2*). At the time of COVACTA study enrollment, corticosteroids were not standard of care for patients hospitalized with COVID-19. Therefore, in COVACTA relatively few patients received corticosteroid treatment, the treatment was not randomized, and the doses and timing for treatment varied considerably, which precludes analysis of tocilizumab’s mechanism of action in combination with steroids. In the baseline samples from the approximately 20% of COVACTA patients who received corticosteroids at baseline, corticosteroid treatment was associated with effects on several cytokines associated with inflammatory and antiviral pathways. The combined anti-inflammatory effects of steroids (eg, on neutrophilic inflammation) with tocilizumab may yield faster and stronger resolution of immune dysregulation.

Temporal dynamics of the host immune response to SARS-CoV-2 is a key component of COVID-19 pathogenesis and the timing for clinical intervention. We found that patients who enrolled earlier (<10 days) after initial symptom onset tended to have less severe COVID-19 and elevated baseline expression of proteins and genes related to antiviral, interferon, and innate immune pathways, consistent with the earlier stages of immune response to viruses, whereas patients who enrolled later, after symptom onset (≥ 10 days), tended to have more severe COVID-19 and higher levels of proteins reflecting diverse biologic mechanisms that were also elevated in patients with severe vs moderate disease. The notable exceptions were CXCL10 and AGER (RAGE), which are biomarkers elevated in early patients that were also elevated at baseline in patients with severe vs moderate COVID-19 and prognostic for clinical failure and longer time to hospital discharge/ready for discharge. These findings are consistent with other recent reports identifying CXCL10 and AGER as strong prognostic biomarkers for mortality and ICU admission and showing biomarkers of alveolar injury, including AGER, are elevated early in COVID-19 disease course (*29–35*). Despite differences in time from symptom onset that could reflect different stages of COVID-19, no difference in tocilizumab efficacy was observed in patients further grouped by timing of initial symptom onset in the COVACTA study (*28*).

The main strength of this study is the rigorous transcriptomic and proteomic analysis used to determine the effect of tocilizumab in patients hospitalized with severe COVID-19 in a randomized, double-blind, placebo-controlled trial that also involved longitudinal sampling. This study was conducted before widespread use of efficacious antivirals and corticosteroids, and thereby allows evaluation of tocilizumab’s mechanism of action without confounding effects of other treatments. This analysis has several limitations. First, the organ-specific protein sets for analysis of organ damage were generated using tissue-specific transcriptome data from healthy individuals provided by the GTEX consortium. Second, most data were derived from whole blood and serum samples of patients, which may reflect changes in cellularity rather than alterations of cell-intrinsic pathways and may not accurately represent the tissue specific pathogenesis of COVID-19. Third, although meta-analysis of multiple studies supports the efficacy of tocilizumab for treatment of COVID-19, the COVACTA study did not meet its primary end point of improved clinical status on day 28, although some clinical benefits (such as shortened time to hospital discharge/ready for discharge) were associated with tocilizumab treatment (*15*). Therefore, the ability to associate changes in levels of biomarkers identified here to clinical efficacy or identify baseline biomarkers predictive of clinical response to tocilizumab is limited. Fourth, in contrast to RNA-seq, where hundreds of genes showed differential response to treatment with tocilizumab, relatively few proteins showed a significant difference in response to tocilizumab based on Olink. This discrepancy could be attributable to the differences in coverage (1472 proteins in Olink vs the entire transcriptome for RNA-seq) and sensitivity to detect and measure changes. Thus, in order to reveal trends in the serum protein response to tocilizumab treatment, the thresholds were made less stringent by using an unadjusted *P* value cut-off.

In summary, the present study identified prognostic biomarkers for COVID-19 disease progression, confirmed the pharmacologic activity and identified novel pharmacodynamic biomarkers for tocilizumab in patients with COVID-19, uncovered the potential mechanism of action of tocilizumab in patients hospitalized with COVID-19, and revealed pathways elevated in patients who did not recover that may be targets for new COVID-19 therapeutics to complement anti-inflammatory treatments like tocilizumab. These results suggest that more rapid resolution of inflammation, myeloid dysregulation, and lymphopenia in response to tocilizumab treatment may potentially lead to faster recovery, as demonstrated by the shorter time to hospital discharge reported in the COVACTA trial and supported by the WHO meta-analysis of a larger number of patients (*2*). These findings advance understanding of the pathogenesis of viral pneumonia and ARDS and the potential benefit of anti-inflammatory treatments including IL-6R blockade for these diseases.

## Materials and Methods

### Study design

Details of COVACTA have been previously reported (ClinicalTrials.gov, NCT04320615) (*15*). Briefly, COVACTA was a randomized, placebo-controlled, double-blind, global, multicenter, phase 3 trial investigating the efficacy and safety of tocilizumab versus placebo in patients hospitalized with severe COVID-19 pneumonia after receiving standard care according to local practice. Key inclusion criteria were hospitalization with SARS-CoV-2 infection confirmed by polymerase chain reaction of any specimen and blood oxygen saturation (SpO_2_) ≤93% or partial pressure of oxygen/fraction of inspired oxygen (PaO_2_/FiO_2_) <300 mm/Hg despite receiving local standard care (could include antivirals or low-dose steroids). Key exclusion criteria were active or suspected infection (other than SARS-CoV-2); alanine aminotransferase or aspartate aminotransferase >10× the upper limit of normal; absolute neutrophil count <1000/mL; and platelet count <50,000/mL. Eligible adult patients were stratified by geographic region (North America/Europe) and MV (yes/no) and randomly assigned 2:1 to receive a single intravenous tocilizumab infusion (8 mg/kg body weight) or placebo in addition to standard care. Patients who showed disease worsening or no clinical improvement were allowed to receive a second tocilizumab/placebo infusion 8 to 24 hours after the first dose. The primary end point (improved clinical status) was assessed at baseline and after dose throughout the study using a 7-category ordinal scale with increasing severity from 1 to 7 as follows: 1, discharged or ready for discharge; 2, non-ICU hospital ward, not requiring supplemental oxygen; 3, non-ICU hospital ward, requiring supplemental oxygen; 4, ICU or non-ICU hospital ward, requiring noninvasive ventilation or high-flow oxygen; 5, ICU, requiring intubation and mechanical ventilation; 6, ICU, requiring extracorporeal membrane oxygenation or mechanical ventilation and additional organ support; 7, and death (*14*). At baseline, “Moderate” COVID-19 was defined as an ordinal scale score <4 and “severe” COVID-19 was defined as ordinal scale score ≥4. Secondary end points included mortality at day 28, incidence of MV (in patients not receiving MV at randomization), and time to hospital discharge. The study was conducted in accordance with the International Council for Harmonization E6 guideline for good clinical practice and the Declaration of Helsinki or local regulations, whichever afforded greater patient protection. The protocol was reviewed and approved by all appropriate institutional review boards and ethics committees. Informed consent was obtained for all enrolled patients.

### Transcriptomic and proteomic assessments

Serum IL-6R levels were measured using an immunoassay method validated at QPS (Quantikine ELISA; R&D Systems Minneapolis, MN). Serum IL-6 and CRP levels were measured using in vitro diagnostic methods validated at PPD (Roche Cobas; Roche Diagnostics, Indianapolis, IN).

Serum proteomics were assessed for 437 patients from COVACTA and the 16 healthy individuals (control, run as quadruplets) using the Olink Explore 1536 platform to measure 1472 proteins and 48 control assays dividing into four 384-plex panels constituting inflammation, oncology, cardiometabolic, and neurology proteins (Olink, Uppsala, Sweden). Samples were treated with 1% Triton X-100 for virus inactivation followed by random allocation onto 96-well plates based on treatment, ordinal scale at time of randomization and at day 28, sex, geographic location, MV status, and age. All visits for each patient were measured on the same plate.

Transcriptomic assessment was performed on RNA isolated from blood PaxGene (Qiagen, Hilden, Germany) samples using the central laboratory Q2 Solutions. 1.25 µg of RNA was used for sequencing libraries prepared with TruSeq^®^ Stranded mRNA Library Prep (Illumina, San Diego, CA, USA). Samples were randomized based on clinical parameters of patients (age, sex, baseline severity, geographic location, MV status, baseline antiviral treatment, and the day of sample collection) and sequenced by Illumina RNA-seq by 50 base-pair single-end reads at a read depth of 50 million reads per sample.

### RNA-seq analysis

RNA-seq reads were mapped to the human reference genome (Version 38) using gsnap (*36*). Trimmed reads mapping to ribosomal genes were filtered out before downstream analysis. Read counts were generated for the Gencode v27 annotation using the summarize Overlaps method from bioC in mode “IntersectionStrict” (*36, 37*). Differential expression analysis was performed as described in the DESeq2 vignette (*38*). To determine severe COVID-19-associated dysregulation, baseline (pretreated) samples were divided into moderate (baseline score <4) and severe (baseline score ≥4) groups according to severity scores. For identification of severity-associated genes, differential expression analysis was performed by adding age as a covariate to the model. For analysis of prognostic association, differential expression analysis was performed between baseline samples from patients progressing to better and worse clinical outcomes based on mortality. The baseline severity score and age were added to this model as covariates. Longitudinal treatment response was assessed by comparing gene expression at baseline and at day 3, 7, or 28 using patients with measurements available at both timepoints used for the analysis. Differential treatment effects on gene expression were determined using the likelihood ratio test as described in the DESeq2 vignette. No covariates were added to the models used for longitudinal gene expression analysis.

Gene set enrichment analyses were performed on unfiltered DESeq2 outputs ranked by log2fold change using the FGSEA Bioconductor package (*38, 39*). Enrichment of immune pathways were calculated using gene sets generated from the whole-blood transcriptome modules (*40*). Gene sets associated with different blood cell types were defined using scRNA-seq analysis outputs (see Table S3 from reference (*24*)) generated from the cohort of healthy individuals and patients with mild or severe COVID-19. Gene sets specific to blood cell types were defined by extracting the top 20 genes from the cluster assignments ranked in decreasing order of “avg LogFC” and “pct.1”, and increasing order of “Pct.2” from “Whole blood + peripheral blood mononuclear cells (PBMC)” dataset (see Table S4 and Fig. 7A from reference (*24*)) and “PBMC only” dataset (see Table S4 and Fig. 2D from reference (*24*)). The quality of the gene sets was evaluated by checking for concordance between the gene sets derived from the “Whole blood + PBMC” and “PBMC only” datasets **(Fig. S12).** Since the gene sets for nonclassical monocytes failed to show concordance between the 2 source datasets, the dataset showing better correlation with a published nonclassical monocyte gene set was picked (*41*). The mature neutrophil gene set was derived by intersecting the gene sets derived from “Whole blood + PBMC’’ and “PBMC only” datasets and evaluated by correlating with a published mature neutrophil gene set (*42*) neutrophil frequency in the COVACTA patients measured at baseline **(Fig. S12C).** HLA-DR^low^ and HLA-DR^high^ specific gene sets were defined using the PBMC-only scRNA-seq data (see Fig. 2D from reference (*24*)) filtered for genes enriched in clusters 1-3 by at least avg logFC ≥1 and those that lack enrichment in the neutrophil specific gene sets. Cell type specific gene sets are provided in **Supplemental Table 4**.

Eigengene expression of the specified gene sets were calculated as described using the GSDecon package (https://github.com/JasonHackney/GSDecon/https://github.com/JasonHackney/GSDecon/https://github.com/JasonHackney/GSDecon/). Briefly, DESeq2 normalized counts associated with the individual gene sets were standardized to a mean value of 0 and standard deviation of 1, followed by singular value decomposition of the matrix. The resulting matrix was then adjusted to retain only the first singular value. The normalized count matrix was regenerated using the adjusted singular value decomposition diagonal matrix and eigengene values were derived as the mean expression per sample of the supplied genes.

Heatmaps were generated using ComplexHeatmap (*43*). Volcano plots were generated using EnhancedVolcano (https://github.com/kevinblighe/EnhancedVolcanohttps://github.com/kevinblighe/EnhancedVolcanohttps://github.com/kevinblighe/EnhancedVolcano). Box plots were generated using ggplot2 (https://github.com/tidyverse/ggplot2) and the KEGG pathway diagrams were generated using the Pathview package (*43, 44*). Gene ontology analysis shown in **Fig. S5D** were performed using the GOnet webtool (*43–45*).

### Olink proteomics analysis

Protein abundance measurements (normalized protein expression, NPX) were determined by Olink using their standard pipeline. The quality of Olink data were assessed and outlier samples were filtered out prior to downstream analysis (**Fig. S13**). The quality of these measurements was evaluated at the sample and assay level based on Olink’s internal QC flag and the limit of detection (LOD) of the individual assays. The quality of individual assays was assessed by flagging assays for which <25% of samples pass Olink’s internal QC and have measurements above the assay LOD. Although these low-quality assays were flagged, they were not filtered out from downstream analysis. Outlier samples were identified based on their interquartile range and PCA analysis, which were filtered out prior to downstream analysis. Additional samples were filtered out if the fraction of assays that pass Olink’s internal QC and have measurements above their LOD is <25%. 388 samples remained (tocilizumab, *n* = 266; placebo, *n* = 122) for downstream analysis after filtering out low-quality samples. Differential protein abundance analysis was performed on all 1472 proteins included in the Olink explore 1536 using the limma Bioconductor package. The baseline, prognostic, and longitudinal analyses were performed as described for RNA-seq. The generated limma outputs for each assay were mapped to their official gene symbols prior to plotting. For proteins with repeat measurements across Olink panels (IL6, CXCL8 and TNF), the assays showing the highest standard deviation were plotted.

### PCA analysis

PCA analysis of RNA-seq data were performed using the following steps: (1) count data were DESeq2 normalized and transformed using variance stabilizing transformation; (2) data was filtered for the first 2000 most variable genes (standard deviation); (3) PCA was performed and plotted using the R package ‘ggfortify’; (4) density and cumulative distribution plots were then plotted using PC1 scores; (5) differences in the distributions of sample subgroups highlighted in the figures were tested using the Kolmogorov-Smirnov test. PCA analysis of Olink data were performed in the same way except all protein data were used for the analysis followed by the steps 3-5.

### Defining organ-specific protein signatures

Tissue-specific gene expression of proteins represented in the Olink explore 1536 platform were determined using healthy tissue-derived RNA-seq data (GTEX, downloaded from https://www.proteinatlas.org/about/download). Tissue-specific expression values were summarized at the organ level by assigning the maximum transcript per million value to the available organs for each of the 1472 genes. A protein was considered to be enriched in an organ if its gene was expressed the highest in that organ and was at least 4-fold higher than the next highly expressed gene. Organ-specific protein sets were further refined to include those that were positively correlated to each other based on their protein expression (NPX) in our study. In addition, proteins annotated to be blood secreted (Human Protein atlas) were filtered out. To associate the liver-specific proteins to liver injury, we calculated Spearman correlation between their protein levels (NPX) and ALT/AST values across all baseline samples in the COVACTA study.

### Statistical analysis

Analysis was performed in R version 4.1.0. Statistical analysis of Olink and RNA-seq data were performed using Limma and DESeq2 R packages, respectively. Individual protein biomarker data from ELISA and comparisons of cell frequencies were tested using Wilcoxon test. Statistical comparisons of PC1 distributions for Olink and RNA-seq data were performed using Kolmogorov-Smirnov test. A Cox proportional hazard model was fit using the Survival package (https://github.com/therneau/survival). The model included expression of a gene set (eigengene value) or protein abundance (NPX) with additional covariates, age, treatment arm and baseline severity. The gene/protein expression values and age were included as continuous variables; treatment arm and baseline severity were coded as factors and ordinals, respectively. Hazard ratios and 95% confidence interval estimates were calculated, and forest plots were generated using ggplot2. *P* values for multiple hypothesis testing were adjusted using the Benjamini-Hochberg procedure.

## Acknowledgements

We thank Anna Hupalowska for her preparation of Figure 1. We also thank Bongin Yoo for thoughtful review of the manuscript. Editorial and administrative assistance was provided by ApotheCom (San Francisco, CA) and was funded by F. Hoffmann-La Roche Ltd.

## Funding

F. Hoffmann-La Roche Ltd and the US Department of Health and Human Services, Office of the Assistant Secretary for Preparedness and Response, Biomedical Advanced Research and Development Authority, under OT number HHSO100201800036C.

## Author contributions

HS: Conceptualization, Methodology, Software, Formal Analysis, Data Curation, Writing – Original Draft Preparation, Writing – Review & Editing

JAH: Resources, Writing – Review & Editing, Supervision

CMR: Conceptualization, Writing – Original Draft Preparation, Writing – Review & Editing

AT: Formal Analysis, Data Curation, Writing – Review & Editing

AQ: Data Curation

OO: Writing – Original Draft Preparation, Writing – Review & Editing

JM: Conceptualization, Resources, Writing – Original Draft Preparation, Writing – Review & Editing, Visualization, Supervision, Project Administration

FC: Conceptualization, Methodology, Investigation, Writing – Review & Editing

MB: Conceptualization, Methodology, Investigation, Writing – Review & Editing

LT: Conceptualization, Methodology, Investigation, Writing – Review & Editing, Funding Acquisition

AR: Formal analysis, Writing – Review & Editing, Supervision

IOR: Conceptualization, Writing – Review & Editing, Visualization

RNB: Conceptualization, Methodology, Investigation, Resources, Writing – Original Draft Preparation, Writing – Review & Editing, Supervision, Project Administration

## Declaration of interest

HS, JAH, CMR, AQ, OO, JM, FC, MB, LT, AR, and RNB are employees of Roche/Genentech and hold stock and/or stock options in Roche/Genentech.

AT is an employee of Roche/Genentech.

FC has a patent pending to Genentech for biomarkers for predicting a response to an interleukin (IL)-6 antagonist (P36367-US).

LT is an author of a patent Method for Treating Pneumonia, including COVID-19 Pneumonia with an IL-6 Antagonist pending, owned by Genentech/Roche.

AR is a co-founder and equity holder of Celsius Therapeutics, an equity holder in Immunitas Therapeutics and, until July 31, 2020, was a scientific advisory board member of ThermoFisher Scientific, Syros Pharmaceuticals, Asimov, and Neogene Therapeutics. AR is a named inventor on multiple patents related to single cell and spatial genomics filed by or issued to the Broad Institute.

IOR has nothing to declare.

## Data materials and availability

Qualified researchers may request access to RNA-seq, proteomics, and clinical metadata for the analyses presented here through the European Genome-Phenome Archive (EGA; accession number EGAS00001006688; https://ega-archive.org/) upon publication. Individual patient level clinical data for the COVACTA study is available through the clinical study data request platform (https://vivli.org/). Further details on Roche’s criteria for eligible studies are available here (https://vivli.org/members/ourmembers/). For further details on Roche’s Global Policy on the Sharing of Clinical Information and how to request access to related clinical study documents, see here (https://www.roche.com/research_and_development/who_we_are_how_we_work/clinical_trials/our_commitment_to_data_sharing.htm).

## Supplementary materials

**Supplemental Table 1:** Log_2_fold change outputs from longitudinal Limma analysis for severity-associated proteins.

**Supplemental Table 2:** Hazard modeling of serum proteomic data for length of hospital stay and clinical failure.

**Supplemental Table 3:** Hazard modeling of gene expression signatures for length of hospital stay and clinical failure.

**Supplemental Table 4:** Cell type specific gene sets used for RNA-seq analysis

**Supplemental Table 5:** Sample counts per category corresponding to box and line plots.

**Fig S1.** Severity-associated changes in serum levels of organ-specific proteins.

**Fig S2.** Association of liver-specific proteins to clinical biomarkers indicative of liver dysfunction.

**Fig S3.** Proteins prognostic for higher mortality and clinical failure.

**Fig S4.** Changes in blood cell counts and biological pathways in severe COVID-19 cases.

**Fig S5.** Time to symptom onset-dependent changes in protein and transcript levels at baseline.

**Fig S6.** PCA analysis evaluating the benefit of tocilizumab treatment.

**Fig S7.** Tocilizumab treatment leads to upregulation of genes prognostic for better clinical outcomes.

**Fig S8.** Effect of tocilizumab treatment on immune pathways.

**Fig S9.** Effect of tocilizumab treatment on blood cell counts.

**Fig S10.** Effect of tocilizumab treatment on cell type signatures.

**Fig S11.** Effect of corticosteroid treatment on blood transcript and serum protein levels at baseline.

**Fig S12.** Quality assessment of blood cell type gene sets.

**Fig S13.** Quality assessment of Olink data.

## Notes

### Summary of Updates

Revisions to figures 4, 6, and S10. Text updates to the introduction, results and discussion section to clarify.

