## Supplemental Figs + Tables (Fig S1-S13 and tables 1-5) for "Tocilizumab treatment leads to early resolution of lymphopenia and myeloid dysregulation in patients hospitalized with COVID-19"

**Fig S1. Severity-associated changes in serum levels of organ-specific proteins.** Boxplots showing changes in serum levels across baseline severity scores for **(A)** lung-, **(B)** heart-/skeletal muscle-, **(C)** pancreas-, and **(D)** liver-specific proteins. Number of patient samples analyzed is indicated in the table **(E).**

CTRL, control; NPX, normalized protein expression.

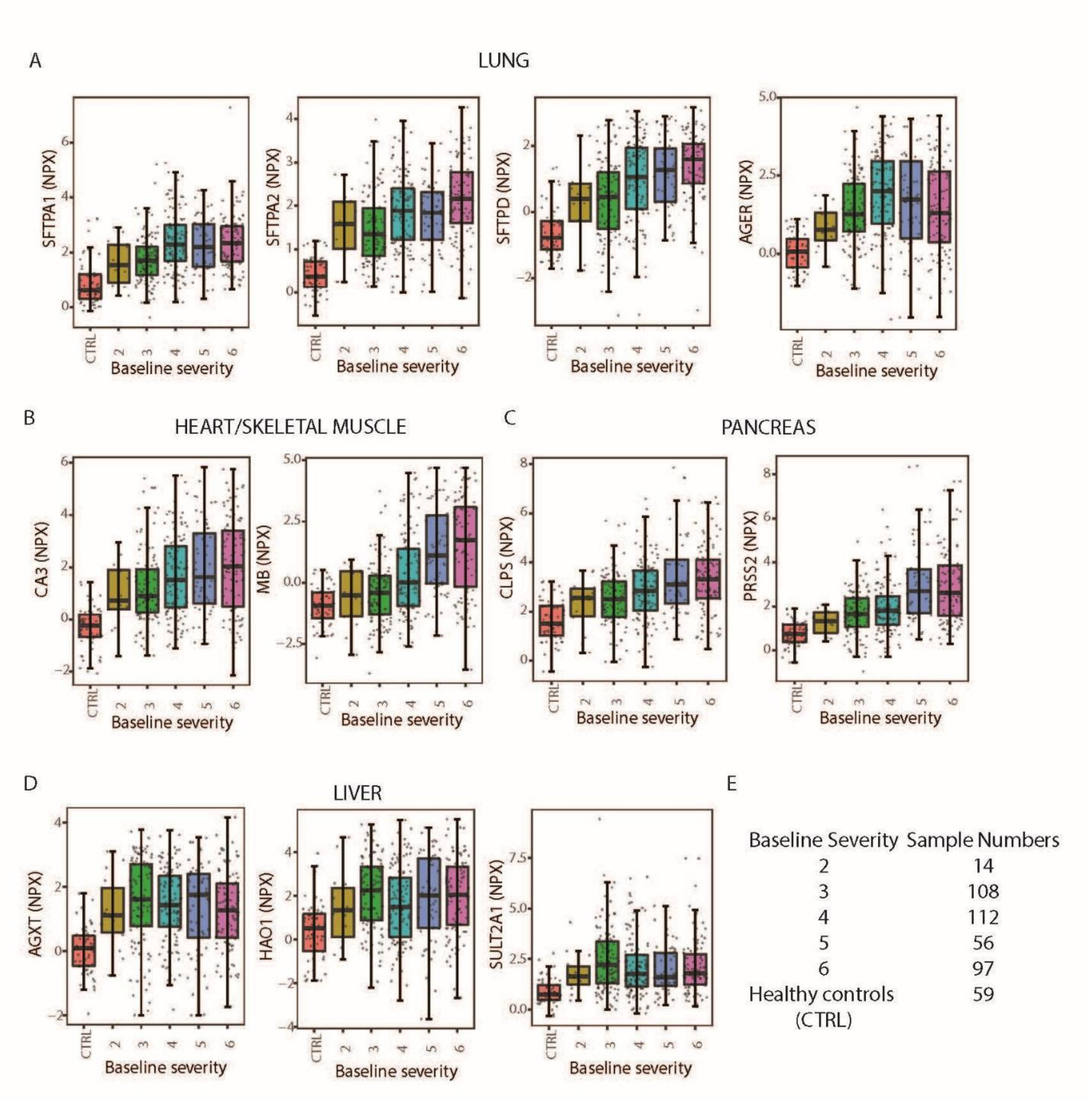

**Fig S2. Association of liver-specific proteins to clinical biomarkers indicative of liver dysfunction.** Scatter plot comparing Olink measurements of liver-specific proteins (X-axis) to liver function test results (Y-axis) – alanine transaminase (top) and aspartate transaminase (bottom).

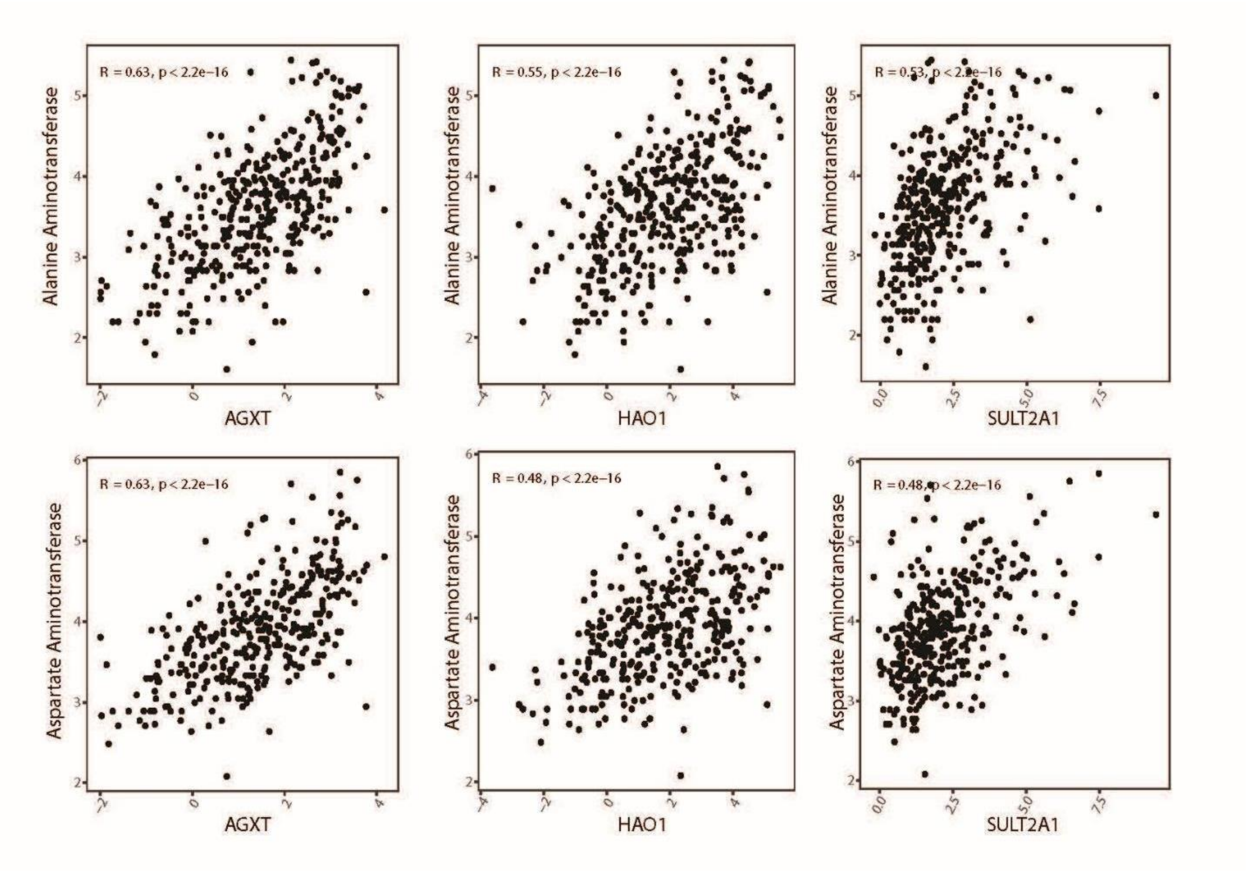

**Fig S3. Proteins prognostic for higher mortality and clinical failure. (A)** Volcano plots showing proteins differentially regulated in severe COVID-19 as determined by Olink. Proteins highlighted represent those that are significantly upregulated by at least +0.5 $log_2$fold in nonsurvivors by day 28. **(B)** Proteins prognostic for worse clinical outcome (time to clinical failure) identified using Cox proportional hazard model. Only proteins showing statistically significant prognostic association are shown (Benjamini-Hochberg adjusted *P* < 0.05, represented by ***).

HR, hazard ratio.

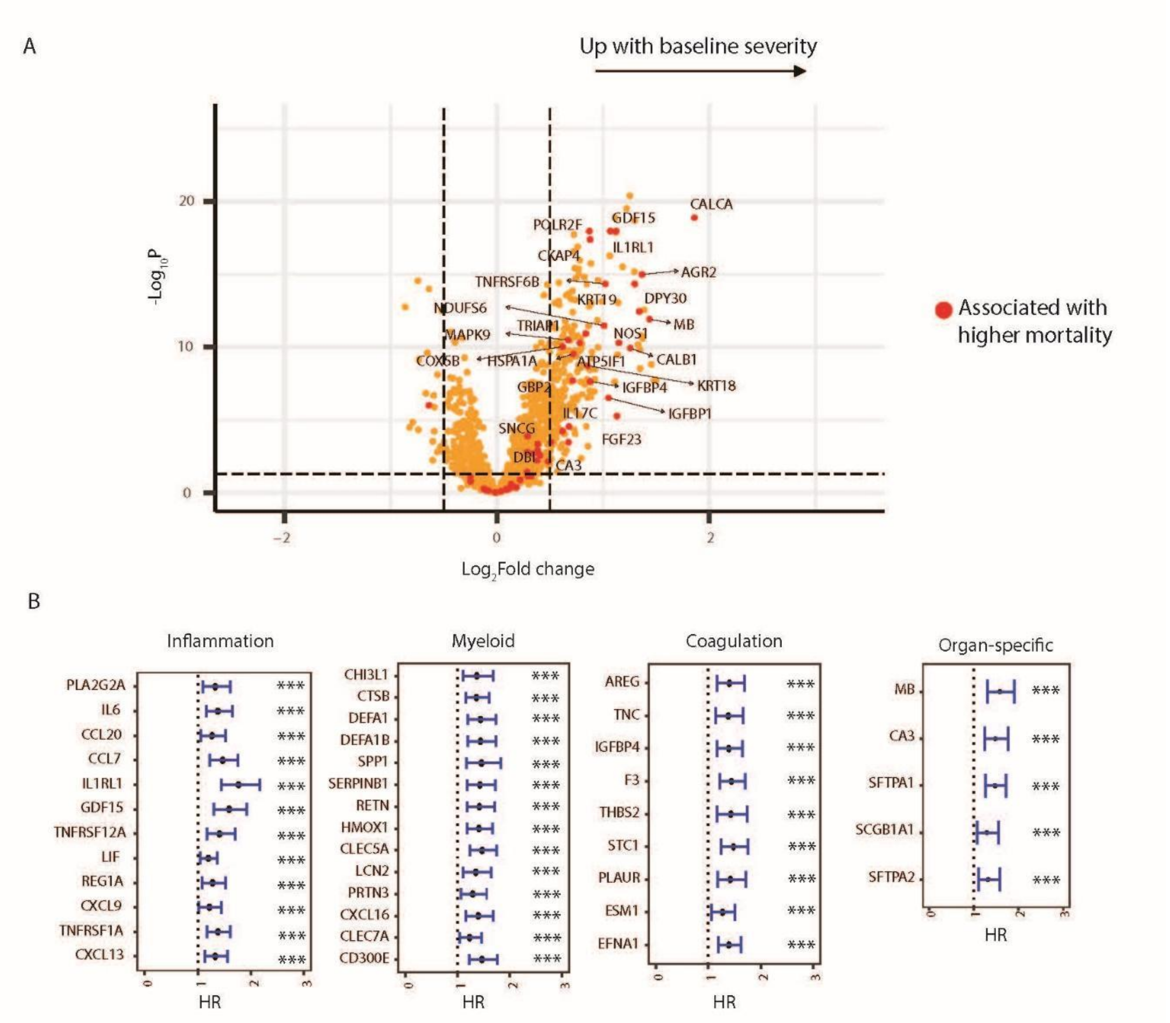

**Fig S4. Changes in blood cell counts and biological pathways in severe COVID-19 cases.** **(A)** Boxplots showing changes of blood cell counts with baseline severity. Percentages were calculated as ratio of specified cell type to leukocyte counts. The normal range upper limit of the absolute counts can range from 2.4-5.2 $10^9$/L for lymphocytes, 0.36-1.5 $10^9$/L for monocytes, and 4.8-8.89 $10^9$/L for neutrophils. The normal range lower limit can range from 0.5-1.5 $10^9$/L for lymphocytes, 0-0.5 $10^9$/L for monocytes, and 0.95-3.15 $10^9$/L for neutrophils. **(B)** Fast gene set enrichment analysis of differentially expressed genes between cases associated with higher mortality (death by day 28, n = 83) and survivors (n = 321). Higher NES represents upregulation of the indicated immune pathway. (**C)** Forest plot showing HRs with 95% confidence intervals identified using Cox proportional hazard model depicting recovery (left) and clinical failure (right) from COVID-19 for cell types shown in Figure 3D. ***Represents Benjamini-Hochberg adjusted $P < 0.05$.

Abs, absolute; CD, cluster of differentiation; gMDSC; granulocytic myeloid-derived suppressor cells; HLA, human leukocyte antigen; HR, hazard ratio; NES, normalized enrichment score; NK, natural killer; NS, not significant; Pct, percent; pval, *P* value; padj, adjusted *P* value.

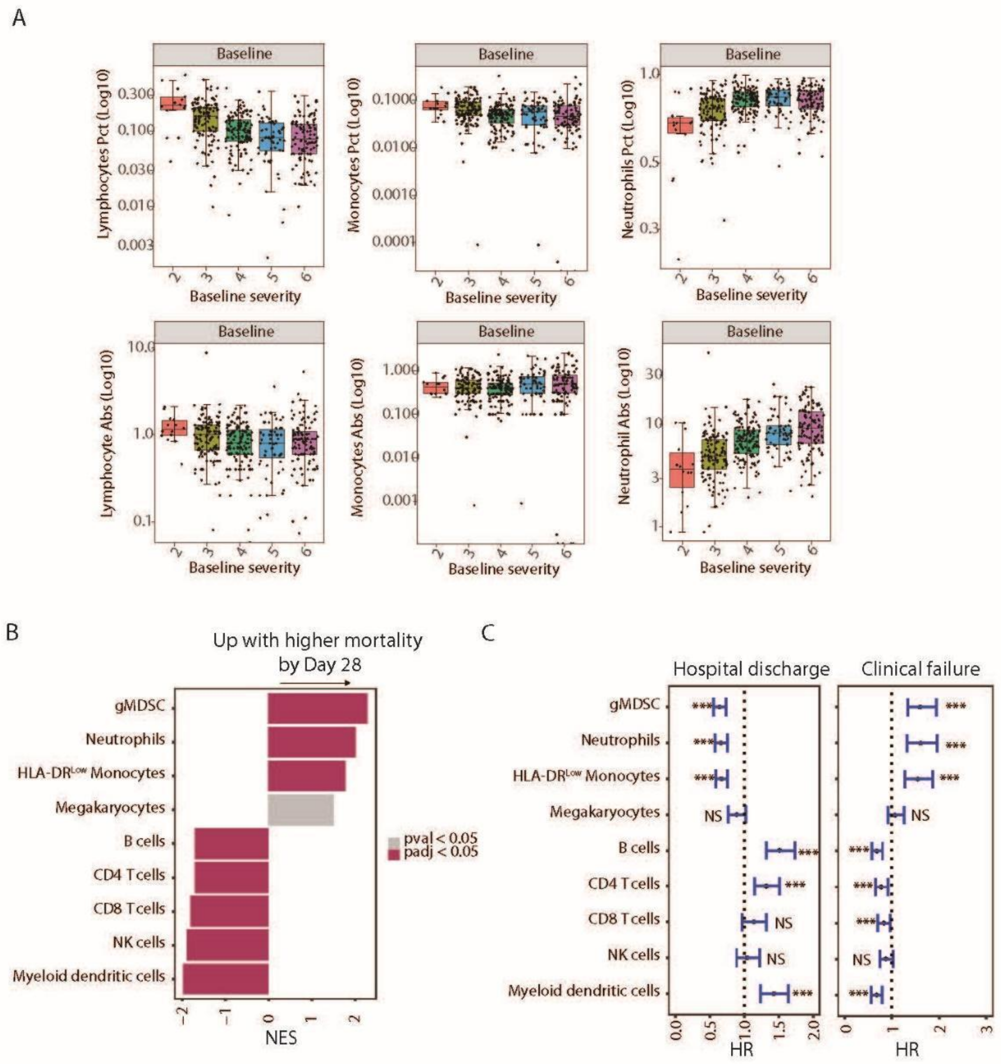
A
Baseline
Lymphocytes Pct (Log10)
Monocytes Pct (Log10)
Neutrophils Pct (Log10)
Lymphocyte Abs (Log10)
Monocytes Abs (Log10)
Neutrophil Abs (Log10)
Baseline severity
B
Up with higher mortality by Day 28
gMDSC
Neutrophils
HLA-DRLow Monocytes
Megakaryocytes
B cells
CD4 Tcells
CD8 Tcells
NK cells
Myeloid dendritic cells
pval < 0.05
padj < 0.05
NES
C
Hospital discharge
Clinical failure
HR
NS
***

**Fig S5. Time to symptom onset–dependent changes in protein and transcript levels at baseline.** (**A)** Column plot showing fraction of samples per baseline severity score sampled within 10 days of symptom onset vs later. (**B)** Column plot showing $\log_2$fold difference in the abundance (Y axis) for the serum proteins significantly different between COVID-19 subjects sampled within 10 days (n = 182) of symptom onset vs later (n = 202). (**C)** Line plot showing baseline serum levels of specified proteins on Y axis and time from symptom onset of the corresponding patients. Error bars represent 95% confidence intervals around the mean. (**D)** Bar plot showing fast gene set enrichment analysis results for immune pathways enriched among genes differentially expressed between COVID-19 subjects sampled within 10 days from symptom onset (early, n = 186) vs greater than 10 days of from symptom onset (late, n = 214).

IFN, interferon; LPS, lipopolysaccharide; NES, normalized enrichment score; NK, natural killer; NPX, normalized protein expression; padj, adjusted *P* value.

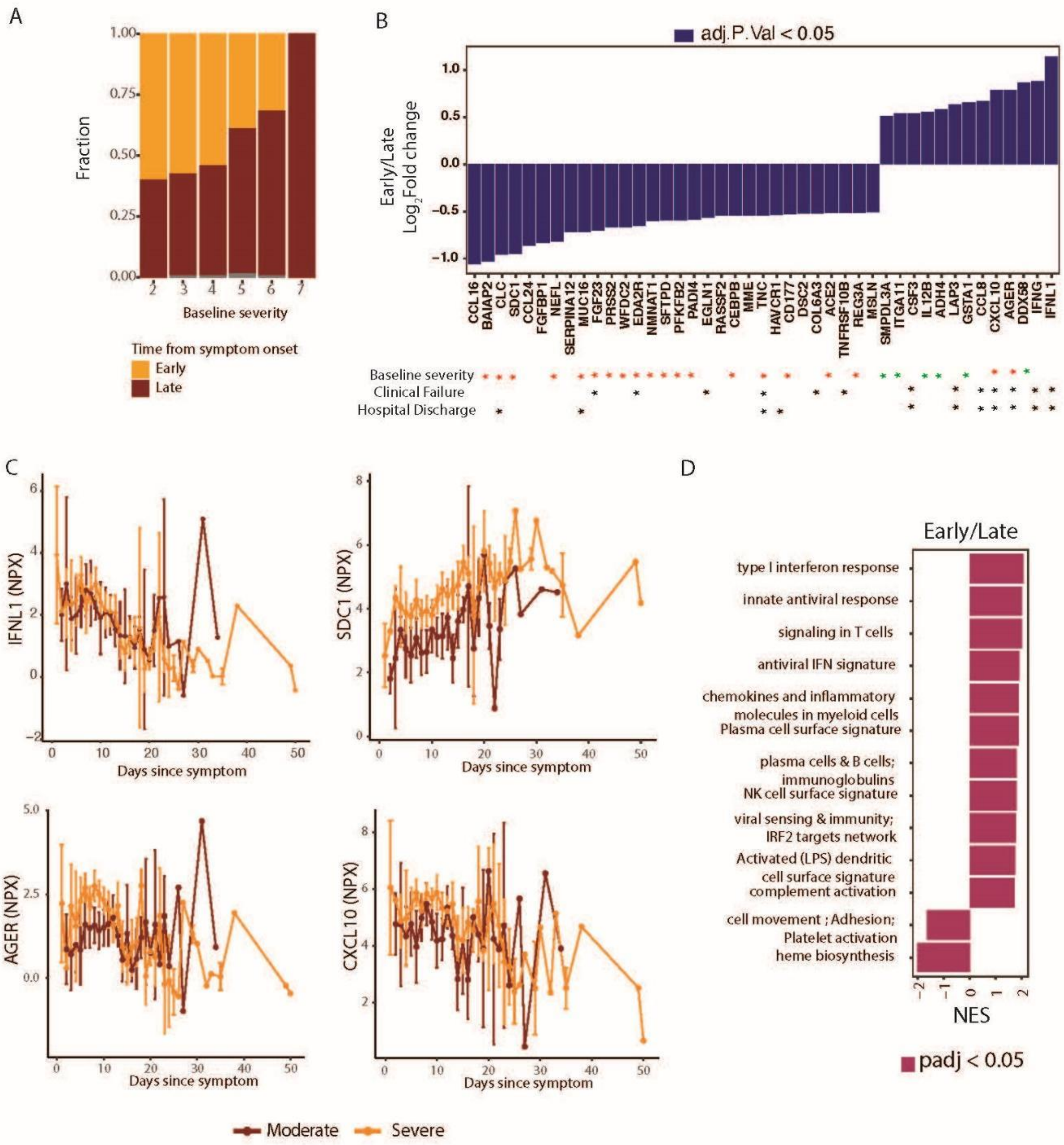

A
Fraction
1.00
0.75
0.50
0.25
0.00
2
3
4
5
6
7
Baseline severity
Time from symptom onset
Early
Late
B
adj.P.Val < 0.05
Early/Late
Log2Fold change
1.0
0.5
0.0
-0.5
-1.0
CCL16
BAIAP2
CLC
SDC1
CCL24
FGFBP1
NEFL
SERPINA12
MUC16
FGF23
PRSS2
WFDC2
EDA2R
NMNAT1
SFTPD
PFKFB2
PADI4
EGLN1
RASSF2
CEBPB
MME
TNC
HAVCR1
CD177
DSC2
COL6A3
ACE2
TNFRSF10B
REG3A
MSLN
SMPDL3A
ITGA11
CSF3
IL12B
ADH4
LAP3
GSTA1
CCL8
CXCL10
AGER
DDX58
IFNG
IFNL1
Baseline severity
Clinical Failure
Hospital Discharge
C
IFNL1 (NPX)
SDC1 (NPX)
AGER (NPX)
CXCL10 (NPX)
Days since symptom
Moderate
Severe
D
Early/Late
type I interferon response
innate antiviral response
signaling in T cells
antiviral IFN signature
chemokines and inflammatory
molecules in myeloid cells
Plasma cell surface signature
plasma cells & B cells;
immunoglobulins
NK cell surface signature
viral sensing & immunity;
IRF2 targets network
Activated (LPS) dendritic
cell surface signature
complement activation
cell movement ; Adhesion;
Platelet activation
heme biosynthesis
NES
padj < 0.05

**Fig S6. PCA analysis evaluating the benefit of tocilizumab treatment. (A)** PCA (top) and density plots (bottom) showing longitudinal changes in serum protein signal measured by Olink for tocilizumab-treated and placebo samples collected at baseline, day 3, day 7, day 28, and healthy controls. **(B)** Cumulative distribution function plot showing distribution of Olink data across indicated Olink treatment groups. *Indicates $P$ <0.05 from Kolmogorov-Smirnov test performed between the indicated timepoints relative to baseline. **(C)** Cumulative distribution function plot showing distribution of indicated RNA-seq treatment groups. *Indicates $P$ value <0.05 from Kolmogorov-Smirnov test performed between the indicated timepoints relative to baseline.

CDF, cumulative distribution function; CTRL, control; PBO, placebo; PCA, principal component analysis; TCZ, tocilizumab.

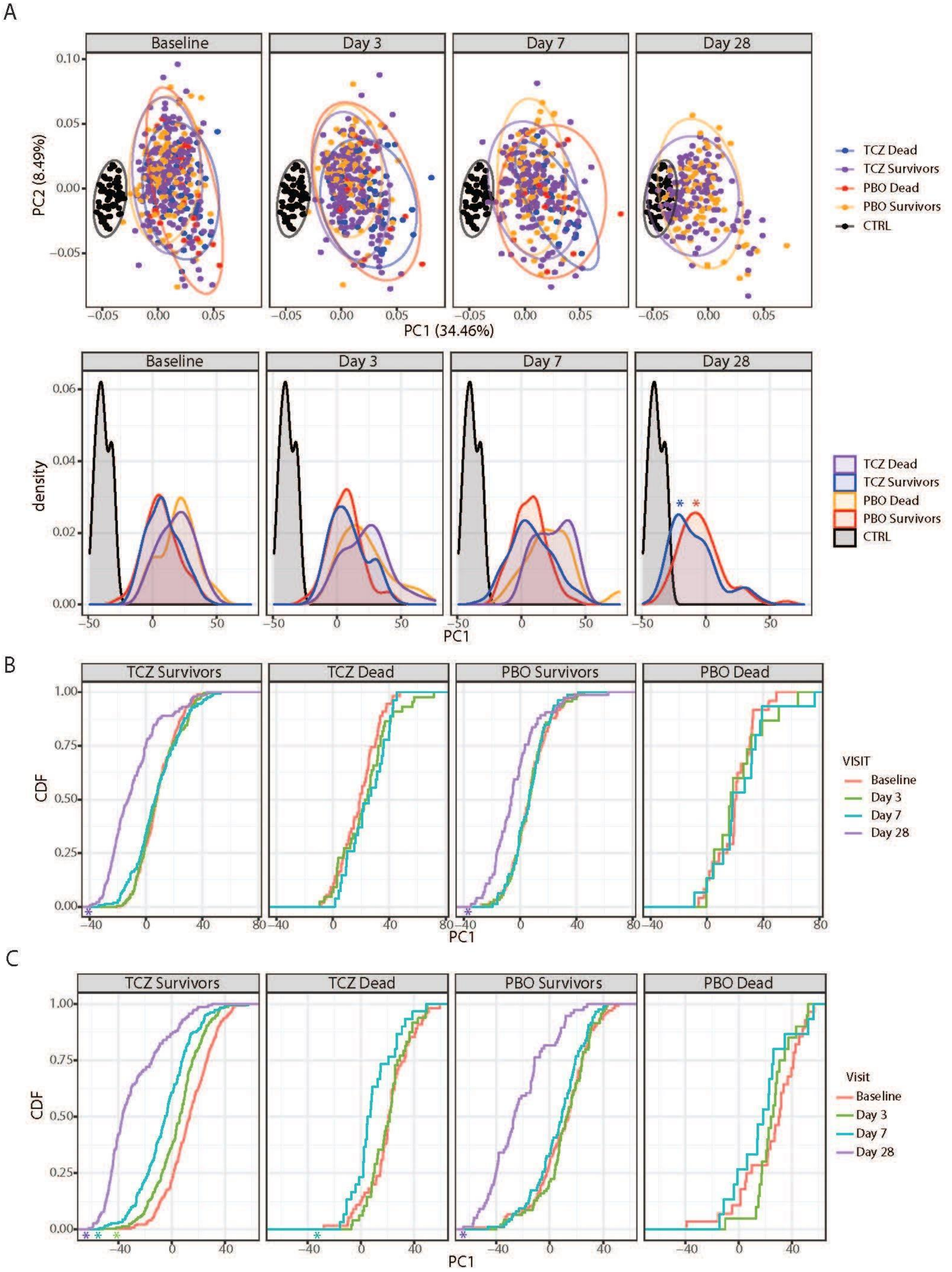

A
Baseline
Day 3
Day 7
Day 28
PC2 (8.49%)
PC1 (34.46%)
TCZ Dead
TCZ Survivors
PBO Dead
PBO Survivors
CTRL
density
PC1
B
TCZ Survivors
TCZ Dead
PBO Survivors
PBO Dead
CDF
PC1
VISIT
Baseline
Day 3
Day 7
Day 28
C
Visit

**Fig S7. Tocilizumab treatment leads to upregulation of genes prognostic for better clinical outcomes.** Forest plot showing hazard ratios with 95% confidence intervals identified using Cox proportional hazard model depicting clinical failure (left) from COVID-19 and time to hospital discharge (right) for pathways shown in Figure 5D. ***Represents Benjamini-Hochberg adjusted *P* <0.05. Only patients with moderate severity (Baseline severity score < 4) were included for the analysis.

HR, hazard ratio.

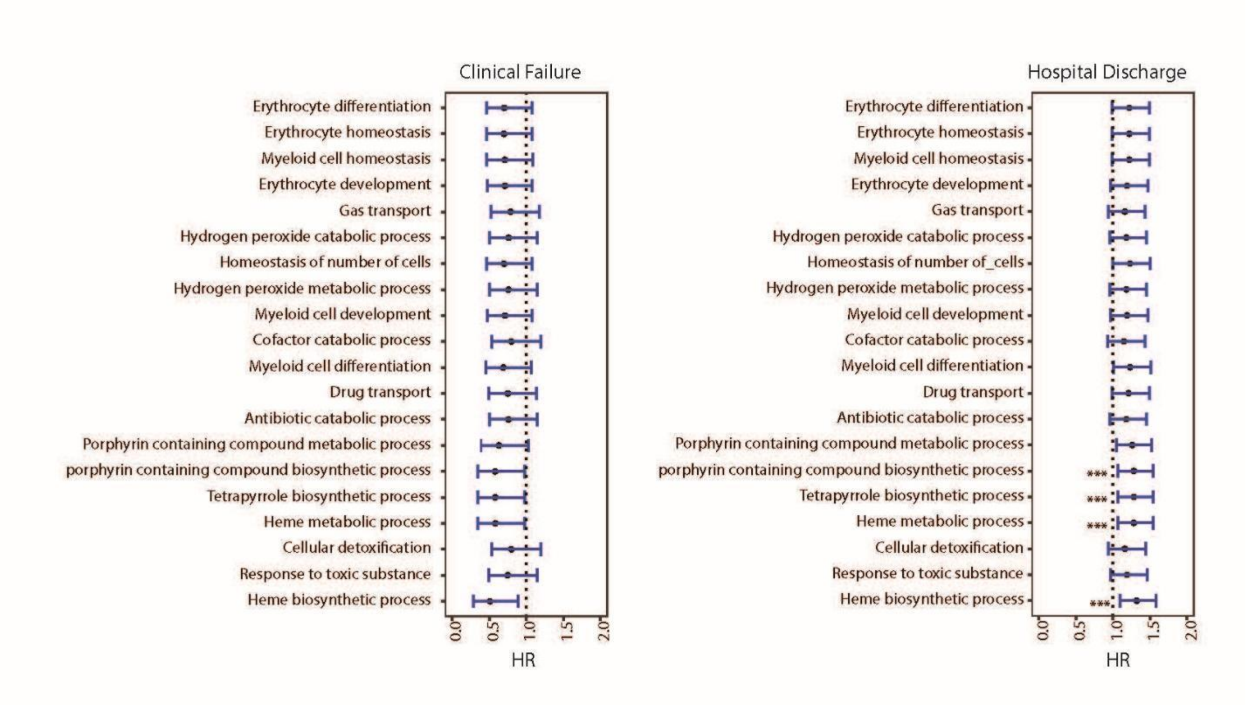

**Fig S8. Effect of tocilizumab treatment on immune pathways. (A)** Bar plot showing immune pathways enriched among genes showing a greater response to tocilizumab compared to placebo by day 3 (calculated as [tocilizumab day 3 – tocilizumab day 1] – [placebo day 3 – placebo day 1]). **(B)** Bar plot showing fast gene set enrichment analysis results for immune pathways enriched among genes responsive to tocilizumab and placebo by Day 7.

CD, cluster of differentiation; HLA, human leukocyte antigen; IFN, interferon; IRF, interferon regulatory factors; ITK, IL-2 inducible T cell kinase; LPS, lipopolysaccharide; NES, normalized enrichment score; NK, natural killer; padj, adjusted *P* value; PKC, protein kinase C; PBO, placebo; TCZ, tocilizumab; Th, T helper cell; TLR, toll-like receptor.

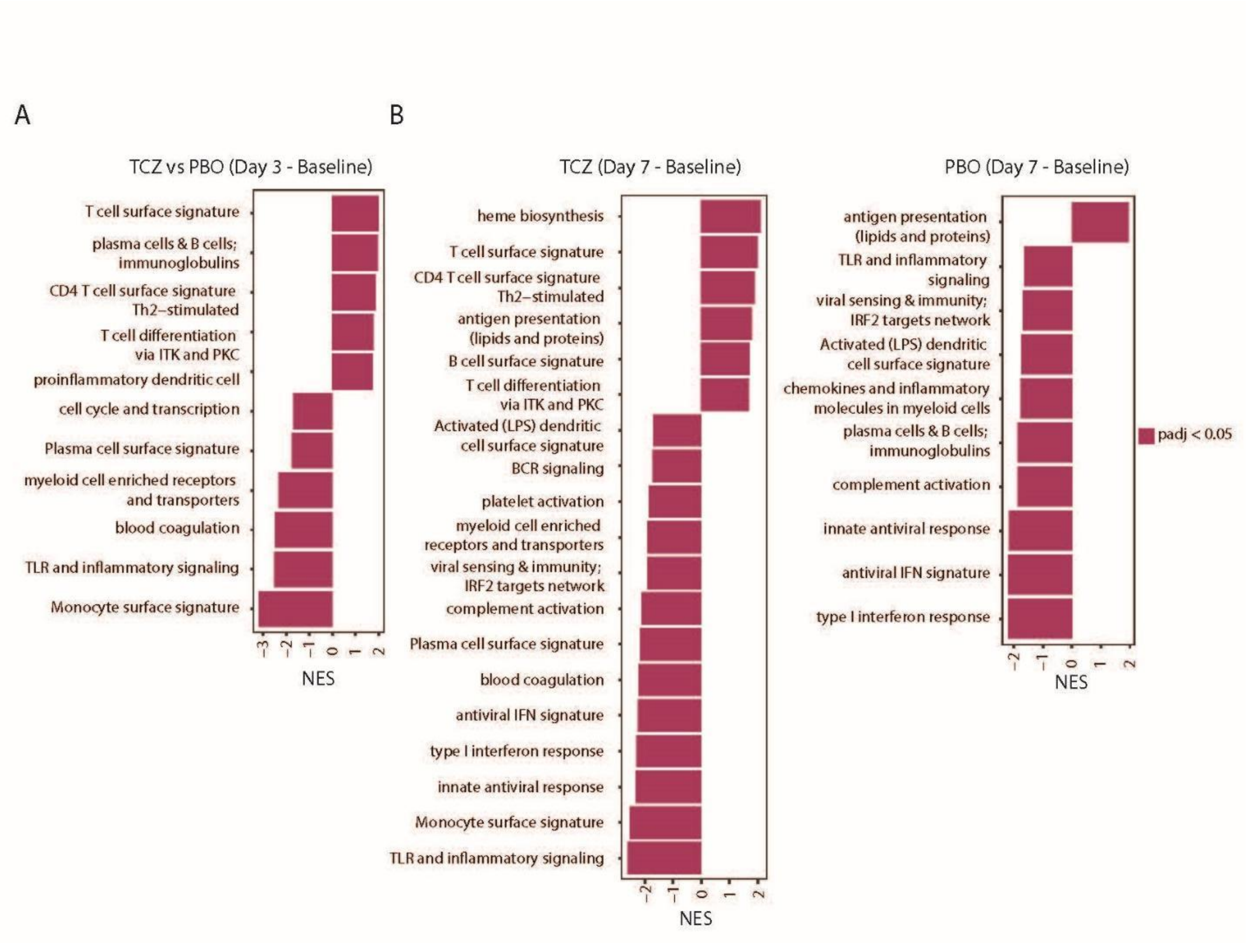

**Fig S9. Effect of tocilizumab treatment on blood cell counts.** Box plot showing eigengene expression of gene sets corresponding to blood cell types for patients treated with tocilizumab and placebo across timepoints. *Represents statistical significance using pairwise T tests at each timepoint using baseline as reference the reference group. * ≤0.05, ** ≤0.01, *** ≤0.001, and **** ≤0.0001. Number of patients per time point are indicated in Supplementary Table 5.

Abs, absolute; NS, not significant; Pct, percent.

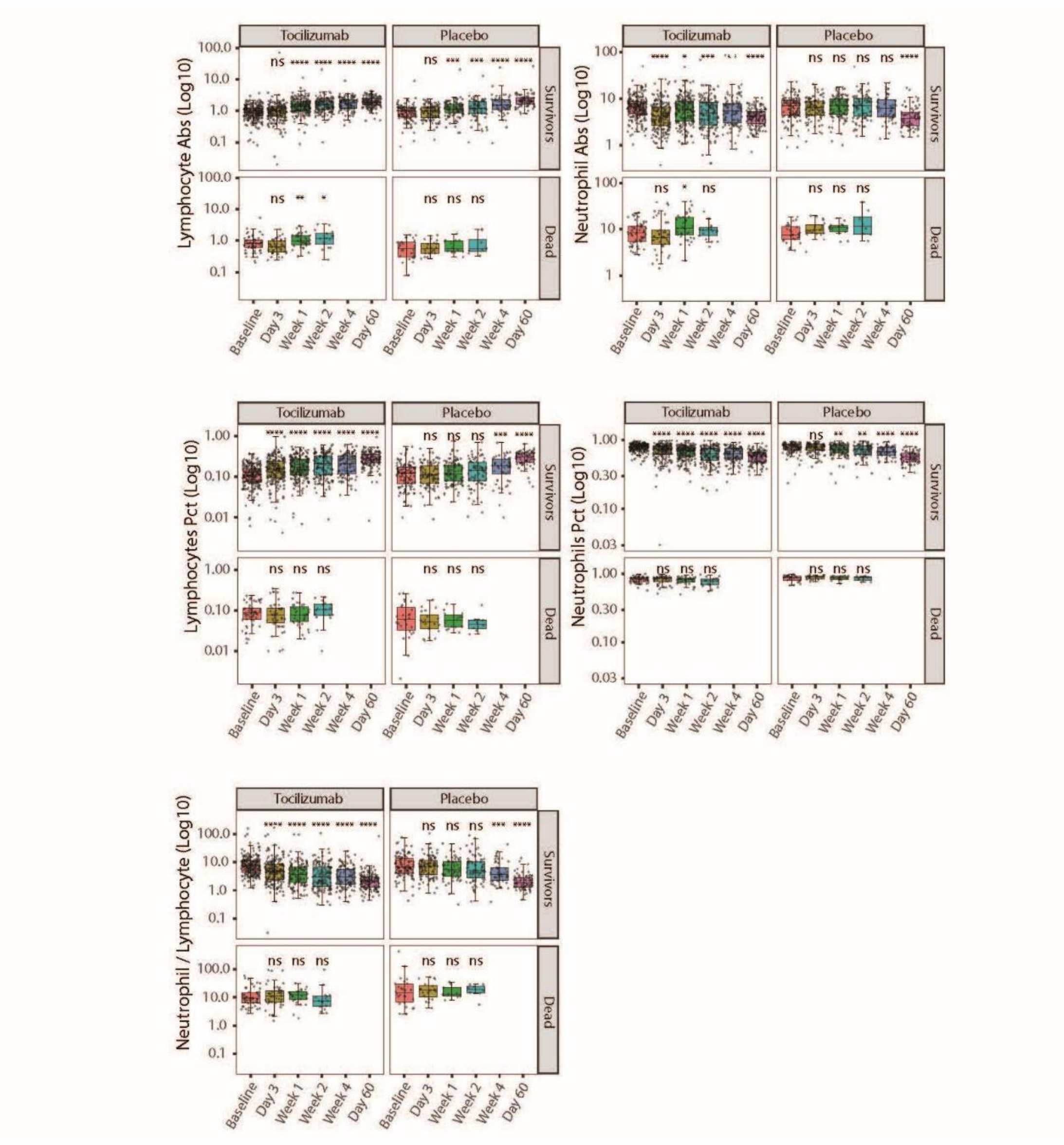

**Fig S10. Effect of tocilizumab treatment on cell type signatures. (A)** Bar plot showing cell types enriched among genes showing a greater response to tocilizumab compared to placebo by day 3 (calculated as [tocilizumab day 3 – tocilizumab day 1] – [placebo day 3 – placebo day 1]). **(B)** Bar plots showing fast gene set enrichment analysis results for blood cell types enriched among genes responsive to tocilizumab and placebo treatment by day 7. (C) Bar plot showing cell types enriched among genes showing a greater response to tocilizumab compared to placebo by day 7 (calculated as [tocilizumab day 7 – tocilizumab day 1] – [placebo day 7 – placebo day 1]) for survivors compared to those who died by day 28. (D) Line plots showing eigengene expression of gene sets corresponding to blood cell types for cases treated with tocilizumab and placebo across timepoints facetted by patients surviving and those dead by day 28.

CD, cluster of differentiation; gMDSC, granulocytic myeloid-derived suppressor cells; HLA, human leukocyte antigen; mMDSC, monocytic myeloid-derived suppressor cells; NES, normalized enrichment score; NK, natural killer; padj, adjusted *P* value; pval, *P* value; PBO, placebo; TCZ, tocilizumab.

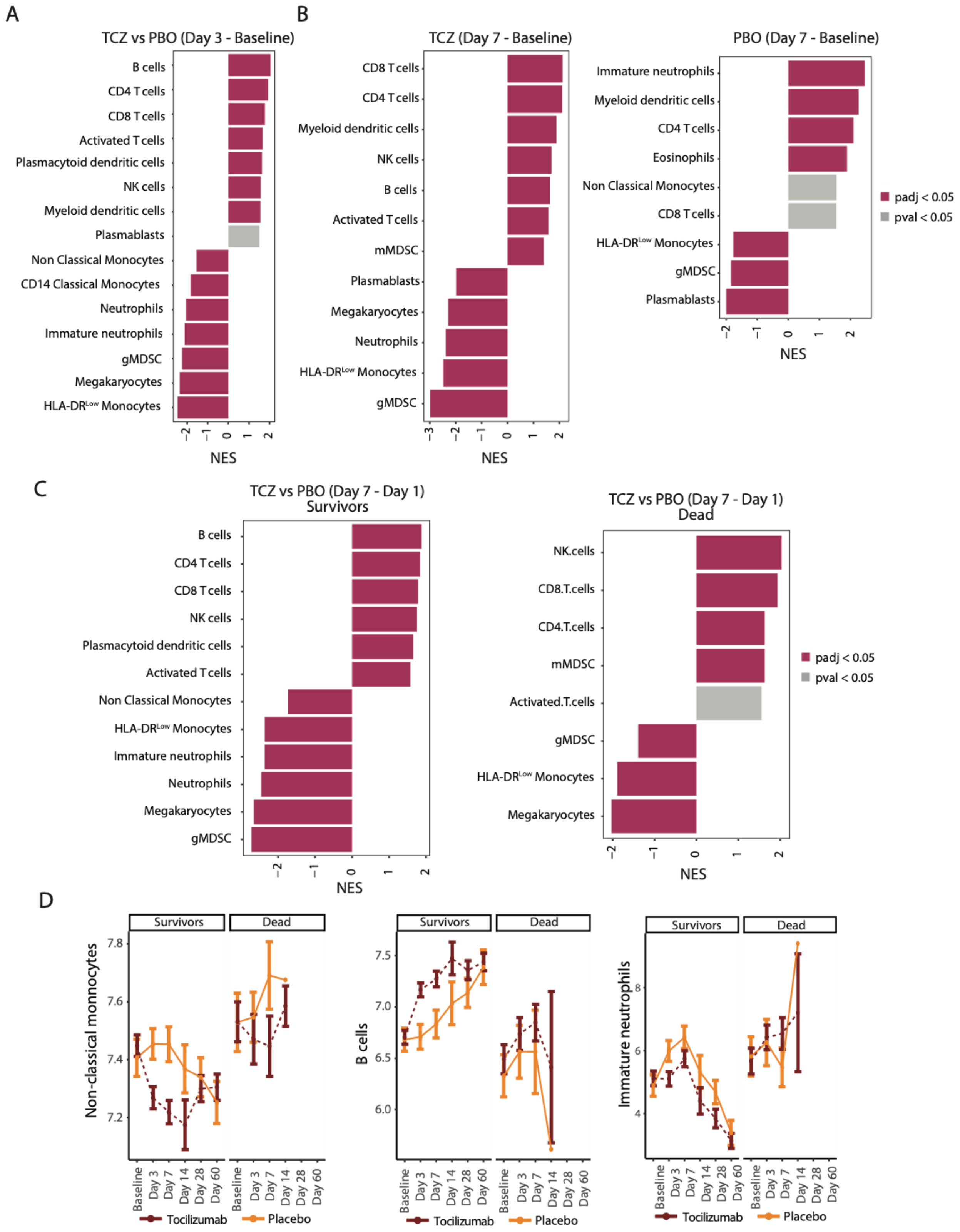

A
TCZ vs PBO (Day 3 - Baseline)
B cells
CD4 T cells
CD8 T cells
Activated T cells
Plasmacytoid dendritic cells
NK cells
Myeloid dendritic cells
Plasmablasts
Non Classical Monocytes
CD14 Classical Monocytes
Neutrophils
Immature neutrophils
gMDSC
Megakaryocytes
HLA-DR^Low Monocytes
NES
B
TCZ (Day 7 - Baseline)
CD8 T cells
CD4 T cells
Myeloid dendritic cells
NK cells
B cells
Activated T cells
mMDSC
Plasmablasts
Megakaryocytes
Neutrophils
HLA-DR^Low Monocytes
gMDSC
NES
PBO (Day 7 - Baseline)
Immature neutrophils
Myeloid dendritic cells
CD4 T cells
Eosinophils
Non Classical Monocytes
CD8 T cells
HLA-DR^Low Monocytes
gMDSC
Plasmablasts
NES
padj < 0.05
pval < 0.05
C
TCZ vs PBO (Day 7 - Day 1)
Survivors
B cells
CD4 T cells
CD8 T cells
NK cells
Plasmacytoid dendritic cells
Activated T cells
Non Classical Monocytes
HLA-DR^Low Monocytes
Immature neutrophils
Neutrophils
Megakaryocytes
gMDSC
NES
TCZ vs PBO (Day 7 - Day 1)
Dead
NK.cells
CD8.T.cells
CD4.T.cells
mMDSC
Activated.T.cells
gMDSC
HLA-DR^Low Monocytes
Megakaryocytes
NES
padj < 0.05
pval < 0.05
D
Survivors
Dead
Non-classical monnocytes
B cells
Immature neutrophils
Baseline
Day 3
Day 7
Day 14
Day 28
Day 60
Tocilizumab
Placebo

**Fig S11. Effect of corticosteroid treatment on blood transcript and serum protein levels at baseline. (A)** Bar plot showing immune pathways enriched among genes showing differential expression between subjects treated with (n = 93) or without corticosteroids (n = 316) sampled at baseline. **(B)** Column plot showing $\log_2$fold difference in the abundance (Y axis) for the serum proteins significantly different at baseline between COVID-19 subjects untreated (n = 304) or treated with corticosteroids (n = 84) at baseline.

CD, cluster of differentiation; DC, dendritic cell; IFN, interferon; ITK, IL-2 inducible T cell kinase; NES, normalized enrichment score; PKC, protein kinase C; pval, *P* value; Th, T helper cell; TLR, toll-like receptor.

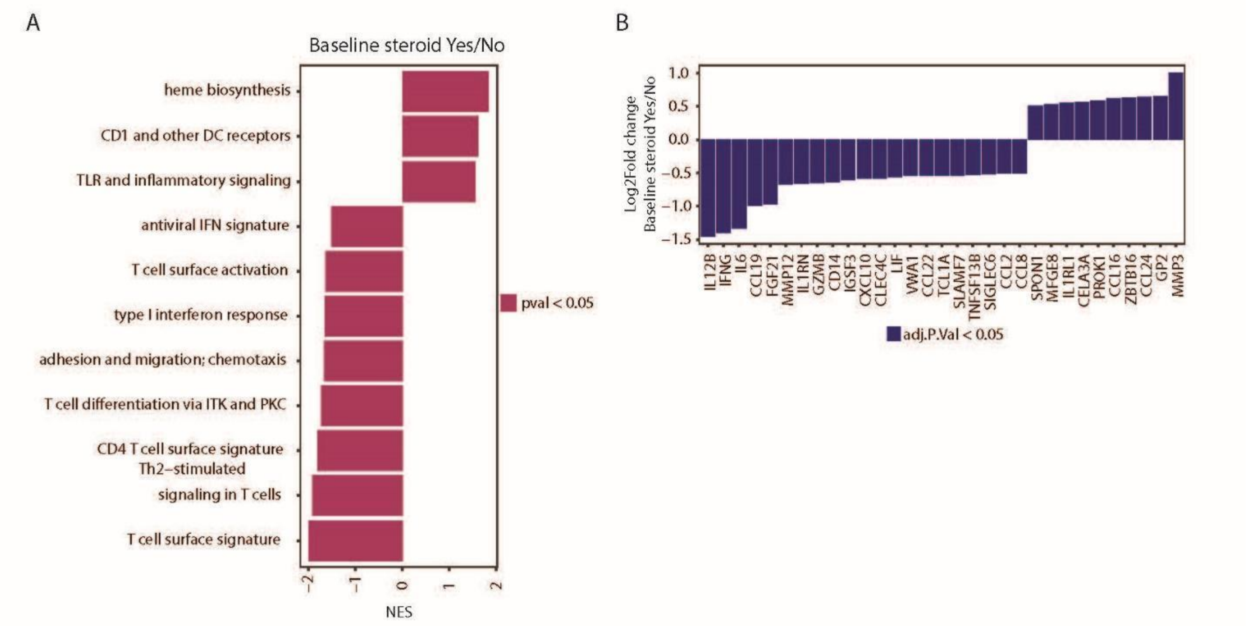

**Fig S12. Quality assessment of blood cell type gene sets. (A)** Scatter plots showing correlation between the eigengene values of gene sets generated from pooled samples from whole blood and PBMCs (Y axis) compared to the eigengene values from PBMCs. The cell types corresponding to the gene sets are indicated on the axis labels. **(B)** Scatter plots showing correlation between the eigengene values of mature neutrophils and nonclassical monocytes gene sets generated in this work (X axis) compared to the eigengene values from published gene sets (Y axis). The cell types corresponding to the gene sets are indicated on the axis labels. **(C)** Scatter plots showing correlation between the eigengene values of gene sets derived in this study (X axis) compared to clinical hematology-based blood cell frequency. The cell types corresponding to the gene sets are indicated on axis labels.

CD4, cluster of differentiation 4; HLA, human leukocyte antigen; NK, natural killer; PBMC, peripheral blood mononuclear cell; WB, whole blood.

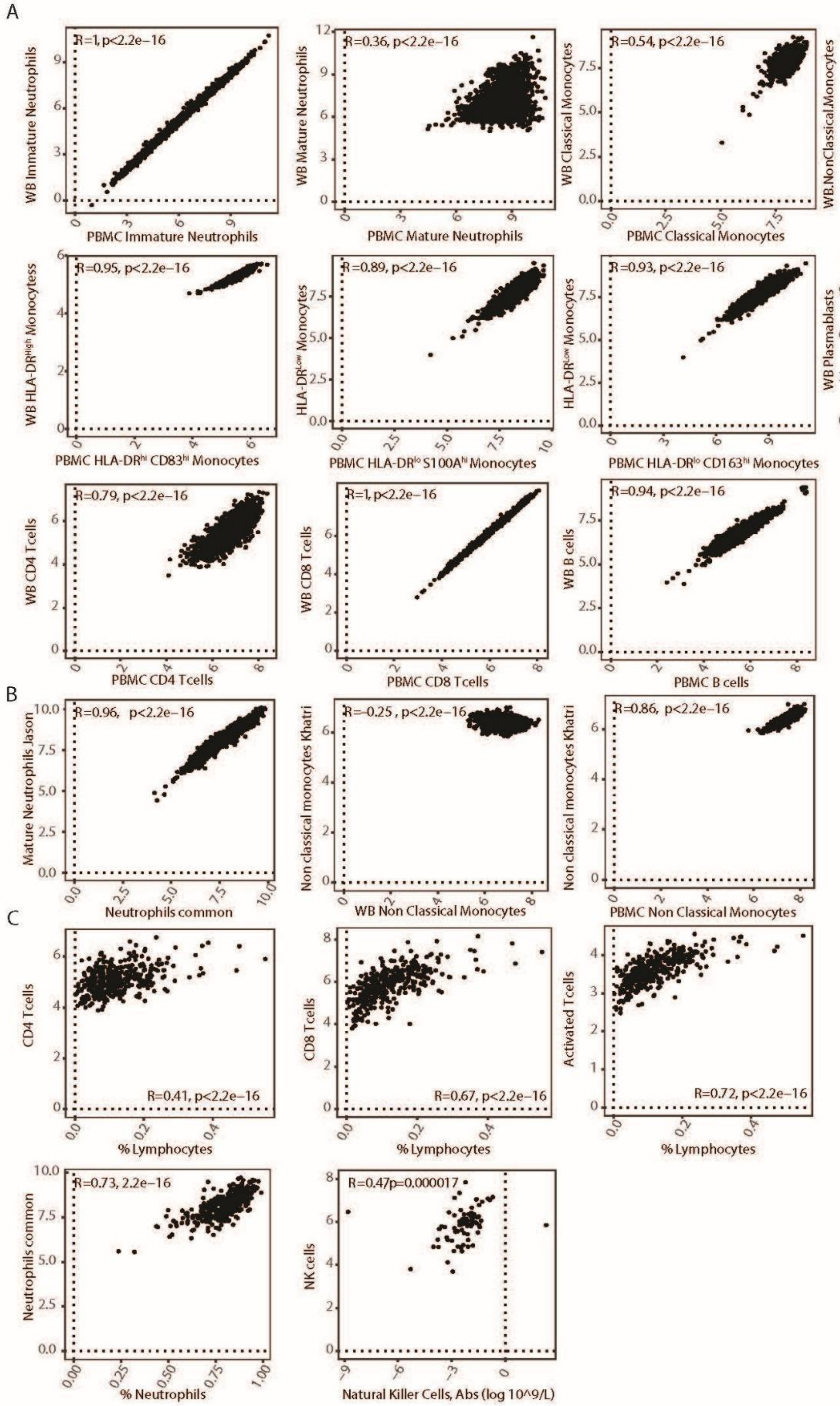
A
R=1, p<2.2e−16
WB Immature Neutrophils
PBMC Immature Neutrophils
R=0.36, p<2.2e−16
WB Mature Neutrophils
PBMC Mature Neutrophils
R=0.54, p<2.2e−16
WB Classical Monocytes
PBMC Classical Monocytes
R=−0.094p=0.00014
WB.NonClassical.Monocytes
PBMC Non Classical Monocytes
R=0.95, p<2.2e−16
WB HLA-DRHigh Monocytess
PBMC HLA-DRhi CD83hi Monocytes
R=0.89, p<2.2e−16
HLA-DRLow Monocytes
PBMC HLA-DRlo S100Ahi Monocytes
R=0.93, p<2.2e−16
HLA-DRLow Monocytes
PBMC HLA-DRlo CD163hi Monocytes
R=1, p<2.2e−16
WB Plasmablasts
PBMC Plasmablasts
R=0.79, p<2.2e−16
WB CD4 Tcells
PBMC CD4 Tcells
R=1, p<2.2e−16
WB CD8 Tcells
PBMC CD8 Tcells
R=0.94, p<2.2e−16
WB B cells
PBMC B cells
B
R=0.96, p<2.2e−16
Mature Neutrophils Jason
Neutrophils common
R=-0.25 , p<2.2e−16
Non classical monocytes Khatri
WB Non Classical Monocytes
R=0.86, p<2.2e−16
Non classical monocytes Khatri
PBMC Non Classical Monocytes
C
CD4 Tcells
R=0.41, p<2.2e−16
% Lymphocytes
CD8 Tcells
R=0.67, p<2.2e−16
% Lymphocytes
Activated Tcells
R=0.72, p<2.2e−16
% Lymphocytes
R=0.73, 2.2e−16
Neutrophils common
% Neutrophils
R=0.47p=0.000017
NK cells
Natural Killer Cells, Abs (log 10^9/L)

**Fig S13. Quality assessment of Olink data. (A)** Scatter plot between interquartile range and median calculated across all samples for 1472 proteins split by the representative panels. **(B)** PCA analysis plotting PC1 and PC2 for 4 Olink panels where each point represents a sample. **(C)** Histogram plot wherein the x axis represents the fraction of assays per sample with NPX values above LOD. Y axis represents the number of samples. Two samples are highlighted where the fraction of assays with NPX above LOD were <75%. These were removed from downstream analysis. **(D)** Histogram plot wherein the x axis represents the fraction of assays per sample that pass Olink's internal QC. Y axis represents the number of samples. 86 samples are highlighted where the fraction of assays that pass QC were <75%. These were removed from downstream analysis. LOD, limit of detection; PCA, principal component analysis; QC, quality control.

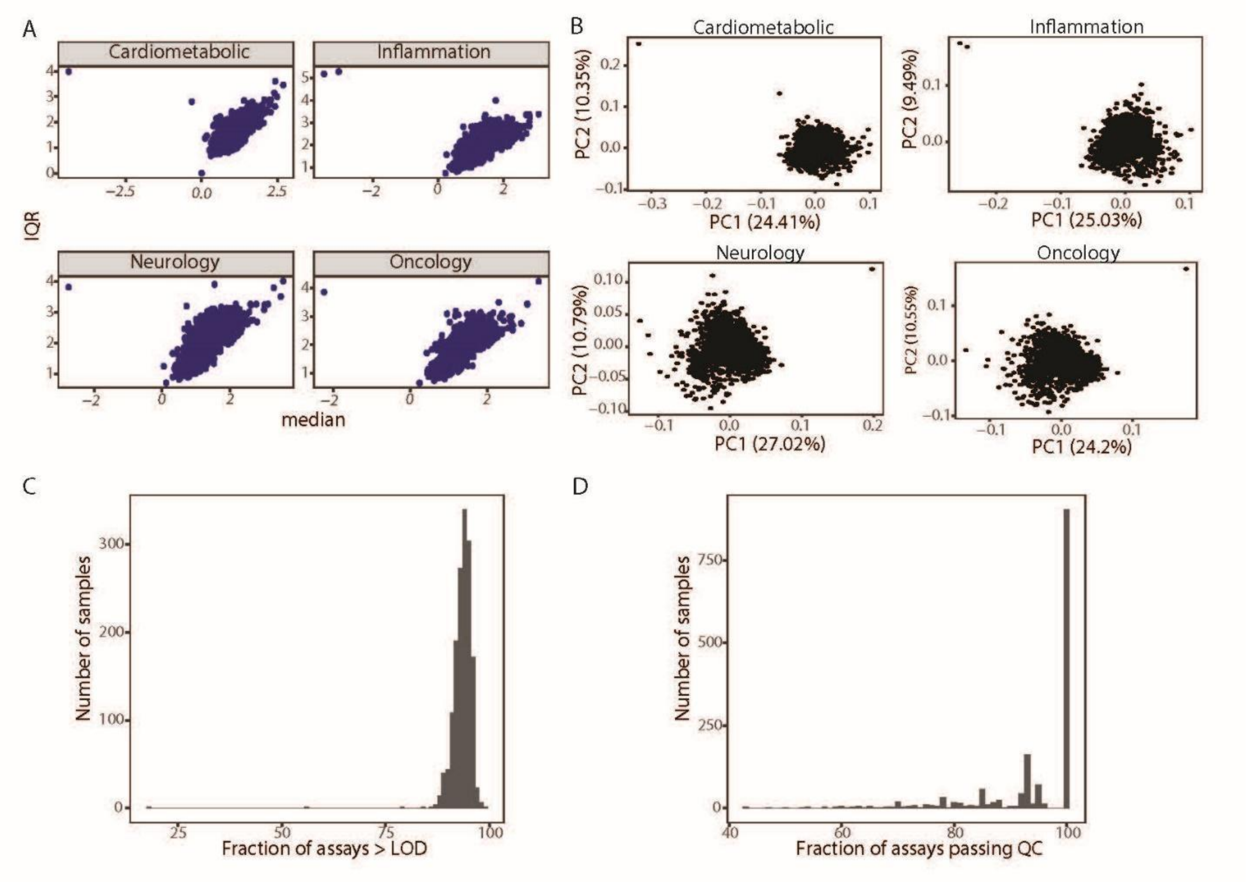

**Supplemental Table 1. $Log_2$fold change outputs from longitudinal Limma analysis for severity-associated proteins.**

Each row represents a severity-associated protein and each column represents the indicated comparison analyzed with Limma. For each column, logFC corresponding to only the significantly different proteins (Benjamini-Hochberg adjusted $P < 0.05$, Limma) are listed.

FC, fold change; NA, not assessed; PBO, placebo; TCZ, tocilizumab.

| Description | Uniprot ID | severe_up | TCZ (Day 3 - baseline) | TCZ (Day 7 - baseline) | TCZ (Day 28 - baseline) | TCZ (Day 60 - baseline) | PBO (Day 3 - baseline) | PBO (Day 7 - baseline) | PBO (Day 28 - baseline) | PBO (Day 60 - baseline) | Day 60 – Healthy control |
|---|---|---|---|---|---|---|---|---|---|---|---|
| ACE2 | Q9BYF1 | 0.63585976 | 0.51141781 | 0.58367898 | 0.4637277 | -1.2984634 | NA | 0.67412558 | NA | -1.0671755 | 0.74514044 |
| ACTA2 | P62736 | 0.78102619 | NA | NA | NA | -0.3899357 | NA | NA | NA | NA | 0.96862137 |
| AZU1 | P20160 | 0.6099465 | -0.4886078 | -0.380442 | -0.3059177 | -0.6701125 | NA | NA | NA | NA | 0.84636259 |
| NTproBNP | NA | 0.79334476 | NA | NA | NA | -0.647875 | NA | NA | NA | NA | 0.91566419 |
| CA3 | P07451 | 0.67417169 | NA | NA | -1.0791931 | -1.2117366 | NA | NA | -0.9933426 | -1.1590286 | 0.36755541 |
| CCDC80 | Q76M96 | 0.86931165 | NA | NA | -0.4272077 | -0.4684429 | NA | NA | NA | NA | 0.39741318 |
| CCL15 | Q16663 | 0.56325393 | NA | NA | -0.3536969 | -0.5838196 | NA | NA | NA | -0.4070918 | 0.31768284 |
| CEBPB | P17676 | 0.94677173 | NA | NA | NA | -0.5820625 | NA | NA | NA | NA | 0.57211456 |
| CGREF1 | Q99674 | 0.70680769 | NA | NA | -0.2481254 | -0.3319795 | NA | NA | NA | NA | 0.26852912 |
| CLEC5A | Q9NY25 | 0.69888154 | NA | NA | -0.5412185 | -0.9611188 | NA | NA | NA | -0.7457714 | 0.29744655 |
| COL6A3 | P12111 | 0.72462936 | 0.31212146 | 0.54810284 | 0.75165538 | 0.71161071 | NA | 0.53576047 | 0.88363088 | 0.8708449 | 1.07447341 |
| CTSB | P07858 | 0.86292858 | NA | NA | NA | -0.487025 | NA | NA | NA | -0.4221633 | 0.4454941 |
| CTSL | P07711 | 0.77816681 | NA | -0.5180193 | -1.10839 | -1.4113607 | NA | NA | -0.9440103 | -1.3730694 | 0.34248785 |
| CXCL16 | Q9H2A7 | 0.58178182 | NA | -0.402329 | -0.4783531 | -0.6756259 | NA | NA | -0.3336074 | -0.5639878 | 0.24318522 |
| DEFA1 | P59665 | 0.80186855 | NA | 0.31170852 | NA | -0.9459848 | NA | NA | NA | -0.7716918 | 0.45009563 |
| FABP4 | P15090 | 0.61285933 | NA | 0.50577784 | 0.57789385 | NA | NA | NA | 0.88948088 | NA | 0.72888343 |
| GDF15 | Q99988 | 1.06703484 | NA | NA | -0.9197608 | -1.5302705 | NA | NA | -0.7240529 | -1.1618816 | 1.23732392 |
| GP2 | P55259 | 0.50951997 | NA | NA | NA | -0.8137991 | NA | NA | NA | -0.9967469 | NA |
| GPR37 | O15354 | 0.88093642 | -0.5405557 | -0.7992426 | -1.0239331 | -1.4005446 | NA | NA | -0.7176147 | -1.3159327 | 0.53360726 |
| HMOX1 | P09601 | 0.70882502 | NA | -0.2763261 | -1.2092308 | -1.5859705 | NA | NA | -0.6808676 | -1.169802 | NA |
| HSPG2 | P98160 | 0.53273473 | 0.2364242 | 0.39419261 | 0.59922769 | 0.56312589 | NA | NA | 0.59118235 | 0.65014286 | 0.50407365 |
| IGFBP1 | P08833 | 1.05008775 | NA | -0.704408 | -1.1137185 | -1.275233 | NA | NA | NA | -1.4180388 | NA |
| IGFBP2 | P18065 | 0.52154324 | NA | -0.3197653 | -0.8072454 | -1.0940045 | NA | NA | -0.4845765 | -0.9272755 | 0.72069661 |
| IL1RL1 | Q01638 | 1.12177177 | -0.2889361 | -0.7370864 | -1.46908 | -2.095108 | NA | NA | -1.3242618 | -1.8930571 | NA |

| | | | | | | | | | | | |
|---|---|---|---|---|---|---|---|---|---|---|---|
| LBP | P18428 | 0.56351577 | -1.332321 | -2.0707648 | -1.90541 | -2.0716196 | NA | NA | -1.2912426 | -1.9309612 | NA |
| LCN2 | P80188 | 0.63771589 | NA | NA | NA | -0.4152804 | NA | NA | NA | NA | 0.51700974 |
| LDLR | P01130 | 0.59675376 | 0.49706073 | 0.43833807 | -0.2084838 | -0.4717161 | NA | NA | NA | NA | 0.43234934 |
| LTBP2 | Q14767 | 0.8483546 | NA | -0.3550318 | -0.4814085 | -1.0780179 | NA | NA | NA | -0.7327653 | 0.79622127 |
| MB | P02144 | 1.43621553 | NA | NA | -1.0412892 | -1.4076384 | NA | NA | -1.0570574 | -1.7882143 | -0.4777072 |
| MNDA | P41218 | 0.63821437 | -0.4594183 | NA | -0.4798869 | -1.0935875 | NA | NA | NA | -0.8207041 | NA |
| NADK | O95544 | 0.69438072 | -0.3665982 | -0.6465426 | -1.3279862 | -1.894325 | NA | -0.4861872 | -1.0533059 | -1.6377061 | 0.54797773 |
| NPDC1 | Q9NQX5 | 0.51224165 | NA | NA | NA | NA | NA | NA | NA | NA | 0.30678749 |
| OLR1 | P78380 | 0.50114116 | -0.4292215 | NA | 0.34380846 | NA | NA | 0.5874407 | 0.82550882 | NA | 0.73625066 |
| SPP1 | P10451 | 0.76029782 | -0.4735548 | -0.7102347 | -1.3761992 | -1.7806554 | NA | NA | -0.8255897 | -1.4072878 | 0.74550525 |
| PAG1 | Q9NWQ8 | 0.82350036 | -0.2707854 | -0.3682619 | -1.19278 | -1.7153634 | NA | NA | -0.8780868 | -1.5196408 | 0.76043082 |
| PGLYRP1 | O75594 | 0.52952502 | NA | 0.22050511 | NA | -0.540208 | NA | 0.35532558 | NA | -0.3916224 | 0.42137789 |
| PI3 | P19957 | 0.56018944 | NA | NA | NA | NA | NA | NA | NA | NA | 0.72845653 |
| PLA2G1B | P04054 | 0.7673784 | NA | NA | NA | -0.5111375 | NA | NA | NA | NA | NA |
| PLA2G2A | P14555 | 1.38877139 | -2.1372868 | -3.3198091 | -2.9758562 | -3.6286661 | NA | NA | -2.2198029 | -3.4410776 | 0.76038861 |
| PPP1R2 | P41236 | 0.53600984 | NA | NA | -0.7963331 | -1.1828688 | NA | NA | -0.5989382 | -0.9836143 | 0.58210707 |
| PRSS2 | P07478 | 0.91672696 | NA | 0.63847216 | NA | -0.6473295 | NA | NA | NA | -0.5878898 | 0.32718866 |
| PRTN3 | P24158 | 0.63691048 | NA | -0.3027142 | -0.7524746 | -1.4424357 | NA | NA | NA | -1.1915592 | 0.77490757 |
| PTN | P21246 | 1.47907066 | NA | NA | -2.2817208 | -3.4571071 | NA | NA | -2.0459029 | -2.9172306 | 0.47179 |
| RCOR1 | Q9UKL0 | 0.56809265 | NA | NA | -0.5005315 | -0.7837277 | NA | NA | -0.5202235 | -0.7234163 | 0.2530556 |
| REG1A | P05451 | 0.84733552 | NA | 0.4448392 | NA | -0.3943321 | NA | NA | NA | NA | 0.61541155 |
| REG1B | P48304 | 0.95228102 | NA | 0.39353636 | NA | -0.7406929 | NA | NA | NA | -0.7022857 | 0.58279279 |
| REG3A | Q06141 | 0.755352 | NA | 0.49446136 | NA | -0.5425938 | NA | 0.6801 | NA | NA | 1.00572709 |
| REN | P00797 | 0.56030102 | NA | NA | -0.5067277 | -0.8071455 | NA | NA | NA | -0.7346898 | NA |
| RETN | Q9HD89 | 0.7158442 | NA | NA | -0.3795615 | -0.9695491 | NA | NA | NA | -0.7486918 | 0.31565476 |
| RNASE3 | P12724 | 0.59784717 | NA | 0.38593807 | 0.60293923 | NA | NA | NA | 0.81948088 | NA | 1.52277804 |
| S100P | P25815 | 0.6733319 | NA | NA | -0.3322477 | -0.8737036 | NA | NA | NA | -0.6335286 | 0.93638482 |
| SDC1 | P18827 | 1.29886127 | 0.78347991 | 0.8140767 | -0.82723 | -2.4830786 | NA | 0.92911047 | NA | -2.0849694 | 0.62822213 |
| SFTPD | P35247 | 0.81897252 | 0.67474155 | 0.61684375 | NA | -0.2759089 | NA | NA | NA | NA | 1.13681704 |

| | | | | | | | | | | | |
|---|---|---|---|---|---|---|---|---|---|---|---|
| TFPI | P10646 | 0.54793597 | 0.23161416 | 0.17540682 | -0.19224 | -0.403033 | NA | NA | NA | -0.3333204 | 0.37464359 |
| TNC | P24821 | 0.95412882 | NA | -0.368717 | -0.4746962 | -1.0466964 | NA | NA | NA | -0.8145429 | 0.51648042 |
| TNNI3 | P19429 | 0.84139171 | NA | -0.4098006 | -0.8130146 | -0.9863679 | NA | NA | -1.083225 | -1.5112796 | NA |
| TSPAN1 | O60635 | 0.53863268 | NA | 0.38078977 | NA | NA | NA | NA | NA | NA | 0.3421106 |
| CHI3L1 | P36222 | 1.29453889 | -0.6435553 | -0.7842943 | -0.8561308 | -1.1721732 | NA | NA | NA | -1.0170306 | 0.53613972 |
| AGRN | O00468 | 0.60386036 | NA | NA | -0.5226738 | -0.9524402 | NA | NA | NA | -0.7396224 | 0.91345001 |
| AGRP | O00253 | 0.64348882 | NA | -0.3590284 | -0.89867 | -1.3513955 | NA | NA | -0.5072838 | -1.1027286 | NA |
| ANGPTL4 | Q9BY76 | 0.6269842 | NA | NA | -0.5707408 | -0.6503946 | NA | NA | -0.3707632 | -0.6811082 | NA |
| ARTN | Q5T4W7 | 0.513233 | NA | NA | -0.2970754 | -0.4594429 | NA | NA | -0.4118897 | -0.5130796 | NA |
| ATP5IF1 | Q9UII2 | 0.77894094 | NA | -0.2844881 | -0.9594238 | -1.5100491 | NA | NA | -0.7024059 | -1.3388735 | NA |
| CCL20 | P78556 | 1.14205178 | NA | -0.5892699 | -0.9202731 | -1.098717 | NA | NA | -0.6371162 | -0.9448959 | 0.96420883 |
| CCL23 | P55773 | 0.64301479 | -0.5258447 | -0.8056375 | -1.06208 | -1.3711661 | NA | NA | -0.721425 | -1.1778367 | 0.38852242 |
| CCL7 | P80098 | 1.13625453 | NA | -1.153992 | -3.10372 | -3.6350875 | NA | -0.8324884 | -2.3262529 | -3.2338551 | 0.96687553 |
| CD79B | P40259 | 0.59663833 | -0.3468123 | -0.4027653 | -0.3948792 | -0.7057902 | NA | NA | NA | NA | 0.97236887 |
| CEACAM21 | Q3KPI0 | 0.63724936 | NA | NA | NA | -0.7081411 | NA | NA | NA | NA | 0.76914443 |
| CKAP4 | Q07065 | 0.87770259 | NA | -0.5326085 | -1.1777292 | -1.541558 | NA | NA | -0.8349191 | -1.3692102 | 0.54062382 |
| CLEC4D | Q8WXI8 | 0.5940659 | NA | NA | NA | -0.4468884 | NA | NA | NA | NA | 0.6210571 |
| CLEC7A | Q9BXN2 | 0.57458266 | 0.47545845 | 0.61813636 | NA | NA | NA | NA | NA | NA | 0.62071034 |
| CRELD2 | Q6UXH1 | 0.5884858 | NA | -0.2242733 | -0.4977723 | -0.8361964 | NA | NA | NA | -0.7722429 | 0.63097245 |
| CST7 | O76096 | 0.52091557 | NA | -0.4517057 | -1.45644 | -2.0768938 | NA | NA | -1.3746191 | -2.0196184 | 0.50259909 |
| CXCL17 | Q6UXB2 | 0.59801887 | NA | NA | -0.3934054 | -0.7831277 | NA | NA | NA | -0.5521102 | 0.45551461 |
| CXCL9 | Q07325 | 0.82688956 | 0.87713151 | 0.35301193 | -0.8588831 | -1.1360991 | NA | NA | NA | -1.1249816 | 1.17052826 |
| EGLN1 | Q9GZT9 | 0.92452173 | NA | NA | NA | -0.9668759 | NA | NA | NA | -0.6401592 | 0.79927169 |
| ENAH | Q8N8S7 | 0.63388677 | NA | NA | -0.2324823 | -0.354042 | NA | NA | NA | NA | 0.8564763 |
| EPO | P01588 | 0.61512089 | NA | NA | NA | NA | NA | NA | NA | NA | 0.3883022 |
| ESM1 | Q9NQ30 | 0.56391977 | NA | -0.3180955 | -1.2438131 | -1.7042339 | NA | NA | -1.0465412 | -1.6816429 | NA |
| FSTL3 | O95633 | 0.67831872 | NA | NA | -0.3887992 | -0.625633 | NA | NA | NA | -0.5702347 | 0.58271337 |
| GBP2 | P32456 | 0.71519422 | NA | -0.4661074 | -1.6865792 | -2.1115518 | NA | -0.7822919 | -1.5126721 | -1.8112449 | 0.77191969 |
| HGF | P14210 | 1.2524268 | NA | NA | -1.0627962 | -1.7128661 | NA | NA | -0.7934382 | -1.381049 | 0.55635302 |

| | | | | | | | | | | | |
|---|---|---|---|---|---|---|---|---|---|---|---|
| HSPA1A | P0DMV8 | 0.71936318 | NA | -0.3990722 | -1.5003485 | -2.1121375 | NA | NA | -1.0414324 | -1.8299469 | 0.78371139 |
| IL17C | Q9P0M4 | 0.67897824 | NA | NA | -0.4020892 | -0.5566759 | NA | NA | NA | NA | 0.69431103 |
| IL1RN | P18510 | 0.68271949 | NA | NA | -1.0185954 | -1.9076938 | NA | NA | -0.5988676 | -1.8515816 | 0.85301167 |
| IL5RA | Q01344 | 0.68932438 | -0.7591279 | -1.0588199 | -0.9011562 | -1.299383 | NA | NA | -0.5789765 | -1.0163204 | NA |
| IL24 | Q13007 | 0.65826743 | NA | NA | -0.5426438 | -0.7282679 | NA | NA | -0.4479044 | -0.5200959 | NA |
| IL6 | P05231 | 1.34731518 | 3.58683607 | 2.56774148 | -1.2838123 | -3.7308571 | NA | NA | -2.3349603 | -3.3569102 | 0.4194863 |
| ITGB6 | P18564 | 0.74270356 | 0.28961963 | 0.4059267 | 0.50015385 | NA | NA | NA | 0.47435294 | NA | 0.5107074 |
| JUN | P05412 | 0.77094689 | -0.2260119 | -0.5051074 | -1.3201246 | -1.6026571 | NA | -0.454314 | -1.0168647 | -1.4965673 | 0.27854455 |
| KRT19 | P08727 | 1.29876824 | NA | -0.7606193 | -2.7265677 | -3.3892652 | NA | -0.9101791 | -2.4860897 | -3.4281592 | 0.96242948 |
| LAIR1 | Q6GTX8 | 0.54750487 | NA | NA | NA | -0.5650313 | NA | NA | NA | NA | 0.72075612 |
| LILRB4 | Q8NHJ6 | 0.54320211 | NA | NA | -0.9817715 | -1.3009116 | NA | NA | -0.6094426 | -1.0418878 | 0.6832148 |
| LTBR | P36941 | 0.55263554 | NA | NA | NA | -0.4380295 | NA | NA | NA | NA | 0.58797985 |
| MAPK9 | P45984 | 0.67146926 | NA | -0.5709455 | -1.2921862 | -1.4553241 | NA | NA | -1.1135941 | -1.4149816 | 0.27625323 |
| MMP10 | P09238 | 0.72649429 | NA | NA | NA | -0.5820045 | NA | NA | NA | -0.4198449 | NA |
| MZB1 | Q8WU39 | 0.89023523 | NA | -0.2980517 | -0.9683738 | -1.2501161 | NA | NA | -0.4836353 | -1.1317959 | 0.73318885 |
| NBN | O60934 | 0.68325842 | -0.3359635 | NA | -0.7551515 | -1.3462839 | NA | NA | -0.5021206 | -1.0746755 | NA |
| NCF2 | P19878 | 0.78889333 | -0.3608046 | NA | NA | -0.490167 | NA | NA | NA | NA | NA |
| TNFRSF11B | O00300 | 0.72508944 | NA | -0.3271364 | -0.5936331 | -0.7413741 | NA | NA | -0.3639853 | -0.5506245 | 0.30337467 |
| OSM | P13725 | 0.75623855 | -0.365695 | NA | -0.4799485 | -1.1304295 | NA | NA | NA | -0.8176245 | 0.70926324 |
| PARP1 | P09874 | 0.56935359 | NA | NA | -0.5755154 | -0.8763321 | NA | NA | -0.5314441 | -0.5927184 | NA |
| PLAUR | Q03405 | 0.58357709 | NA | NA | -0.2323592 | -0.6089089 | NA | NA | NA | -0.3543776 | 0.87848038 |
| PREB | Q9HCU5 | 0.5525716 | NA | NA | -0.5317654 | -0.6601107 | NA | NA | -0.3476471 | -0.6735082 | 0.22975696 |
| PSIP1 | O75475 | 0.70923867 | NA | NA | -1.2444162 | -1.6902241 | NA | NA | -0.9700132 | -1.3154184 | 1.01443927 |
| TGFA | P01135 | 0.6932859 | NA | NA | NA | -0.5214393 | NA | 0.54109302 | NA | NA | 0.56950053 |
| PTPN6 | P29350 | 0.57220835 | -0.2755018 | NA | -0.4193592 | -1.0138304 | NA | NA | NA | -0.7279694 | 0.57202946 |
| TNFRSF11A | Q9Y6Q6 | 0.58464347 | NA | 0.38599205 | NA | -0.3727321 | NA | NA | NA | NA | 0.82583082 |
| SCGB1A1 | P11684 | 0.53046249 | 0.28572694 | 0.42333807 | 0.65242154 | 0.51792321 | NA | NA | 0.7254 | 0.48513878 | 0.80675431 |
| SCGB3A2 | Q96PL1 | 0.58253928 | NA | NA | -0.4671215 | -0.6877268 | NA | NA | NA | -0.5664286 | NA |
| SERPINB8 | P50452 | 0.56126803 | NA | -0.272829 | -0.9830954 | -1.4021036 | NA | NA | -0.8168838 | -1.1937612 | 0.74102709 |

| | | | | | | | | | | | |
|---|---|---|---|---|---|---|---|---|---|---|---|
| SIGLEC10 | Q96LC7 | 0.54729938 | NA | -0.2337239 | -0.6493462 | -1.0019134 | NA | NA | -0.37995 | -0.8762327 | 0.61600525 |
| SPON1 | Q9HCB6 | 0.69012832 | 0.31697397 | 0.35577614 | NA | NA | NA | NA | NA | NA | 0.74478866 |
| SRPK2 | P78362 | 0.71308181 | NA | NA | -0.3885169 | -0.8375304 | NA | NA | NA | -0.6890735 | 0.86785734 |
| TFF2 | Q03403 | 0.50451263 | 0.41818904 | 0.42332955 | 0.39169692 | NA | NA | NA | 0.56571912 | NA | 0.92943676 |
| ACVRL1 | P37023 | 0.55317538 | NA | NA | NA | NA | NA | NA | NA | NA | 0.56644526 |
| AGR2 | O95994 | 1.36763003 | NA | -0.3650869 | -2.0014208 | -2.2487643 | NA | -0.9518477 | -1.9578662 | -2.4480653 | 0.95540878 |
| ARID4B | Q4LE39 | 0.52268572 | -0.1752685 | -0.2351119 | -0.6170208 | -0.8964866 | NA | NA | -0.4592676 | -0.7007122 | NA |
| BAG3 | O95817 | 0.59841503 | NA | -0.3302903 | -0.8591892 | -0.6862339 | NA | NA | -0.8064529 | -0.5211286 | NA |
| BRK1 | Q8WUW1 | 0.894158 | NA | NA | -2.1508923 | -2.594342 | NA | NA | -1.9434162 | -2.2965551 | 0.63341962 |
| CCL19 | Q99731 | 0.72720067 | NA | -0.5412176 | -0.8788138 | -0.9513982 | NA | NA | NA | -0.726602 | 0.65238871 |
| CD177 | Q8N6Q3 | 0.78055655 | NA | NA | -0.3948185 | -1.0693241 | NA | NA | NA | -0.9635082 | 0.64899482 |
| CERT1 | Q9Y5P4 | 0.56060001 | -0.311489 | -0.228029 | -0.6695469 | -1.1716304 | NA | NA | -0.5160338 | -0.9399653 | 0.77857066 |
| CALCA | P01258 | 1.86079488 | -0.5121612 | -1.4141852 | -1.7892292 | -2.205817 | NA | NA | -1.2185721 | -1.9745939 | NA |
| CLPS | P04118 | 0.62528388 | 0.32420365 | 0.44590966 | -0.54078 | -1.1599696 | NA | NA | NA | -0.9088796 | NA |
| CXCL13 | O43927 | 0.6394359 | -0.2822881 | -0.4777483 | -0.8196531 | -1.2041384 | NA | NA | -0.6775485 | -0.9867306 | NA |
| DBI | P07108 | 0.51030067 | NA | NA | -0.5630138 | -0.9479491 | NA | NA | NA | -0.8289959 | 1.19059081 |
| DRAXIN | Q8NBI3 | 0.74957841 | 0.48229087 | NA | NA | NA | NA | NA | NA | NA | 0.81707625 |
| EFNA1 | P20827 | 0.52427156 | -0.4457753 | -0.5365028 | -0.5103192 | -0.7455777 | NA | NA | NA | -0.6306102 | 0.3949567 |
| EZR | P15311 | 0.51920514 | NA | -0.4156784 | -1.1713592 | -1.4008411 | NA | -0.537114 | -1.0961853 | -1.2250857 | 0.30978792 |
| FCRL5 | Q96RD9 | 0.53094027 | NA | -0.3597341 | -0.29229 | NA | NA | NA | NA | NA | NA |
| FGR | P09769 | 0.7591518 | -0.441026 | NA | -0.5452854 | -1.1158473 | NA | NA | NA | -1.0092327 | NA |
| FMNL1 | O95466 | 0.70744409 | NA | NA | -0.3660046 | -0.8977027 | NA | NA | NA | -0.6175612 | 0.60276407 |
| GPKOW | Q92917 | 0.58768065 | NA | NA | -0.9374385 | -1.3487545 | NA | NA | -0.8129809 | -1.2097408 | 0.33474784 |
| HMOX2 | P30519 | 0.56874776 | NA | NA | -0.7219569 | -1.0908777 | NA | NA | -0.4974515 | -0.9712388 | 0.49341435 |
| IDI2 | Q9BXS1 | 0.75720271 | NA | NA | -0.6041854 | -0.6948464 | NA | NA | -0.5217559 | -0.8778122 | NA |
| IGFBP4 | P22692 | 0.87673384 | NA | NA | NA | NA | NA | NA | NA | NA | 0.87817439 |
| ILKAP | Q9H0C8 | 0.52141296 | NA | NA | -0.8325692 | -1.2252911 | NA | NA | -0.5591515 | -1.0053694 | 0.46606376 |
| ING1 | Q9UK53 | 0.54889852 | NA | -0.2096727 | -0.8038623 | -1.0898563 | NA | NA | -0.7418103 | -1.0366143 | 0.34109222 |
| CXCL8 | P10145 | 0.66938586 | 0.30331963 | NA | -0.7673215 | -1.1848464 | NA | NA | NA | -1.1059592 | 0.96662538 |

| | | | | | | | | | | | |
|---|---|---|---|---|---|---|---|---|---|---|---|
| IPCEF1 | Q8WWN9 | 0.53010052 | -0.4868639 | NA | NA | -0.8389268 | NA | NA | NA | NA | 0.80193478 |
| LAYN | Q6UX15 | 0.51985268 | 0.26431142 | 0.29265057 | NA | NA | NA | NA | NA | NA | 0.7383946 |
| LBR | Q14739 | 0.68594354 | NA | NA | -0.8643554 | -1.425617 | NA | NA | -0.5757559 | -1.3177714 | NA |
| LIF | P15018 | 0.89416345 | 0.51700776 | NA | -0.7281715 | -1.0413821 | NA | NA | -0.5114426 | -0.9686265 | NA |
| LPO | P22079 | 0.55919389 | NA | NA | -0.6142685 | -0.8153741 | NA | NA | -0.7212809 | -0.9174429 | NA |
| LRPAP1 | P30533 | 0.61934545 | NA | NA | -0.56619 | -0.7567795 | NA | NA | -0.3660882 | -0.6782918 | NA |
| MAD1L1 | Q9Y6D9 | 0.88536232 | NA | -0.3363892 | -1.6646938 | -2.3085598 | NA | NA | -1.1811941 | -2.0001082 | 0.60134944 |
| MATN3 | O15232 | 0.50238467 | -0.4147498 | -0.8799517 | -1.3224523 | -1.7412286 | NA | -0.4420477 | -1.1275206 | -1.5246184 | NA |
| MFGE8 | Q08431 | 0.5163751 | 0.27036667 | 0.39010625 | NA | -0.3981089 | NA | NA | NA | NA | NA |
| MMP8 | P22894 | 0.87328515 | -0.3935973 | NA | -0.7001146 | -1.7437536 | NA | NA | NA | -1.3449286 | 0.93528935 |
| MMP9 | P14780 | 0.51936281 | -0.2196662 | NA | NA | -0.3280821 | NA | NA | NA | NA | 0.40634789 |
| MUC13 | Q9H3R2 | 0.63431747 | NA | NA | NA | -0.2950054 | NA | NA | NA | NA | 0.44108148 |
| NEFL | P07196 | 0.8544322 | NA | 0.85582727 | 0.66456077 | NA | NA | 0.83050349 | 1.49449118 | NA | 1.32291301 |
| NMNAT1 | Q9HAN9 | 0.80236291 | NA | NA | NA | -0.5873384 | NA | NA | NA | NA | NA |
| NOS1 | P29475 | 1.14826236 | NA | NA | -0.9740631 | -1.4022598 | NA | NA | -0.7241441 | -1.5019224 | NA |
| NPM1 | P06748 | 0.63069904 | -0.2554712 | -0.2698705 | -1.1268138 | -1.647567 | NA | NA | -0.8023412 | -1.3405265 | NA |
| OXT | P01178 | 0.65491087 | NA | NA | -0.5085462 | NA | NA | NA | NA | NA | NA |
| PADI4 | Q9UM07 | 0.72744124 | NA | NA | NA | -0.7789902 | NA | 0.6784814 | NA | NA | 0.84176085 |
| PAEP | P09466 | 0.60473301 | NA | NA | NA | NA | NA | NA | NA | NA | NA |
| PRL | P01236 | 0.59604587 | 0.36586027 | 0.43572386 | NA | -0.4448661 | NA | NA | NA | NA | NA |
| SPINK1 | P00995 | 1.18341093 | -0.5397333 | -0.776358 | -1.0198562 | -1.4255241 | NA | NA | -0.6189088 | -1.2369918 | 0.58193447 |
| PTK7 | Q13308 | 0.50476252 | 0.22310548 | 0.51784318 | 0.40233692 | NA | NA | 0.48784302 | 0.7309 | NA | 0.53506646 |
| PXN | P49023 | 0.72450854 | -0.3346466 | NA | -0.3572908 | -0.918558 | NA | NA | NA | -0.7289429 | 0.69594669 |
| RELT | Q969Z4 | 0.58037958 | NA | NA | NA | -0.2553598 | NA | NA | NA | NA | 0.4073837 |
| RSPO1 | Q2MKA7 | 1.11017417 | NA | NA | -0.7280585 | -0.8558634 | NA | NA | NA | -0.6140612 | 0.29438742 |
| SCARB2 | Q14108 | 0.51463816 | NA | NA | NA | -0.6040402 | NA | NA | NA | NA | 0.69726476 |
| SERPINB1 | P30740 | 0.73432531 | NA | NA | -0.8780092 | -1.5668696 | NA | NA | -0.5151162 | -1.2446551 | 0.5518 |
| SFRP1 | Q8N474 | 1.50029863 | NA | NA | -1.6962077 | -2.518767 | NA | NA | -1.3945632 | -2.3142735 | 1.18900196 |
| SNCG | O76070 | 0.61729998 | NA | NA | NA | -0.3723116 | NA | NA | NA | NA | 0.6050089 |

| | | | | | | | | | | | |
|---|---|---|---|---|---|---|---|---|---|---|---|
| STC1 | P52823 | 0.64259414 | NA | NA | -0.7430508 | -0.9221045 | NA | NA | -0.4827015 | -0.6486571 | 0.34253738 |
| TDGF1 | P13385 | 0.56944026 | NA | NA | NA | NA | NA | NA | NA | NA | NA |
| TFF1 | P04155 | 0.73709758 | 0.50211553 | 0.69160227 | 0.53261923 | NA | NA | NA | 0.75078529 | NA | 1.25316171 |
| THBS2 | P35442 | 0.75011575 | NA | NA | -0.31982 | -0.7069009 | NA | NA | NA | -0.4527694 | NA |
| TMSB10 | P63313 | 0.71424851 | NA | NA | -0.9165792 | -1.2971759 | NA | NA | -0.65715 | -1.1591204 | 1.09354058 |
| TNFRSF10A | O00220 | 0.55676992 | -0.1963237 | -0.4385375 | -0.8233569 | -1.1295554 | NA | NA | -0.4824147 | -0.8984612 | 0.40963349 |
| TNFRSF10B | O14763 | 1.2218294 | NA | NA | -0.7772862 | -1.3070598 | NA | NA | NA | -0.9637816 | 0.93528372 |
| TNFRSF1A | P19438 | 0.64110734 | NA | NA | -0.4977662 | -0.8992929 | NA | NA | NA | -0.7195082 | 0.63709711 |
| TNFRSF6B | O95407 | 1.02099301 | 0.43388082 | NA | -0.8874246 | -1.2654741 | NA | NA | -0.5198397 | -1.0245735 | 0.82264626 |
| TXLNA | P40222 | 0.51464007 | NA | NA | -1.0273315 | -1.5770795 | NA | NA | -0.758425 | -1.3010837 | 0.90281175 |
| TXNDC5 | Q8NBS9 | 0.64419442 | NA | NA | -0.7044892 | -0.9264911 | NA | NA | -0.4188294 | -0.9109735 | 0.60807355 |
| ULBP2 | Q9BZM5 | 0.79212916 | NA | NA | NA | -0.5496634 | NA | NA | NA | NA | 0.70605483 |
| VSIG4 | Q9Y279 | 0.78785622 | NA | -0.6408943 | -1.6392454 | -2.1875714 | NA | NA | -1.1594221 | -2.0524755 | 0.64936909 |
| ADM | P35318 | 0.65402504 | NA | -0.2790722 | -0.5330408 | -0.6561768 | NA | NA | NA | -0.6505 | NA |
| APBB1IP | Q7Z5R6 | 0.73770881 | -0.2139114 | NA | -0.5275131 | -1.0617946 | NA | NA | NA | -0.7493837 | 0.92788577 |
| AREG | P15514 | 1.11847818 | NA | -0.3803426 | -1.6161092 | -2.1301696 | NA | NA | -1.3308456 | -2.0899163 | 0.54918911 |
| ARHGAP25 | P42331 | 0.61254305 | -0.3511374 | NA | NA | -0.7017554 | NA | NA | NA | NA | 0.87025175 |
| BGN | P21810 | 0.88359171 | 0.48848174 | 0.39714489 | -0.4874569 | -0.7172438 | NA | NA | NA | -0.7276429 | NA |
| MUC16 | Q8WXI7 | 0.6396812 | NA | 0.58317614 | 0.79408538 | NA | NA | NA | 0.77132206 | NA | NA |
| CALB1 | P05937 | 1.25522407 | NA | NA | -0.3946969 | -0.8157116 | NA | NA | NA | -0.9671 | NA |
| CD300E | Q496F6 | 0.5050925 | -0.3879694 | -0.6910102 | -1.0829646 | -1.3330027 | NA | NA | -0.6283559 | -1.051751 | 0.81092041 |
| CD302 | Q8IX05 | 0.5060034 | NA | 0.40314886 | NA | NA | NA | NA | NA | NA | 0.57551841 |
| CEACAM5 | P06731 | 0.98795378 | 0.27727032 | 0.38698466 | -0.2735231 | -0.7872063 | NA | NA | NA | -0.6937184 | 0.81491969 |
| COX5B | P10606 | 0.61941762 | NA | NA | -0.3925169 | -0.7905196 | NA | NA | NA | -0.6807469 | NA |
| KRT18 | P05783 | 0.85509489 | 0.63927078 | NA | -1.4439823 | -1.8392723 | NA | -0.5722512 | -1.5889426 | -1.9535735 | 0.99659293 |
| DPY30 | Q9C005 | 1.34145848 | NA | -0.478046 | -2.4004446 | -3.1426455 | NA | -0.8735395 | -2.1333971 | -2.8801633 | NA |
| EDA2R | Q9HAV5 | 0.72597939 | 0.32823562 | 0.60440057 | 0.48573615 | NA | NA | 0.6615686 | 0.83599412 | NA | 1.38208962 |
| S100A12 | P80511 | 0.73405994 | NA | NA | -0.3178723 | -0.8864473 | NA | NA | NA | -0.5345735 | 0.77449241 |
| EPS8L2 | Q9H6S3 | 0.61185913 | 0.42433379 | 0.32072159 | -0.5983023 | -0.7443125 | NA | NA | -0.6344956 | -0.7828143 | 0.41887259 |

| | | | | | | | | | | | |
|---|---|---|---|---|---|---|---|---|---|---|---|
| F3 | P13726 | 0.76943182 | NA | NA | -0.5666315 | -0.5528554 | NA | NA | -0.7847309 | -0.592798 | NA |
| FEN1 | P39748 | 0.72022239 | -0.3241607 | NA | -0.3966785 | -1.059683 | NA | NA | NA | -0.8630102 | 1.17857331 |
| FGF21 | Q9NSA1 | 0.85773904 | NA | -0.5777937 | NA | NA | NA | NA | NA | NA | 1.36483937 |
| FGF23 | Q9GZV9 | 1.13128154 | 0.56067945 | NA | NA | NA | NA | NA | NA | NA | 0.62565989 |
| FGFBP1 | Q14512 | 1.45407909 | 0.65700137 | 0.9003392 | -0.6491338 | -1.4245893 | NA | NA | NA | -1.1045163 | NA |
| FUS | P35637 | 0.52649609 | NA | NA | -0.4925585 | -0.890008 | NA | NA | -0.4117721 | -0.7932551 | 0.39669733 |
| WFDC2 | Q14508 | 1.11359034 | NA | 0.30017159 | -0.3866023 | -1.1229357 | NA | 0.46101512 | NA | -0.8051878 | 1.18322683 |
| HS3ST3B1 | Q9Y662 | 0.50794368 | NA | -0.2956864 | -1.0564238 | -1.2573152 | NA | NA | -0.6302603 | -0.9134755 | NA |
| HS6ST1 | O60243 | 0.62399385 | -0.4202324 | -0.416108 | -0.8451777 | -1.1710018 | NA | NA | -0.6600779 | -0.786551 | 1.01183433 |
| HSPB6 | O14558 | 0.63247372 | NA | NA | NA | NA | NA | NA | NA | NA | 0.21745106 |
| IGSF3 | O75054 | 0.7685517 | -0.9566548 | -1.1955807 | -1.0241485 | -1.0822634 | NA | NA | -0.7053824 | -0.9359041 | 0.36770759 |
| HAVCR1 | Q96D42 | 0.69077251 | NA | 0.3364733 | NA | NA | NA | NA | NA | NA | 1.32924051 |
| MDK | P21741 | 1.34818099 | NA | NA | -1.7835946 | -2.5319893 | NA | NA | -1.3108294 | -2.1018816 | 0.74069515 |
| MMP12 | P39900 | 0.51277931 | NA | NA | 0.71381692 | 0.90118929 | NA | NA | 0.75156029 | 0.92394082 | 0.7631756 |
| MSLN | Q13421 | 0.72717531 | 0.35778995 | 0.53096477 | NA | -0.610458 | NA | 0.48549884 | 0.41691176 | NA | 1.60659651 |
| NDUFS6 | O75380 | 1.00880903 | NA | NA | -0.8712254 | -1.2985714 | NA | NA | -0.5719662 | -1.3265551 | 0.35751726 |
| PFKFB2 | O60825 | 0.92430638 | -0.3967594 | NA | -0.4422531 | -1.1133241 | NA | NA | NA | -0.8262878 | 0.45700291 |
| POLR2F | P61218 | 0.87032316 | NA | -0.2313153 | -1.0164815 | -1.4863393 | NA | NA | -0.7395074 | -1.3060184 | 0.54003416 |
| PQBP1 | O60828 | 0.71168809 | NA | NA | -1.0240085 | -1.4550848 | NA | NA | -1.0873441 | -1.3626143 | NA |
| RASSF2 | P50749 | 0.8725256 | -0.4467539 | NA | NA | -0.7806134 | NA | NA | NA | NA | 0.88719957 |
| RRM2 | P31350 | 0.75463791 | NA | -0.8326318 | -2.8942862 | -3.8360821 | NA | -0.9136547 | -2.5499 | -3.3872143 | 0.56681781 |
| RSPO3 | Q9BXY4 | 0.79220248 | NA | NA | -0.5662192 | -0.6544696 | NA | NA | NA | -0.4768551 | 0.63778623 |
| SFTPA1 | Q8IWL2 | 0.61555134 | NA | NA | -1.1024123 | -1.4633527 | NA | NA | -1.0040074 | -1.3490592 | -0.3069958 |
| SFTPA2 | Q8IWL1 | 0.52209612 | NA | 0.35824432 | -0.2667308 | -0.8798563 | NA | NA | NA | -0.8187245 | 0.40873488 |
| SMOC1 | Q9H4F8 | 0.59872866 | NA | NA | -1.4726315 | -2.0270768 | NA | NA | -1.0627765 | -1.7535714 | 1.23925333 |
| SRP14 | P37108 | 0.52073902 | -0.3733986 | NA | -0.3130015 | -0.7413973 | NA | NA | NA | -0.4268939 | 0.78014087 |
| TACSTD2 | P09758 | 0.72394645 | NA | NA | -0.6035285 | -0.6926964 | NA | -0.3977767 | -0.6206735 | -0.5932551 | 0.2612566 |
| TFPI2 | P48307 | 0.72232754 | NA | NA | -0.8239485 | -1.1650259 | NA | NA | NA | -0.8743694 | 0.40199475 |
| TNFRSF12A | Q9NP84 | 1.06307458 | NA | NA | NA | -0.4832277 | NA | NA | NA | NA | 0.45050597 |

| | | | | | | | | | | | |
|---|---|---|---|---|---|---|---|---|---|---|---|
| TRIAP1 | O43715 | 0.83714346 | NA | NA | -1.1883662 | -1.6262063 | NA | NA | -1.0839059 | -1.506551 | 0.63921977 |
| FLT1 | P17948 | 0.61864805 | NA | NA | -0.4158015 | -0.6803679 | NA | NA | NA | -0.5229571 | NA |
| CCN4 | O95388 | 0.65310575 | NA | NA | NA | NA | NA | NA | NA | NA | 0.71219162 |

**Supplemental Table 2. Hazard modeling of serum proteomic data for length of hospital stay and clinical failure.**

Each row represents a protein significantly associated with the clinical endpoints “time to hospital discharge” and “clinical failure.” The outputs include hazard ratios with lower and upper 95% confidence intervals, unadjusted and Benjamini-Hochberg adjusted *P* values.

HR, hazard ratio.

| OLINK Prognostic Model Output for Time to Hospital Discharge | | | | | |
|---|---|---|---|---|---|
| **Protein** | **HR** | **Lower** | **Upper** | ***P* value** | ***P* adjusted** |
| ADA2 | 1.25 | 1.09 | 1.43 | 0.0016439 | 0.013749 |
| AHCY | 0.8 | 0.69 | 0.93 | 0.00339067 | 0.02332275 |
| ANGPTL1 | 0.71 | 0.62 | 0.81 | 6.596E-07 | 4.2215E-05 |
| ACTA2 | 0.72 | 0.6 | 0.87 | 0.0006236 | 0.00646439 |
| TNFSF13B | 0.76 | 0.66 | 0.87 | 0.00011036 | 0.00202492 |
| DIABLO | 0.68 | 0.58 | 0.79 | 9.7042E-07 | 5.0745E-05 |
| CA3 | 0.76 | 0.65 | 0.88 | 0.00039291 | 0.00477986 |
| CCDC80 | 0.77 | 0.65 | 0.92 | 0.00350212 | 0.02386628 |
| CCL5 | 1.22 | 1.05 | 1.42 | 0.00853722 | 0.04259926 |
| CD14 | 0.81 | 0.72 | 0.92 | 0.00106138 | 0.00976469 |
| CD209 | 0.82 | 0.7 | 0.95 | 0.0081481 | 0.04093517 |
| CD93 | 0.8 | 0.69 | 0.94 | 0.00678238 | 0.03565595 |
| CLC | 1.27 | 1.09 | 1.47 | 0.00222443 | 0.01732465 |
| CLEC5A | 0.72 | 0.6 | 0.87 | 0.0004778 | 0.00548145 |
| CLTA | 0.67 | 0.57 | 0.78 | 7.4055E-07 | 4.2775E-05 |
| COL4A1 | 0.74 | 0.63 | 0.87 | 0.00034466 | 0.00452976 |
| CSTB | 0.74 | 0.63 | 0.87 | 0.00018709 | 0.00281016 |
| CTSB | 0.77 | 0.64 | 0.92 | 0.0045261 | 0.02789798 |
| CTSL | 0.76 | 0.64 | 0.9 | 0.00172724 | 0.01436438 |
| CST3 | 0.78 | 0.67 | 0.92 | 0.00282409 | 0.02008239 |
| DCN | 0.65 | 0.55 | 0.77 | 8.0888E-07 | 4.4099E-05 |
| DEFA1B | 0.8 | 0.68 | 0.95 | 0.00894145 | 0.04431589 |
| DEFA1 | 0.8 | 0.68 | 0.95 | 0.00894145 | 0.04431589 |
| DKK3 | 0.68 | 0.58 | 0.8 | 3.7874E-06 | 0.00014604 |
| EPHB4 | 0.75 | 0.64 | 0.89 | 0.00085605 | 0.00812967 |

| | | | | | |
|---|---|---|---|---|---|
| FABP2 | 0.76 | 0.65 | 0.9 | 0.00135043 | 0.0117623 |
| FABP6 | 0.78 | 0.66 | 0.92 | 0.00290364 | 0.02045052 |
| F9 | 1.27 | 1.09 | 1.48 | 0.00209216 | 0.01673728 |
| FAP | 0.82 | 0.71 | 0.94 | 0.00570934 | 0.03183316 |
| FETUB | 1.41 | 1.16 | 1.71 | 0.0005033 | 0.00564419 |
| GAS6 | 0.75 | 0.65 | 0.86 | 5.9075E-05 | 0.00122986 |
| GDF15 | 0.71 | 0.59 | 0.86 | 0.00033386 | 0.00442736 |
| IL6ST | 0.8 | 0.68 | 0.92 | 0.00284361 | 0.02012402 |
| GPR37 | 0.8 | 0.68 | 0.93 | 0.00317821 | 0.02206759 |
| HMOX1 | 0.82 | 0.7 | 0.95 | 0.00900099 | 0.04446127 |
| IGFBP1 | 0.67 | 0.57 | 0.79 | 7.5553E-07 | 4.2775E-05 |
| IL1RL1 | 0.56 | 0.47 | 0.67 | 1.5119E-10 | 1.1127E-07 |
| IL2RA | 0.78 | 0.67 | 0.92 | 0.00236359 | 0.01802336 |
| IL18BP | 0.77 | 0.67 | 0.9 | 0.00075083 | 0.00741761 |
| LEPR | 0.78 | 0.68 | 0.9 | 0.00068464 | 0.00699857 |
| LGALS1 | 0.73 | 0.62 | 0.87 | 0.00027019 | 0.00386893 |
| LILRA5 | 0.7 | 0.6 | 0.81 | 2.6769E-06 | 0.00011258 |
| LTBP2 | 0.6 | 0.5 | 0.71 | 1.7893E-09 | 5.2677E-07 |
| MARCO | 1.21 | 1.05 | 1.4 | 0.00931683 | 0.04541183 |
| MCFD2 | 0.79 | 0.68 | 0.91 | 0.00150635 | 0.01281704 |
| MB | 0.63 | 0.53 | 0.76 | 6.2624E-07 | 4.1901E-05 |
| NADK | 0.78 | 0.68 | 0.9 | 0.00060363 | 0.00630168 |
| NID1 | 0.82 | 0.71 | 0.95 | 0.00632455 | 0.034227 |
| SPP1 | 0.83 | 0.72 | 0.96 | 0.00983911 | 0.04740958 |
| OSMR | 0.74 | 0.64 | 0.86 | 4.9619E-05 | 0.00109013 |
| PAG1 | 0.73 | 0.63 | 0.85 | 7.108E-05 | 0.00139506 |
| PDGFA | 1.35 | 1.13 | 1.61 | 0.00073042 | 0.00726826 |
| PDGFRA | 0.66 | 0.56 | 0.78 | 7.3489E-07 | 4.2775E-05 |
| PI3 | 0.76 | 0.64 | 0.91 | 0.00312463 | 0.02179837 |
| PLA2G2A | 0.8 | 0.69 | 0.92 | 0.00211591 | 0.01683575 |
| PLIN3 | 0.76 | 0.65 | 0.88 | 0.00038922 | 0.00477986 |
| PLXNB2 | 0.8 | 0.7 | 0.93 | 0.00246134 | 0.01848518 |
| PPP1R2 | 0.79 | 0.67 | 0.92 | 0.00245356 | 0.01848518 |
| RCOR1 | 0.72 | 0.6 | 0.87 | 0.00055998 | 0.00593012 |
| REG1A | 0.77 | 0.64 | 0.92 | 0.00492614 | 0.02898673 |
| REG1B | 0.76 | 0.64 | 0.91 | 0.00214687 | 0.01689942 |
| RETN | 0.8 | 0.68 | 0.94 | 0.0064013 | 0.03438948 |
| S100A11 | 0.79 | 0.67 | 0.92 | 0.00294022 | 0.02060957 |
| SDC4 | 1.24 | 1.06 | 1.45 | 0.00622005 | 0.03402017 |
| SEMA3F | 0.8 | 0.69 | 0.94 | 0.00524864 | 0.03006226 |

| | | | | | |
|---|---|---|---|---|---|
| SPARCL1 | 0.82 | 0.71 | 0.95 | 0.00807214 | 0.04083225 |
| SSC4D | 1.24 | 1.07 | 1.43 | 0.00365437 | 0.02449695 |
| THBD | 0.76 | 0.64 | 0.91 | 0.00260672 | 0.01918545 |
| THBS4 | 0.79 | 0.69 | 0.92 | 0.00221798 | 0.01732465 |
| TINAGL1 | 0.7 | 0.6 | 0.81 | 3.8693E-06 | 0.00014604 |
| TNC | 0.79 | 0.68 | 0.92 | 0.00274484 | 0.01984493 |
| PLAT | 0.8 | 0.69 | 0.93 | 0.00275025 | 0.01984493 |
| TNNI3 | 0.72 | 0.61 | 0.86 | 0.00015584 | 0.00246662 |
| VCAM1 | 0.74 | 0.64 | 0.86 | 6.7923E-05 | 0.00135112 |
| CHI3L1 | 0.79 | 0.66 | 0.94 | 0.00703632 | 0.03634196 |
| ADGRE2 | 0.81 | 0.7 | 0.95 | 0.00702527 | 0.03634196 |
| AGRN | 0.75 | 0.63 | 0.89 | 0.00087647 | 0.00827024 |
| AMBN | 1.3 | 1.15 | 1.46 | 1.5431E-05 | 0.00042859 |
| ANGPTL4 | 0.67 | 0.57 | 0.79 | 1.8807E-06 | 8.6511E-05 |
| TNFSF13 | 0.75 | 0.64 | 0.88 | 0.0003755 | 0.00472428 |
| ARNT | 1.21 | 1.05 | 1.4 | 0.00700645 | 0.03634196 |
| ATP5IF1 | 0.64 | 0.54 | 0.75 | 4.1935E-08 | 4.9709E-06 |
| B4GALT1 | 0.82 | 0.71 | 0.95 | 0.00955922 | 0.04643951 |
| CCL20 | 0.76 | 0.64 | 0.91 | 0.0023585 | 0.01802336 |
| CCL7 | 0.78 | 0.67 | 0.9 | 0.00054881 | 0.00592576 |
| CD200R1 | 1.23 | 1.05 | 1.43 | 0.0093057 | 0.04541183 |
| CD276 | 0.82 | 0.7 | 0.95 | 0.00978861 | 0.04739747 |
| CD40LG | 1.3 | 1.09 | 1.54 | 0.00373525 | 0.02465599 |
| CD84 | 1.27 | 1.07 | 1.5 | 0.00653041 | 0.03495551 |
| CDSN | 0.78 | 0.66 | 0.93 | 0.004894 | 0.02898673 |
| CHRDL1 | 0.79 | 0.67 | 0.94 | 0.00599511 | 0.03292839 |
| CKAP4 | 0.6 | 0.5 | 0.72 | 3.219E-08 | 4.3076E-06 |
| CRELD2 | 0.74 | 0.62 | 0.88 | 0.00075892 | 0.00744758 |
| CRIM1 | 0.73 | 0.63 | 0.84 | 2.0126E-05 | 0.00054863 |
| CRLF1 | 0.74 | 0.64 | 0.86 | 5.3778E-05 | 0.00114726 |
| CTSC | 0.76 | 0.65 | 0.88 | 0.00025893 | 0.00381152 |
| CTSO | 0.78 | 0.68 | 0.89 | 0.00029372 | 0.00404068 |
| CXCL10 | 0.78 | 0.69 | 0.89 | 0.00015014 | 0.00241564 |
| CXCL12 | 1.23 | 1.06 | 1.41 | 0.00531086 | 0.03024004 |
| LAMP3 | 0.81 | 0.72 | 0.91 | 0.00048037 | 0.00548145 |
| DECR1 | 0.81 | 0.7 | 0.94 | 0.00443688 | 0.02767408 |
| DFFA | 0.75 | 0.64 | 0.88 | 0.00042564 | 0.00501232 |
| EGF | 1.38 | 1.16 | 1.64 | 0.00027072 | 0.00386893 |
| ENAH | 0.75 | 0.64 | 0.88 | 0.00055764 | 0.00593012 |
| EPCAM | 0.78 | 0.68 | 0.91 | 0.0014584 | 0.01248116 |

| | | | | | |
|---|---|---|---|---|---|
| ESM1 | 0.77 | 0.67 | 0.9 | 0.0005664 | 0.00595527 |
| FABP9 | 0.82 | 0.71 | 0.95 | 0.00867206 | 0.04312592 |
| FLT3LG | 1.23 | 1.05 | 1.44 | 0.01028924 | 0.04873455 |
| FST | 0.71 | 0.6 | 0.83 | 4.0784E-05 | 0.00093802 |
| FSTL3 | 0.63 | 0.52 | 0.77 | 4.4692E-06 | 0.00016046 |
| GBP2 | 0.81 | 0.7 | 0.94 | 0.00494271 | 0.02898673 |
| CSF3 | 0.8 | 0.69 | 0.94 | 0.00563038 | 0.03161946 |
| GLOD4 | 0.79 | 0.68 | 0.92 | 0.00176075 | 0.0145608 |
| HEXIM1 | 0.67 | 0.58 | 0.78 | 4.7837E-07 | 3.5541E-05 |
| HSD11B1 | 0.82 | 0.72 | 0.95 | 0.00625675 | 0.03402017 |
| HSPA1A | 0.75 | 0.64 | 0.88 | 0.00031243 | 0.00421925 |
| IFNG | 0.83 | 0.72 | 0.95 | 0.00758782 | 0.03878218 |
| IFNGR1 | 0.74 | 0.62 | 0.88 | 0.00071462 | 0.00720497 |
| IL18R1 | 0.74 | 0.64 | 0.86 | 7.7313E-05 | 0.00149743 |
| IL1RN | 0.74 | 0.64 | 0.86 | 6.5008E-05 | 0.00131086 |
| IL4R | 0.79 | 0.68 | 0.93 | 0.00416302 | 0.02641363 |
| IL13 | 1.19 | 1.06 | 1.33 | 0.00254047 | 0.01888673 |
| IL15 | 0.73 | 0.63 | 0.85 | 4.4141E-05 | 0.00098448 |
| IL4 | 1.21 | 1.05 | 1.4 | 0.00919444 | 0.04526494 |
| IL6 | 0.7 | 0.6 | 0.83 | 3.3101E-05 | 0.00079876 |
| ITGB6 | 0.7 | 0.58 | 0.84 | 0.00012334 | 0.00213598 |
| JUN | 0.68 | 0.58 | 0.8 | 3.7174E-06 | 0.00014604 |
| KRT19 | 0.72 | 0.63 | 0.82 | 1.2257E-06 | 6.014E-05 |
| LAP3 | 0.78 | 0.68 | 0.9 | 0.00077788 | 0.00758307 |
| LGALS9 | 0.71 | 0.61 | 0.83 | 9.2845E-06 | 0.00027891 |
| LIFR | 0.8 | 0.7 | 0.92 | 0.00152784 | 0.01292518 |
| LILRB4 | 0.8 | 0.68 | 0.93 | 0.00448797 | 0.02787465 |
| LTBR | 0.74 | 0.62 | 0.89 | 0.0009429 | 0.00884048 |
| MAPK9 | 0.69 | 0.59 | 0.8 | 9.9973E-07 | 5.0745E-05 |
| MATN2 | 0.75 | 0.64 | 0.88 | 0.00050614 | 0.00564419 |
| CSF1 | 0.79 | 0.69 | 0.9 | 0.00054866 | 0.00592576 |
| METAP1D | 0.81 | 0.69 | 0.94 | 0.00664169 | 0.03520125 |
| MPIG6B | 1.23 | 1.06 | 1.42 | 0.00583026 | 0.03226371 |
| NUB1 | 0.8 | 0.69 | 0.93 | 0.00376065 | 0.02471286 |
| NUDC | 0.81 | 0.7 | 0.94 | 0.00475915 | 0.02859375 |
| OMD | 0.81 | 0.72 | 0.92 | 0.00144371 | 0.01248116 |
| TNFRSF11B | 0.68 | 0.56 | 0.83 | 0.00011816 | 0.002101 |
| PADI2 | 0.78 | 0.65 | 0.93 | 0.00670115 | 0.03535516 |
| PARP1 | 0.73 | 0.61 | 0.88 | 0.00107066 | 0.0097889 |
| PDGFB | 1.28 | 1.08 | 1.52 | 0.00493916 | 0.02898673 |

| | | | | | |
|---|---|---|---|---|---|
| PLAUR | 0.75 | 0.62 | 0.9 | 0.0018924 | 0.01539015 |
| PGF | 0.77 | 0.64 | 0.92 | 0.00426326 | 0.02693355 |
| PRDX3 | 0.79 | 0.69 | 0.91 | 0.00145532 | 0.01248116 |
| PREB | 0.6 | 0.49 | 0.73 | 2.8764E-07 | 2.3522E-05 |
| PRELP | 0.78 | 0.67 | 0.9 | 0.00080949 | 0.00773748 |
| PROK1 | 0.71 | 0.61 | 0.82 | 6.6926E-06 | 0.00022911 |
| PSIP1 | 0.76 | 0.65 | 0.88 | 0.00018147 | 0.00275385 |
| PTX3 | 0.8 | 0.7 | 0.92 | 0.0013012 | 0.011401 |
| AGER | 0.72 | 0.63 | 0.83 | 4.2379E-06 | 0.00015595 |
| SCRN1 | 0.81 | 0.7 | 0.94 | 0.00502756 | 0.0292466 |
| SERPINB8 | 0.77 | 0.67 | 0.89 | 0.0003743 | 0.00472428 |
| SIGLEC1 | 0.78 | 0.69 | 0.89 | 0.00028418 | 0.00394641 |
| SRPK2 | 0.81 | 0.69 | 0.95 | 0.00988773 | 0.04740958 |
| TRIM21 | 0.84 | 0.74 | 0.96 | 0.00988146 | 0.04740958 |
| ACVRL1 | 0.75 | 0.62 | 0.92 | 0.00461697 | 0.02815318 |
| AGR2 | 0.69 | 0.59 | 0.79 | 5.0704E-07 | 3.5541E-05 |
| ANXA5 | 0.7 | 0.58 | 0.83 | 5.9321E-05 | 0.00122986 |
| APP | 1.36 | 1.14 | 1.63 | 0.00051329 | 0.00568089 |
| ARID4B | 0.74 | 0.62 | 0.87 | 0.00033249 | 0.00442736 |
| BAG3 | 0.74 | 0.64 | 0.84 | 1.0509E-05 | 0.00029749 |
| BCAM | 0.77 | 0.66 | 0.9 | 0.00079092 | 0.00765948 |
| CD300C | 0.81 | 0.7 | 0.95 | 0.00838219 | 0.04196796 |
| CD74 | 0.82 | 0.71 | 0.94 | 0.00554974 | 0.03129969 |
| CDHR1 | 1.36 | 1.2 | 1.55 | 2.3076E-06 | 9.9905E-05 |
| CALCA | 0.64 | 0.54 | 0.76 | 2.6131E-07 | 2.2627E-05 |
| CLEC11A | 0.81 | 0.7 | 0.94 | 0.00471948 | 0.02856524 |
| CLEC1B | 1.22 | 1.06 | 1.4 | 0.0056494 | 0.03161946 |
| CLSTN1 | 1.19 | 1.05 | 1.36 | 0.0059504 | 0.0328052 |
| CRIP2 | 0.77 | 0.67 | 0.88 | 0.00017933 | 0.00274977 |
| CX3CL1 | 0.68 | 0.59 | 0.8 | 1.3716E-06 | 6.5127E-05 |
| EFNA1 | 0.69 | 0.57 | 0.83 | 7.9776E-05 | 0.00152506 |
| ENO1 | 0.8 | 0.7 | 0.92 | 0.00223743 | 0.01733421 |
| EREG | 1.23 | 1.07 | 1.41 | 0.00364504 | 0.02449695 |
| EZR | 0.63 | 0.54 | 0.73 | 1.3213E-09 | 4.8623E-07 |
| FABP5 | 0.8 | 0.68 | 0.92 | 0.00274053 | 0.01984493 |
| FCRL5 | 1.25 | 1.07 | 1.45 | 0.00404823 | 0.02579652 |
| FGR | 0.81 | 0.69 | 0.94 | 0.00730695 | 0.03760779 |
| FKBP4 | 0.82 | 0.72 | 0.94 | 0.00462844 | 0.02815318 |
| FOSB | 0.74 | 0.63 | 0.86 | 0.00014841 | 0.00241564 |
| GP6 | 1.22 | 1.07 | 1.4 | 0.0037072 | 0.02460801 |

| | | | | | |
|---|---|---|---|---|---|
| GPC5 | 1.24 | 1.06 | 1.44 | 0.00664806 | 0.03520125 |
| GPKOW | 0.7 | 0.6 | 0.83 | 2.8535E-05 | 0.00071193 |
| GRN | 0.77 | 0.67 | 0.88 | 0.00010553 | 0.00196634 |
| GSTP1 | 0.81 | 0.69 | 0.95 | 0.00813858 | 0.04093517 |
| HMOX2 | 0.75 | 0.64 | 0.87 | 0.00024138 | 0.00358907 |
| IDI2 | 0.64 | 0.53 | 0.77 | 3.6359E-06 | 0.00014604 |
| IGFBP4 | 0.77 | 0.64 | 0.92 | 0.0050547 | 0.0292466 |
| IL1RAP | 0.83 | 0.72 | 0.96 | 0.01013417 | 0.04843343 |
| ING1 | 0.69 | 0.59 | 0.81 | 7.6127E-06 | 0.00024902 |
| IFNL1 | 0.77 | 0.67 | 0.88 | 0.00014849 | 0.00241564 |
| IL34 | 0.72 | 0.61 | 0.84 | 3.449E-05 | 0.00081885 |
| CXCL8 | 0.73 | 0.61 | 0.87 | 0.00048953 | 0.00554299 |
| LRPAP1 | 0.71 | 0.59 | 0.86 | 0.0003489 | 0.00454493 |
| MAD1L1 | 0.74 | 0.63 | 0.86 | 0.00013931 | 0.00233031 |
| MATN3 | 0.63 | 0.54 | 0.73 | 4.3221E-09 | 1.0603E-06 |
| CCL2 | 0.79 | 0.69 | 0.91 | 0.00120357 | 0.0108028 |
| MYOC | 0.79 | 0.68 | 0.93 | 0.00366123 | 0.02449695 |
| NOS1 | 0.66 | 0.55 | 0.8 | 1.035E-05 | 0.00029749 |
| NPM1 | 0.78 | 0.67 | 0.9 | 0.00095894 | 0.00893396 |
| NUDT5 | 0.78 | 0.67 | 0.9 | 0.00080363 | 0.00773164 |
| PAK4 | 1.18 | 1.05 | 1.33 | 0.00573083 | 0.03183316 |
| PAMR1 | 1.27 | 1.09 | 1.49 | 0.00184307 | 0.0150722 |
| PDCD5 | 0.8 | 0.69 | 0.93 | 0.00440375 | 0.02758433 |
| CD274 | 0.74 | 0.63 | 0.86 | 0.00013666 | 0.00231226 |
| PFDN2 | 0.72 | 0.62 | 0.84 | 2.1857E-05 | 0.00057451 |
| PHOSPHO1 | 0.73 | 0.63 | 0.84 | 2.2459E-05 | 0.00057999 |
| PLIN1 | 0.73 | 0.61 | 0.87 | 0.00040984 | 0.00486524 |
| PPCDC | 0.8 | 0.68 | 0.93 | 0.00431569 | 0.02714829 |
| PRDX1 | 0.66 | 0.56 | 0.77 | 1.2328E-07 | 1.2097E-05 |
| SPINK1 | 0.76 | 0.65 | 0.88 | 0.00038933 | 0.00477986 |
| PVR | 0.84 | 0.73 | 0.96 | 0.01029392 | 0.04873455 |
| RBKS | 0.82 | 0.72 | 0.94 | 0.00505537 | 0.0292466 |
| SCARB1 | 0.77 | 0.64 | 0.92 | 0.00344928 | 0.02361551 |
| SCARB2 | 0.71 | 0.6 | 0.85 | 0.00012783 | 0.00218802 |
| SEMA4D | 1.33 | 1.13 | 1.57 | 0.00063221 | 0.00650781 |
| SERPINB6 | 0.79 | 0.67 | 0.93 | 0.00486282 | 0.02898673 |
| SETMAR | 0.71 | 0.61 | 0.83 | 9.6964E-06 | 0.00028546 |
| SOD2 | 0.75 | 0.65 | 0.87 | 0.00011143 | 0.00202492 |
| SSB | 0.78 | 0.67 | 0.9 | 0.00103852 | 0.00961444 |
| STC1 | 0.76 | 0.65 | 0.89 | 0.00044916 | 0.005206 |

| | | | | | |
|---|---|---|---|---|---|
| STC2 | 0.83 | 0.73 | 0.94 | 0.00371126 | 0.02460801 |
| THBS2 | 0.75 | 0.63 | 0.89 | 0.00129944 | 0.011401 |
| TNFRSF10A | 0.79 | 0.67 | 0.93 | 0.00452963 | 0.02789798 |
| TNFRSF6B | 0.75 | 0.62 | 0.89 | 0.00117123 | 0.01057701 |
| TPPP3 | 0.8 | 0.68 | 0.95 | 0.0102965 | 0.04873455 |
| TREML2 | 1.21 | 1.05 | 1.4 | 0.0092279 | 0.04527821 |
| TXNRD1 | 0.75 | 0.65 | 0.87 | 0.00017219 | 0.00267365 |
| VCAN | 0.79 | 0.68 | 0.92 | 0.00278013 | 0.01996268 |
| VSIG4 | 0.84 | 0.75 | 0.95 | 0.00636662 | 0.03432842 |
| WFIKKN1 | 1.29 | 1.1 | 1.51 | 0.00212737 | 0.01683592 |
| XRCC4 | 0.78 | 0.67 | 0.91 | 0.00109345 | 0.00993551 |
| AIF1 | 0.73 | 0.63 | 0.84 | 2.0894E-05 | 0.00055919 |
| AREG | 0.81 | 0.69 | 0.94 | 0.00655832 | 0.03497771 |
| ARSB | 1.36 | 1.17 | 1.56 | 3.0362E-05 | 0.00074487 |
| DEFB4A | 0.82 | 0.71 | 0.94 | 0.004735 | 0.02856524 |
| DEFB4B | 0.82 | 0.71 | 0.94 | 0.004735 | 0.02856524 |
| MUC16 | 0.78 | 0.68 | 0.89 | 0.00026981 | 0.00386893 |
| CALB1 | 0.72 | 0.59 | 0.88 | 0.00154363 | 0.0129841 |
| CAPG | 0.68 | 0.59 | 0.78 | 4.39E-08 | 4.9709E-06 |
| CCL8 | 0.82 | 0.72 | 0.94 | 0.00382566 | 0.02480295 |
| CD300E | 0.73 | 0.63 | 0.85 | 2.5083E-05 | 0.00063659 |
| CDKN1A | 0.81 | 0.7 | 0.94 | 0.00626322 | 0.03402017 |
| CES2 | 0.75 | 0.65 | 0.88 | 0.00028315 | 0.00394641 |
| COX5B | 0.61 | 0.51 | 0.72 | 1.5611E-08 | 2.4953E-06 |
| CPXM1 | 1.24 | 1.07 | 1.44 | 0.00378803 | 0.02478211 |
| CTSF | 1.24 | 1.08 | 1.43 | 0.00252372 | 0.01885746 |
| CTSV | 1.27 | 1.09 | 1.48 | 0.00272518 | 0.01984493 |
| KRT18 | 0.75 | 0.65 | 0.87 | 8.163E-05 | 0.0015405 |
| DCTN2 | 0.63 | 0.53 | 0.74 | 8.3475E-08 | 8.7768E-06 |
| DDAH1 | 0.64 | 0.55 | 0.74 | 5.1511E-09 | 1.0832E-06 |
| DLL1 | 0.76 | 0.65 | 0.89 | 0.0007029 | 0.00713568 |
| DPY30 | 0.68 | 0.59 | 0.79 | 4.9615E-07 | 3.5541E-05 |
| DTX3 | 0.77 | 0.65 | 0.92 | 0.0039409 | 0.02522176 |
| EGFL7 | 1.22 | 1.06 | 1.4 | 0.00532077 | 0.03024004 |
| ELOA | 0.75 | 0.64 | 0.87 | 0.00017255 | 0.00267365 |
| EPHA2 | 0.71 | 0.58 | 0.86 | 0.00040502 | 0.00484704 |
| EPS8L2 | 0.74 | 0.63 | 0.86 | 0.00012138 | 0.0021271 |
| ERP44 | 0.81 | 0.69 | 0.95 | 0.00749599 | 0.03844632 |
| F3 | 0.67 | 0.57 | 0.8 | 8.9758E-06 | 0.00027526 |
| FURIN | 0.81 | 0.7 | 0.94 | 0.0054791 | 0.03102013 |

| | | | | | |
|---|---|---|---|---|---|
| FUS | 0.79 | 0.67 | 0.93 | 0.00523344 | 0.03006226 |
| FXN | 0.74 | 0.62 | 0.87 | 0.00035865 | 0.00463099 |
| GALNT2 | 0.79 | 0.67 | 0.92 | 0.00353187 | 0.02395809 |
| GFER | 0.74 | 0.65 | 0.84 | 6.3584E-06 | 0.00022285 |
| GPC1 | 0.74 | 0.62 | 0.87 | 0.00042917 | 0.0050138 |
| GRPEL1 | 0.79 | 0.69 | 0.9 | 0.00039091 | 0.00477986 |
| GSAP | 1.21 | 1.06 | 1.37 | 0.00392697 | 0.02522176 |
| HBEGF | 1.38 | 1.15 | 1.67 | 0.00055151 | 0.00592576 |
| HSPB6 | 0.7 | 0.6 | 0.82 | 8.5421E-06 | 0.00026753 |
| HTRA2 | 0.78 | 0.68 | 0.9 | 0.0004038 | 0.00484704 |
| HAVCR1 | 0.81 | 0.69 | 0.94 | 0.00775179 | 0.03948318 |
| KLK10 | 0.81 | 0.7 | 0.94 | 0.0050665 | 0.0292466 |
| LRIG1 | 0.83 | 0.72 | 0.95 | 0.0069138 | 0.03608904 |
| LRP1 | 0.71 | 0.61 | 0.82 | 7.565E-06 | 0.00024902 |
| LTA4H | 0.83 | 0.72 | 0.95 | 0.00789139 | 0.0400556 |
| NAMPT | 0.8 | 0.69 | 0.93 | 0.00382461 | 0.02480295 |
| NDUFS6 | 0.57 | 0.47 | 0.69 | 1.6951E-08 | 2.4953E-06 |
| NFKBIE | 0.8 | 0.68 | 0.93 | 0.00384176 | 0.02480295 |
| NUCB2 | 0.76 | 0.65 | 0.89 | 0.00073078 | 0.00726826 |
| OGFR | 0.84 | 0.75 | 0.94 | 0.00321849 | 0.02224236 |
| P4HB | 0.76 | 0.64 | 0.9 | 0.00177458 | 0.01459317 |
| ADCYAP1R1 | 1.19 | 1.07 | 1.33 | 0.00202201 | 0.01635386 |
| PODXL2 | 0.79 | 0.68 | 0.92 | 0.00205078 | 0.01649586 |
| POLR2F | 0.59 | 0.49 | 0.7 | 8.229E-09 | 1.5141E-06 |
| PQBP1 | 0.65 | 0.55 | 0.76 | 1.3488E-07 | 1.2409E-05 |
| RAD23B | 0.79 | 0.68 | 0.91 | 0.00123918 | 0.01105498 |
| RBP2 | 0.69 | 0.59 | 0.81 | 8.1029E-06 | 0.00025929 |
| RRM2B | 0.77 | 0.67 | 0.88 | 0.00011847 | 0.002101 |
| SFTPA1 | 0.68 | 0.59 | 0.8 | 2.2633E-06 | 9.9905E-05 |
| SFTPA2 | 0.73 | 0.63 | 0.85 | 4.2034E-05 | 0.00095191 |
| SLAMF8 | 0.79 | 0.68 | 0.92 | 0.00280536 | 0.02004606 |
| SPARC | 1.3 | 1.1 | 1.55 | 0.00237536 | 0.01802336 |
| SRP14 | 0.82 | 0.7 | 0.95 | 0.00688576 | 0.03607062 |
| ST3GAL1 | 0.78 | 0.68 | 0.89 | 0.00037086 | 0.00472428 |
| TACSTD2 | 0.56 | 0.47 | 0.66 | 2.0138E-11 | 2.9644E-08 |
| TNFRSF12A | 0.71 | 0.58 | 0.88 | 0.00127504 | 0.01130636 |
| TRIAP1 | 0.56 | 0.47 | 0.68 | 1.2182E-09 | 4.8623E-07 |
| VEGFC | 1.33 | 1.11 | 1.59 | 0.00230279 | 0.01774715 |
| FLT1 | 0.72 | 0.58 | 0.9 | 0.00455152 | 0.02791601 |
| VWA1 | 0.79 | 0.69 | 0.9 | 0.00031146 | 0.00421925 |

| OLINK Prognostic Model Output for time to Hospital Discharge<br><br>(adjusted for time from symptom onset) | | | | | |
|---|---|---|---|---|---|
| **Protein** | **HR** | **Lower** | **Upper** | ***P* value** | ***P* adjusted** |
| ADA2 | 1.24 | 1.08 | 1.43 | 0.00235192 | 0.02508716 |
| ANGPTL1 | 0.75 | 0.65 | 0.87 | 7.5382E-05 | 0.00209364 |
| ACTA2 | 0.69 | 0.58 | 0.84 | 0.00014224 | 0.00338655 |
| TNFSF13B | 0.81 | 0.7 | 0.94 | 0.00513165 | 0.03934262 |
| DIABLO | 0.69 | 0.59 | 0.8 | 2.0703E-06 | 0.00015237 |
| CA3 | 0.79 | 0.67 | 0.92 | 0.0025273 | 0.02595099 |
| CCDC80 | 0.78 | 0.66 | 0.93 | 0.00444834 | 0.03617654 |
| CD93 | 0.78 | 0.67 | 0.92 | 0.00255631 | 0.02595099 |
| CLEC5A | 0.72 | 0.6 | 0.86 | 0.00034569 | 0.00660018 |
| CLTA | 0.68 | 0.58 | 0.8 | 3.0813E-06 | 0.0001972 |
| COL4A1 | 0.79 | 0.67 | 0.93 | 0.00547781 | 0.04051926 |
| CSTB | 0.74 | 0.63 | 0.87 | 0.00017376 | 0.00387536 |
| CST3 | 0.79 | 0.67 | 0.92 | 0.00200065 | 0.02231033 |
| DCN | 0.68 | 0.58 | 0.81 | 9.4394E-06 | 0.00043421 |
| DEFA1B | 0.78 | 0.66 | 0.92 | 0.00409881 | 0.03444557 |
| DEFA1 | 0.78 | 0.66 | 0.92 | 0.00409881 | 0.03444557 |
| DKK3 | 0.72 | 0.61 | 0.85 | 0.00010436 | 0.00264851 |
| EPHB4 | 0.78 | 0.66 | 0.92 | 0.0025506 | 0.02595099 |
| FABP2 | 0.76 | 0.64 | 0.9 | 0.00107101 | 0.0146391 |
| FABP6 | 0.79 | 0.67 | 0.94 | 0.0061332 | 0.04373772 |
| FETUB | 1.42 | 1.16 | 1.72 | 0.0005441 | 0.00920595 |
| GAS6 | 0.8 | 0.69 | 0.93 | 0.00299469 | 0.02816243 |
| GDF15 | 0.74 | 0.61 | 0.89 | 0.00151555 | 0.01813732 |
| GGH | 1.31 | 1.11 | 1.55 | 0.00127019 | 0.01584509 |
| IL6ST | 0.81 | 0.69 | 0.94 | 0.00560528 | 0.0410496 |
| GPR37 | 0.81 | 0.69 | 0.94 | 0.00625241 | 0.04382642 |
| IGFBP1 | 0.66 | 0.57 | 0.78 | 5.7706E-07 | 6.0673E-05 |
| IL1RL1 | 0.56 | 0.47 | 0.68 | 5.0746E-10 | 7.4698E-07 |
| IL2RA | 0.77 | 0.65 | 0.9 | 0.00121512 | 0.01582878 |
| LEPR | 0.78 | 0.67 | 0.9 | 0.00094315 | 0.01357665 |
| LGALS1 | 0.74 | 0.63 | 0.87 | 0.00033053 | 0.00640192 |
| LILRA5 | 0.72 | 0.62 | 0.83 | 1.0454E-05 | 0.00046633 |
| LILRB2 | 0.82 | 0.71 | 0.94 | 0.00636494 | 0.04440372 |
| LTBP2 | 0.62 | 0.52 | 0.73 | 2.6798E-08 | 7.3022E-06 |

| | | | | | |
|---|---|---|---|---|---|
| MB | 0.65 | 0.54 | 0.78 | 4.5116E-06 | 0.00025543 |
| NADK | 0.82 | 0.71 | 0.94 | 0.00582856 | 0.04185193 |
| OSMR | 0.75 | 0.65 | 0.87 | 0.00011152 | 0.00278236 |
| PAG1 | 0.74 | 0.64 | 0.87 | 0.00014264 | 0.00338655 |
| PDGFA | 1.28 | 1.07 | 1.53 | 0.00571425 | 0.0415923 |
| PDGFRA | 0.67 | 0.57 | 0.79 | 2.9402E-06 | 0.00019672 |
| PI3 | 0.78 | 0.66 | 0.93 | 0.00536116 | 0.03985674 |
| PLIN3 | 0.74 | 0.63 | 0.86 | 0.00013011 | 0.00319194 |
| PPP1R2 | 0.79 | 0.68 | 0.93 | 0.00367144 | 0.0324248 |
| PRCP | 1.24 | 1.07 | 1.45 | 0.00509972 | 0.03930259 |
| RCOR1 | 0.72 | 0.6 | 0.87 | 0.0004407 | 0.0077227 |
| REG1A | 0.77 | 0.64 | 0.93 | 0.00530846 | 0.03985674 |
| REG1B | 0.76 | 0.64 | 0.91 | 0.003042 | 0.02816243 |
| S100A11 | 0.78 | 0.67 | 0.92 | 0.00227304 | 0.02442271 |
| SPARCL1 | 0.79 | 0.68 | 0.91 | 0.0011672 | 0.0153403 |
| THBD | 0.75 | 0.62 | 0.89 | 0.0010973 | 0.01468388 |
| THBS4 | 0.8 | 0.69 | 0.92 | 0.00253752 | 0.02595099 |
| TINAGL1 | 0.74 | 0.63 | 0.87 | 0.00021316 | 0.00454752 |
| TNC | 0.74 | 0.63 | 0.87 | 0.00018181 | 0.00399431 |
| TNNI3 | 0.74 | 0.62 | 0.88 | 0.00086391 | 0.01297624 |
| VCAM1 | 0.78 | 0.68 | 0.91 | 0.00108346 | 0.0146391 |
| AGRN | 0.77 | 0.66 | 0.91 | 0.00263955 | 0.0261273 |
| AMBN | 1.32 | 1.17 | 1.49 | 6.2909E-06 | 0.00030867 |
| ANGPTL4 | 0.67 | 0.57 | 0.79 | 1.5056E-06 | 0.00011665 |
| TNFSF13 | 0.79 | 0.67 | 0.92 | 0.00340717 | 0.03095897 |
| ATP5IF1 | 0.64 | 0.55 | 0.76 | 1.2307E-07 | 1.6469E-05 |
| CD200R1 | 1.32 | 1.12 | 1.55 | 0.00096798 | 0.01370066 |
| CDSN | 0.78 | 0.66 | 0.92 | 0.00367588 | 0.0324248 |
| CKAP4 | 0.63 | 0.53 | 0.76 | 6.7357E-07 | 6.61E-05 |
| CRELD2 | 0.76 | 0.63 | 0.9 | 0.0017462 | 0.02008129 |
| CRIM1 | 0.76 | 0.66 | 0.88 | 0.00030066 | 0.00608621 |
| CRLF1 | 0.79 | 0.68 | 0.92 | 0.00237213 | 0.02512073 |
| CTSC | 0.8 | 0.69 | 0.94 | 0.0055146 | 0.04058749 |
| CXCL12 | 1.24 | 1.07 | 1.44 | 0.00384248 | 0.03307678 |
| LAMP3 | 0.82 | 0.73 | 0.93 | 0.00123478 | 0.01584509 |
| DFFA | 0.76 | 0.65 | 0.89 | 0.00075929 | 0.01189017 |
| EGF | 1.32 | 1.11 | 1.57 | 0.00176403 | 0.02012912 |
| ENAH | 0.73 | 0.62 | 0.86 | 0.00020781 | 0.00449841 |
| EPCAM | 0.79 | 0.68 | 0.92 | 0.00258395 | 0.02605186 |
| ESM1 | 0.81 | 0.7 | 0.95 | 0.00693803 | 0.04706351 |

| | | | | | |
|---|---|---|---|---|---|
| FLT3LG | 1.28 | 1.08 | 1.51 | 0.00378412 | 0.03295988 |
| FST | 0.72 | 0.61 | 0.85 | 0.00016342 | 0.00370092 |
| FSTL3 | 0.67 | 0.55 | 0.81 | 3.3357E-05 | 0.00111595 |
| GLOD4 | 0.81 | 0.7 | 0.94 | 0.00580966 | 0.04185193 |
| HEXIM1 | 0.69 | 0.59 | 0.81 | 5.7542E-06 | 0.00029208 |
| HSPA1A | 0.78 | 0.67 | 0.91 | 0.00140621 | 0.01707026 |
| IFNGR1 | 0.75 | 0.63 | 0.89 | 0.00093191 | 0.01357665 |
| IL18R1 | 0.78 | 0.67 | 0.91 | 0.00141479 | 0.01707026 |
| IL1RN | 0.79 | 0.68 | 0.92 | 0.00190445 | 0.02156427 |
| IL13 | 1.18 | 1.05 | 1.32 | 0.00529926 | 0.03985674 |
| IL15 | 0.76 | 0.65 | 0.89 | 0.00053826 | 0.00920595 |
| IL6 | 0.76 | 0.64 | 0.9 | 0.0016883 | 0.01972364 |
| ITGB6 | 0.66 | 0.55 | 0.79 | 7.6857E-06 | 0.00036495 |
| JUN | 0.71 | 0.6 | 0.84 | 4.3194E-05 | 0.00135279 |
| KRT19 | 0.75 | 0.66 | 0.86 | 5.4948E-05 | 0.00168506 |
| LGALS9 | 0.75 | 0.65 | 0.88 | 0.00035706 | 0.00660018 |
| LTA | 1.22 | 1.07 | 1.4 | 0.00399758 | 0.03381859 |
| LTBR | 0.76 | 0.64 | 0.91 | 0.00202463 | 0.02240789 |
| MAPK9 | 0.72 | 0.62 | 0.84 | 3.6725E-05 | 0.00120132 |
| MATN2 | 0.76 | 0.64 | 0.89 | 0.00077371 | 0.0119885 |
| TNFRSF11B | 0.75 | 0.61 | 0.91 | 0.00344337 | 0.03109599 |
| PADI2 | 0.77 | 0.64 | 0.92 | 0.00420919 | 0.03480855 |
| PARP1 | 0.74 | 0.61 | 0.9 | 0.00213634 | 0.02312273 |
| PLAUR | 0.72 | 0.6 | 0.86 | 0.00038301 | 0.00687557 |
| PGF | 0.78 | 0.65 | 0.93 | 0.00462454 | 0.03699634 |
| PREB | 0.63 | 0.52 | 0.76 | 2.2447E-06 | 0.00015735 |
| PRELP | 0.8 | 0.69 | 0.93 | 0.00386769 | 0.03310023 |
| PROK1 | 0.69 | 0.6 | 0.8 | 1.4018E-06 | 0.00011464 |
| PSIP1 | 0.77 | 0.67 | 0.9 | 0.00059793 | 0.01000177 |
| PTX3 | 0.81 | 0.71 | 0.93 | 0.00261755 | 0.0261273 |
| AGER | 0.77 | 0.67 | 0.9 | 0.00073533 | 0.01176535 |
| SERPINB8 | 0.78 | 0.68 | 0.9 | 0.00080635 | 0.01236402 |
| ACVRL1 | 0.77 | 0.64 | 0.93 | 0.0057359 | 0.0415923 |
| AGR2 | 0.71 | 0.61 | 0.82 | 5.7055E-06 | 0.00029208 |
| ANXA5 | 0.73 | 0.61 | 0.87 | 0.00037409 | 0.00679829 |
| ARID4B | 0.76 | 0.64 | 0.9 | 0.00126689 | 0.01584509 |
| BAG3 | 0.74 | 0.64 | 0.85 | 1.8491E-05 | 0.00073563 |
| CDHR1 | 1.35 | 1.19 | 1.54 | 5.2806E-06 | 0.00028789 |
| CALCA | 0.67 | 0.56 | 0.79 | 4.0498E-06 | 0.00023845 |
| CRIP2 | 0.77 | 0.67 | 0.89 | 0.00030183 | 0.00608621 |

| | | | | | |
|---|---|---|---|---|---|
| CX3CL1 | 0.71 | 0.61 | 0.83 | 1.2912E-05 | 0.00052796 |
| EFNA1 | 0.71 | 0.6 | 0.86 | 0.00025753 | 0.00533912 |
| ENO1 | 0.83 | 0.72 | 0.95 | 0.00730405 | 0.04864962 |
| EZR | 0.66 | 0.56 | 0.76 | 5.2459E-08 | 9.6525E-06 |
| FABP5 | 0.8 | 0.69 | 0.93 | 0.00324194 | 0.02964063 |
| FGR | 0.81 | 0.69 | 0.94 | 0.00728909 | 0.04864962 |
| FOSB | 0.72 | 0.61 | 0.85 | 6.8523E-05 | 0.00197777 |
| GPKOW | 0.73 | 0.62 | 0.86 | 0.0001584 | 0.00370092 |
| GRN | 0.82 | 0.71 | 0.94 | 0.00482479 | 0.0377771 |
| HMOX2 | 0.73 | 0.63 | 0.86 | 9.8883E-05 | 0.00255362 |
| IDI2 | 0.68 | 0.56 | 0.82 | 8.5856E-05 | 0.00229783 |
| IGFBP4 | 0.78 | 0.66 | 0.93 | 0.00652373 | 0.04466478 |
| IL1RAP | 0.81 | 0.71 | 0.93 | 0.0036792 | 0.0324248 |
| ING1 | 0.71 | 0.61 | 0.84 | 3.1129E-05 | 0.00109101 |
| IFNL1 | 0.8 | 0.69 | 0.93 | 0.00323627 | 0.02964063 |
| IL34 | 0.76 | 0.65 | 0.89 | 0.00085977 | 0.01297624 |
| CXCL8 | 0.74 | 0.62 | 0.88 | 0.00088708 | 0.01318972 |
| LRPAP1 | 0.7 | 0.58 | 0.85 | 0.00035738 | 0.00660018 |
| MAD1L1 | 0.76 | 0.65 | 0.89 | 0.00072066 | 0.01174018 |
| MATN3 | 0.65 | 0.56 | 0.76 | 9.2226E-08 | 1.3576E-05 |
| MMP3 | 0.81 | 0.7 | 0.94 | 0.00685603 | 0.04672256 |
| NEFL | 0.74 | 0.61 | 0.9 | 0.00299339 | 0.02816243 |
| NMNAT1 | 0.81 | 0.69 | 0.94 | 0.00698206 | 0.0471449 |
| NOS1 | 0.68 | 0.57 | 0.82 | 4.2082E-05 | 0.00134663 |
| NPM1 | 0.8 | 0.69 | 0.93 | 0.00411849 | 0.03444557 |
| NUDT5 | 0.81 | 0.7 | 0.93 | 0.00382997 | 0.03307678 |
| PAK4 | 1.2 | 1.07 | 1.35 | 0.00208992 | 0.02278783 |
| PAMR1 | 1.25 | 1.07 | 1.46 | 0.00534308 | 0.03985674 |
| PDCD5 | 0.81 | 0.7 | 0.94 | 0.0049931 | 0.03868338 |
| CD274 | 0.77 | 0.66 | 0.91 | 0.00170513 | 0.01976336 |
| PFDN2 | 0.73 | 0.63 | 0.86 | 8.1854E-05 | 0.00223128 |
| PHOSPHO1 | 0.74 | 0.63 | 0.86 | 7.1304E-05 | 0.00201846 |
| PLIN1 | 0.75 | 0.63 | 0.89 | 0.00108401 | 0.0146391 |
| PRDX1 | 0.69 | 0.59 | 0.81 | 3.6121E-06 | 0.00022154 |
| SPINK1 | 0.75 | 0.64 | 0.88 | 0.00031974 | 0.00636026 |
| SCARB2 | 0.75 | 0.63 | 0.89 | 0.00095 | 0.01357665 |
| SEMA4D | 1.28 | 1.09 | 1.51 | 0.00303334 | 0.02816243 |
| SETMAR | 0.72 | 0.62 | 0.84 | 2.2453E-05 | 0.00082626 |
| SOD2 | 0.76 | 0.66 | 0.89 | 0.00048063 | 0.00832346 |
| SSB | 0.8 | 0.69 | 0.94 | 0.00470065 | 0.03720083 |

| | | | | | |
|---|---|---|---|---|---|
| STC1 | 0.8 | 0.68 | 0.93 | 0.00459313 | 0.03699557 |
| THBS2 | 0.77 | 0.65 | 0.93 | 0.00535914 | 0.03985674 |
| TNFRSF10B | 0.74 | 0.6 | 0.9 | 0.00293667 | 0.02816243 |
| TNFRSF6B | 0.78 | 0.65 | 0.93 | 0.00467863 | 0.03720083 |
| TXNRD1 | 0.79 | 0.68 | 0.92 | 0.00264468 | 0.0261273 |
| VSIG4 | 0.82 | 0.73 | 0.93 | 0.00248589 | 0.02595099 |
| WFIKKN1 | 1.27 | 1.08 | 1.5 | 0.00459931 | 0.03699557 |
| XRCC4 | 0.81 | 0.7 | 0.94 | 0.00618005 | 0.04373772 |
| AIF1 | 0.74 | 0.64 | 0.86 | 9.4415E-05 | 0.00248175 |
| APBB1IP | 0.8 | 0.68 | 0.94 | 0.00648005 | 0.04457303 |
| AREG | 0.78 | 0.67 | 0.92 | 0.00274384 | 0.02692617 |
| ARSB | 1.29 | 1.11 | 1.49 | 0.00069746 | 0.01153549 |
| MUC16 | 0.71 | 0.62 | 0.81 | 1.1733E-06 | 0.00010159 |
| CALB1 | 0.71 | 0.58 | 0.87 | 0.00106148 | 0.0146391 |
| CAPG | 0.68 | 0.59 | 0.78 | 6.7756E-08 | 1.1082E-05 |
| CD300E | 0.78 | 0.68 | 0.91 | 0.00137462 | 0.01699446 |
| CD302 | 0.79 | 0.66 | 0.93 | 0.00623011 | 0.04382642 |
| CES2 | 0.8 | 0.68 | 0.93 | 0.00432943 | 0.03540511 |
| COX5B | 0.61 | 0.51 | 0.72 | 2.6329E-08 | 7.3022E-06 |
| KRT18 | 0.78 | 0.68 | 0.91 | 0.00112937 | 0.01497686 |
| DCTN2 | 0.65 | 0.55 | 0.77 | 7.8903E-07 | 7.2591E-05 |
| DDAH1 | 0.66 | 0.56 | 0.77 | 2.2617E-07 | 2.7743E-05 |
| DLL1 | 0.79 | 0.68 | 0.93 | 0.00370066 | 0.0324248 |
| DPY30 | 0.72 | 0.62 | 0.83 | 1.2032E-05 | 0.00050602 |
| DTX3 | 0.75 | 0.63 | 0.89 | 0.00107099 | 0.0146391 |
| EDA2R | 0.78 | 0.65 | 0.93 | 0.00719852 | 0.04838457 |
| ELOA | 0.78 | 0.67 | 0.91 | 0.00125165 | 0.01584509 |
| EPHA2 | 0.73 | 0.6 | 0.87 | 0.00072579 | 0.01174018 |
| EPS8L2 | 0.73 | 0.62 | 0.85 | 6.6288E-05 | 0.00195153 |
| F3 | 0.68 | 0.57 | 0.81 | 1.0964E-05 | 0.00047467 |
| FUS | 0.79 | 0.67 | 0.93 | 0.00496267 | 0.03865105 |
| FXN | 0.74 | 0.63 | 0.87 | 0.00032727 | 0.00640192 |
| GFER | 0.77 | 0.67 | 0.88 | 0.00016233 | 0.00370092 |
| GPC1 | 0.74 | 0.62 | 0.87 | 0.00039209 | 0.00695362 |
| GRPEL1 | 0.82 | 0.71 | 0.94 | 0.00354946 | 0.03185859 |
| HBEGF | 1.34 | 1.11 | 1.61 | 0.00253751 | 0.02595099 |
| HSPB6 | 0.71 | 0.6 | 0.83 | 2.5605E-05 | 0.00091927 |
| HTRA2 | 0.81 | 0.7 | 0.94 | 0.00473214 | 0.03724977 |
| HAVCR1 | 0.79 | 0.68 | 0.93 | 0.0052799 | 0.03985674 |
| LRP1 | 0.73 | 0.63 | 0.85 | 5.8342E-05 | 0.00175264 |

| | | | | | |
|---|---|---|---|---|---|
| NAMPT | 0.81 | 0.7 | 0.94 | 0.00645257 | 0.04457303 |
| NCS1 | 0.79 | 0.68 | 0.93 | 0.00423973 | 0.03486525 |
| NDUFS6 | 0.57 | 0.47 | 0.7 | 3.4725E-08 | 7.3022E-06 |
| NFKBIE | 0.8 | 0.68 | 0.93 | 0.00420518 | 0.03480855 |
| OGFR | 0.84 | 0.74 | 0.94 | 0.00303061 | 0.02816243 |
| P4HB | 0.75 | 0.63 | 0.9 | 0.0016061 | 0.01891343 |
| ADCYAP1R1 | 1.19 | 1.06 | 1.32 | 0.00276774 | 0.02698088 |
| PODXL2 | 0.79 | 0.68 | 0.92 | 0.00198856 | 0.02231033 |
| POLR2F | 0.59 | 0.49 | 0.72 | 3.1495E-08 | 7.3022E-06 |
| PQBP1 | 0.66 | 0.56 | 0.77 | 4.1017E-07 | 4.6444E-05 |
| RAD23B | 0.79 | 0.68 | 0.91 | 0.00154329 | 0.01832029 |
| RBP2 | 0.71 | 0.6 | 0.83 | 2.1044E-05 | 0.00079426 |
| RET | 1.23 | 1.06 | 1.43 | 0.00642461 | 0.04457303 |
| RRM2B | 0.79 | 0.69 | 0.91 | 0.00093081 | 0.01357665 |
| SFTPA1 | 0.71 | 0.6 | 0.83 | 3.2021E-05 | 0.00109615 |
| SFTPA2 | 0.72 | 0.62 | 0.84 | 2.0211E-05 | 0.00078293 |
| ST3GAL1 | 0.8 | 0.7 | 0.92 | 0.00126116 | 0.01584509 |
| TACSTD2 | 0.58 | 0.49 | 0.69 | 1.6153E-09 | 1.1888E-06 |
| TNFRSF12A | 0.72 | 0.59 | 0.88 | 0.00138542 | 0.01699446 |
| TRIAP1 | 0.59 | 0.49 | 0.71 | 2.5304E-08 | 7.3022E-06 |
| FLT1 | 0.72 | 0.57 | 0.9 | 0.00397444 | 0.03381716 |
| VWA1 | 0.83 | 0.73 | 0.95 | 0.00618033 | 0.04373772 |

| **OLINK Prognostic Model Output for Time to Clinical Failure** | | | | | |
|---|---|---|---|---|---|
| **Protein** | **HR** | **Lower** | **Upper** | ***P* value** | ***P* adjusted** |
| ACAN | 1.26 | 1.06 | 1.5 | 0.00764942 | 0.025475 |
| AHCY | 1.3 | 1.1 | 1.54 | 0.00226398 | 0.0100379 |
| ALCAM | 1.33 | 1.12 | 1.57 | 0.00086241 | 0.00482686 |
| ANGPTL1 | 1.43 | 1.2 | 1.7 | 5.314E-05 | 0.00067433 |
| AOC3 | 1.25 | 1.05 | 1.48 | 0.01218144 | 0.03699802 |
| ACTA2 | 1.32 | 1.1 | 1.58 | 0.00281876 | 0.01186665 |
| TNFSF13B | 1.38 | 1.17 | 1.62 | 0.00010445 | 0.00103885 |
| C1QTNF1 | 1.26 | 1.06 | 1.49 | 0.00729828 | 0.02464008 |
| DIABLO | 1.46 | 1.28 | 1.67 | 2.6916E-08 | 2.6413E-06 |
| CA3 | 1.48 | 1.25 | 1.76 | 7.6264E-06 | 0.00016755 |
| CA4 | 1.25 | 1.05 | 1.49 | 0.01029113 | 0.03263727 |
| CCDC80 | 1.39 | 1.16 | 1.66 | 0.0003726 | 0.00250443 |

| | | | | | |
|---|---|---|---|---|---|
| CD14 | 1.43 | 1.18 | 1.72 | 0.00023035 | 0.00170474 |
| CD209 | 1.28 | 1.08 | 1.52 | 0.00447522 | 0.01667727 |
| CD55 | 1.33 | 1.15 | 1.53 | 0.00014516 | 0.00129149 |
| CD59 | 1.29 | 1.1 | 1.51 | 0.0014777 | 0.00722648 |
| CD93 | 1.35 | 1.15 | 1.6 | 0.00032514 | 0.00226827 |
| ADGRE5 | 1.33 | 1.13 | 1.57 | 0.00063922 | 0.00387217 |
| CDH1 | 1.23 | 1.04 | 1.44 | 0.0131575 | 0.03912695 |
| CDH2 | 1.27 | 1.07 | 1.49 | 0.0048741 | 0.01775908 |
| CLEC1A | 1.24 | 1.07 | 1.43 | 0.00396366 | 0.01522298 |
| CLEC5A | 1.47 | 1.24 | 1.75 | 1.3256E-05 | 0.00024211 |
| CLTA | 1.49 | 1.3 | 1.72 | 1.8224E-08 | 2.0635E-06 |
| CLUL1 | 1.24 | 1.04 | 1.48 | 0.01427656 | 0.04188426 |
| COL1A1 | 1.26 | 1.05 | 1.51 | 0.01172306 | 0.03595072 |
| COL4A1 | 1.27 | 1.08 | 1.5 | 0.00463953 | 0.01720247 |
| COL6A3 | 1.27 | 1.05 | 1.53 | 0.01431817 | 0.04189252 |
| CST6 | 1.31 | 1.11 | 1.53 | 0.00093541 | 0.0050997 |
| CSTB | 1.42 | 1.21 | 1.67 | 2.3445E-05 | 0.00035598 |
| CTSB | 1.36 | 1.16 | 1.6 | 0.00018516 | 0.00146536 |
| CTSD | 1.27 | 1.08 | 1.48 | 0.00293623 | 0.01224401 |
| CTSH | 1.29 | 1.08 | 1.55 | 0.0059106 | 0.02081437 |
| CTSL | 1.43 | 1.2 | 1.71 | 9.5165E-05 | 0.0009865 |
| CTSZ | 1.5 | 1.28 | 1.76 | 4.6956E-07 | 2.16E-05 |
| CXCL16 | 1.4 | 1.16 | 1.69 | 0.00046835 | 0.00295883 |
| CST3 | 1.41 | 1.18 | 1.67 | 0.00011063 | 0.00107135 |
| DCTPP1 | 1.34 | 1.13 | 1.6 | 0.00084672 | 0.00477535 |
| DCN | 1.56 | 1.3 | 1.87 | 2.0835E-06 | 6.5253E-05 |
| DEFA1B | 1.44 | 1.2 | 1.74 | 0.00011706 | 0.00109754 |
| DEFA1 | 1.44 | 1.2 | 1.74 | 0.00011706 | 0.00109754 |
| DKK3 | 1.32 | 1.11 | 1.56 | 0.00150456 | 0.00730927 |
| EIF4EBP1 | 1.37 | 1.16 | 1.62 | 0.00016452 | 0.00137599 |
| EPHB4 | 1.44 | 1.23 | 1.69 | 9.9776E-06 | 0.000198 |
| SELE | 1.25 | 1.06 | 1.48 | 0.00861072 | 0.02816662 |
| FABP6 | 1.33 | 1.15 | 1.53 | 8.2029E-05 | 0.00091475 |
| FAM3C | 1.23 | 1.04 | 1.44 | 0.01490228 | 0.0429279 |
| FCGR2A | 1.35 | 1.12 | 1.63 | 0.00162536 | 0.0077428 |
| CEP43 | 1.23 | 1.04 | 1.46 | 0.0175616 | 0.04796045 |
| FUCA1 | 1.29 | 1.06 | 1.58 | 0.01229527 | 0.03705776 |
| GAS6 | 1.39 | 1.17 | 1.65 | 0.00015077 | 0.00131663 |
| GDF15 | 1.59 | 1.31 | 1.92 | 2.529E-06 | 7.5972E-05 |
| IL6ST | 1.23 | 1.04 | 1.45 | 0.01439417 | 0.04191021 |

| | | | | | |
|---|---|---|---|---|---|
| GPR37 | 1.5 | 1.25 | 1.79 | 1.175E-05 | 0.00022175 |
| HMOX1 | 1.41 | 1.18 | 1.67 | 0.00011911 | 0.00110966 |
| HYOU1 | 1.3 | 1.09 | 1.56 | 0.00437473 | 0.01642755 |
| ICAM1 | 1.27 | 1.06 | 1.51 | 0.00911973 | 0.02950382 |
| ICAM3 | 1.21 | 1.04 | 1.41 | 0.01469083 | 0.04252135 |
| ICAM5 | 1.23 | 1.04 | 1.47 | 0.01717165 | 0.0472001 |
| IGFBP1 | 1.43 | 1.19 | 1.71 | 0.00010643 | 0.00104442 |
| IGFBP2 | 1.39 | 1.14 | 1.68 | 0.0008111 | 0.0046098 |
| IGFBP6 | 1.26 | 1.07 | 1.48 | 0.00454191 | 0.01688307 |
| IGFBP7 | 1.33 | 1.14 | 1.56 | 0.00024328 | 0.00175546 |
| IGFBPL1 | 1.24 | 1.04 | 1.48 | 0.01637748 | 0.04584066 |
| IL1RL1 | 1.77 | 1.45 | 2.17 | 3.1874E-08 | 2.8127E-06 |
| IL2RA | 1.46 | 1.24 | 1.71 | 3.4825E-06 | 9.8581E-05 |
| IL18BP | 1.48 | 1.24 | 1.76 | 1.0673E-05 | 0.00020672 |
| KYAT1 | 1.32 | 1.12 | 1.54 | 0.00066874 | 0.00403437 |
| LACTB2 | 1.26 | 1.07 | 1.48 | 0.0049816 | 0.01810596 |
| LCN2 | 1.35 | 1.11 | 1.64 | 0.00293387 | 0.01224401 |
| LEPR | 1.42 | 1.2 | 1.68 | 4.1874E-05 | 0.00058149 |
| LGALS1 | 1.4 | 1.18 | 1.66 | 0.00010205 | 0.00102893 |
| LILRA5 | 1.41 | 1.19 | 1.68 | 8.9621E-05 | 0.00096294 |
| LRP11 | 1.23 | 1.05 | 1.45 | 0.01197188 | 0.03648573 |
| LTBP2 | 1.58 | 1.31 | 1.9 | 1.4472E-06 | 5.1959E-05 |
| MCFD2 | 1.38 | 1.16 | 1.64 | 0.00021909 | 0.00164386 |
| MB | 1.58 | 1.31 | 1.91 | 1.9959E-06 | 6.5253E-05 |
| MPHOSPH8 | 1.46 | 1.25 | 1.7 | 2.0624E-06 | 6.5253E-05 |
| MSMB | 1.28 | 1.09 | 1.51 | 0.00276468 | 0.01176189 |
| NADK | 1.47 | 1.2 | 1.81 | 0.00023694 | 0.00172663 |
| NECTIN2 | 1.25 | 1.05 | 1.49 | 0.01229487 | 0.03705776 |
| NID1 | 1.36 | 1.14 | 1.61 | 0.00048792 | 0.00306928 |
| NOTCH1 | 1.23 | 1.04 | 1.44 | 0.0150067 | 0.04314426 |
| NOTCH3 | 1.24 | 1.04 | 1.48 | 0.0153853 | 0.0438032 |
| NPDC1 | 1.3 | 1.11 | 1.51 | 0.00101768 | 0.00536943 |
| NRCAM | 1.23 | 1.05 | 1.43 | 0.01090689 | 0.03408691 |
| SPP1 | 1.46 | 1.17 | 1.83 | 0.00102662 | 0.00537787 |
| OSMR | 1.4 | 1.19 | 1.65 | 6.6826E-05 | 0.00079974 |
| PAG1 | 1.56 | 1.29 | 1.88 | 4.8483E-06 | 0.00012215 |
| PCDH17 | 1.33 | 1.12 | 1.57 | 0.00089395 | 0.00492843 |
| PDCD6 | 1.28 | 1.07 | 1.53 | 0.00651672 | 0.0226241 |
| PDGFRA | 1.44 | 1.22 | 1.7 | 2.0146E-05 | 0.00032588 |
| PTGDS | 1.23 | 1.05 | 1.44 | 0.01044216 | 0.03283527 |

| | | | | | |
|---|---|---|---|---|---|
| PI3 | 1.31 | 1.11 | 1.55 | 0.00156306 | 0.0075685 |
| PLA2G2A | 1.33 | 1.1 | 1.62 | 0.00352756 | 0.0139466 |
| PLIN3 | 1.5 | 1.25 | 1.79 | 1.0978E-05 | 0.00020987 |
| PLXNB2 | 1.33 | 1.14 | 1.56 | 0.00037575 | 0.00251409 |
| PPP1R2 | 1.52 | 1.28 | 1.8 | 1.1367E-06 | 4.5164E-05 |
| PRKAR1A | 1.25 | 1.05 | 1.48 | 0.01096784 | 0.03420478 |
| PRTN3 | 1.29 | 1.07 | 1.56 | 0.00911074 | 0.02950382 |
| QDPR | 1.25 | 1.07 | 1.46 | 0.00508957 | 0.01840749 |
| RCOR1 | 1.41 | 1.19 | 1.66 | 4.5129E-05 | 0.00062084 |
| REG1A | 1.28 | 1.08 | 1.52 | 0.00398155 | 0.01522298 |
| REG1B | 1.31 | 1.08 | 1.58 | 0.00486654 | 0.01775908 |
| REN | 1.33 | 1.1 | 1.61 | 0.00316418 | 0.01290214 |
| RETN | 1.42 | 1.18 | 1.71 | 0.00018216 | 0.00144943 |
| RNASET2 | 1.31 | 1.11 | 1.55 | 0.00117299 | 0.00605839 |
| S100A11 | 1.44 | 1.19 | 1.73 | 0.00014059 | 0.00126792 |
| SEMA3F | 1.36 | 1.16 | 1.6 | 0.00018962 | 0.00147681 |
| SIRPA | 1.31 | 1.1 | 1.56 | 0.00277285 | 0.01176264 |
| SLITRK6 | 1.22 | 1.04 | 1.43 | 0.01268198 | 0.03794283 |
| SOD1 | 1.23 | 1.04 | 1.45 | 0.01352127 | 0.03996648 |
| SOST | 1.26 | 1.07 | 1.5 | 0.00692165 | 0.02370811 |
| SPARCL1 | 1.38 | 1.17 | 1.64 | 0.00015998 | 0.00136122 |
| ST6GAL1 | 1.31 | 1.11 | 1.56 | 0.00191198 | 0.00882269 |
| STK11 | 1.26 | 1.08 | 1.48 | 0.00436336 | 0.01642676 |
| TCN2 | 1.33 | 1.12 | 1.59 | 0.00138542 | 0.00693652 |
| TFF3 | 1.24 | 1.06 | 1.45 | 0.00855236 | 0.02816349 |
| TGM2 | 1.24 | 1.06 | 1.46 | 0.00693118 | 0.02370811 |
| THBD | 1.4 | 1.18 | 1.66 | 8.0701E-05 | 0.00090681 |
| THOP1 | 1.3 | 1.1 | 1.54 | 0.00217658 | 0.00979792 |
| TIMP1 | 1.44 | 1.2 | 1.72 | 8.8614E-05 | 0.00096294 |
| TINAGL1 | 1.48 | 1.26 | 1.73 | 1.7023E-06 | 5.8664E-05 |
| TNC | 1.38 | 1.14 | 1.67 | 0.00096528 | 0.00518573 |
| PLAT | 1.35 | 1.13 | 1.6 | 0.00085675 | 0.0048135 |
| TNNI3 | 1.23 | 1.06 | 1.42 | 0.00565604 | 0.02011037 |
| TSPAN1 | 1.23 | 1.05 | 1.44 | 0.00925468 | 0.02980937 |
| TYMP | 1.44 | 1.19 | 1.75 | 0.00020669 | 0.00158141 |
| VAMP5 | 1.22 | 1.04 | 1.43 | 0.01617489 | 0.04552473 |
| VASN | 1.3 | 1.11 | 1.52 | 0.00120215 | 0.00618731 |
| VCAM1 | 1.48 | 1.25 | 1.74 | 2.876E-06 | 8.4669E-05 |
| CHI3L1 | 1.37 | 1.11 | 1.69 | 0.00343047 | 0.0136847 |
| ZBTB17 | 1.41 | 1.19 | 1.68 | 0.00011189 | 0.0010765 |

| | | | | | |
|---|---|---|---|---|---|
| ACTN4 | 1.28 | 1.11 | 1.47 | 0.00049019 | 0.00307045 |
| ADA | 1.26 | 1.08 | 1.48 | 0.00432083 | 0.01630838 |
| ADAM23 | 1.27 | 1.07 | 1.51 | 0.00695781 | 0.02370811 |
| ADGRE2 | 1.39 | 1.18 | 1.65 | 0.00010322 | 0.00103356 |
| AGRN | 1.24 | 1.05 | 1.46 | 0.01041264 | 0.03282763 |
| AGRP | 1.3 | 1.1 | 1.54 | 0.00237 | 0.01041385 |
| AMN | 1.26 | 1.07 | 1.5 | 0.00725585 | 0.02455311 |
| ANGPTL4 | 1.6 | 1.32 | 1.94 | 1.9098E-06 | 6.3893E-05 |
| TNFSF13 | 1.42 | 1.22 | 1.64 | 3.6297E-06 | 9.8943E-05 |
| ATP5IF1 | 1.8 | 1.5 | 2.17 | 4.6994E-10 | 1.3835E-07 |
| B4GALT1 | 1.39 | 1.13 | 1.7 | 0.00163499 | 0.00776355 |
| BACH1 | 1.43 | 1.21 | 1.7 | 2.9184E-05 | 0.00042959 |
| BSG | 1.25 | 1.08 | 1.45 | 0.00246511 | 0.01076747 |
| BTN2A1 | 1.22 | 1.05 | 1.42 | 0.01027715 | 0.03263727 |
| BTN3A2 | 1.37 | 1.17 | 1.6 | 0.00011476 | 0.00108285 |
| C1QA | 1.3 | 1.09 | 1.55 | 0.00306215 | 0.01255566 |
| CCL17 | 0.81 | 0.68 | 0.96 | 0.01691068 | 0.04687857 |
| CCL20 | 1.27 | 1.06 | 1.52 | 0.0085928 | 0.02816662 |
| CCL23 | 1.31 | 1.09 | 1.56 | 0.00384416 | 0.01481312 |
| CCL7 | 1.47 | 1.22 | 1.76 | 4.8536E-05 | 0.00064498 |
| CD22 | 0.8 | 0.68 | 0.95 | 0.01190219 | 0.0363486 |
| CD4 | 1.38 | 1.17 | 1.64 | 0.00016877 | 0.00139958 |
| CD40 | 1.31 | 1.13 | 1.52 | 0.00042186 | 0.00271171 |
| CD48 | 1.26 | 1.08 | 1.47 | 0.0035435 | 0.0139466 |
| CD58 | 1.26 | 1.05 | 1.51 | 0.01101899 | 0.03426347 |
| CDSN | 1.29 | 1.1 | 1.5 | 0.00136759 | 0.00687064 |
| CHRDL1 | 1.39 | 1.17 | 1.65 | 0.00017601 | 0.00144206 |
| CKAP4 | 1.68 | 1.41 | 2 | 5.0718E-09 | 8.2952E-07 |
| CLEC4D | 1.24 | 1.04 | 1.49 | 0.01638056 | 0.04584066 |
| CLEC7A | 1.23 | 1.04 | 1.46 | 0.01528497 | 0.0436883 |
| COLEC12 | 1.3 | 1.1 | 1.54 | 0.0026613 | 0.01135487 |
| CRELD2 | 1.33 | 1.15 | 1.54 | 0.00014017 | 0.00126792 |
| CRIM1 | 1.49 | 1.26 | 1.75 | 1.7137E-06 | 5.8664E-05 |
| CRLF1 | 1.26 | 1.06 | 1.48 | 0.00694583 | 0.02370811 |
| CTSC | 1.31 | 1.1 | 1.55 | 0.0020422 | 0.00933577 |
| CTSO | 1.37 | 1.16 | 1.62 | 0.00016435 | 0.00137599 |
| CXADR | 1.33 | 1.13 | 1.56 | 0.0005089 | 0.00317419 |
| CXCL10 | 1.48 | 1.23 | 1.78 | 4.1386E-05 | 0.00058105 |
| CXCL14 | 1.27 | 1.09 | 1.48 | 0.00193158 | 0.00888527 |
| LAMP3 | 1.31 | 1.09 | 1.58 | 0.00397828 | 0.01522298 |

| | | | | | |
|---|---|---|---|---|---|
| DECR1 | 1.43 | 1.23 | 1.66 | 2.4628E-06 | 7.5527E-05 |
| DFFA | 1.47 | 1.23 | 1.74 | 1.2081E-05 | 0.00022511 |
| DNER | 1.25 | 1.04 | 1.49 | 0.01508044 | 0.04324933 |
| DNPH1 | 1.25 | 1.06 | 1.48 | 0.00755626 | 0.02522775 |
| EGF | 0.82 | 0.69 | 0.96 | 0.01603263 | 0.04529756 |
| EGLN1 | 1.26 | 1.05 | 1.51 | 0.01470338 | 0.04252135 |
| EIF4G1 | 1.35 | 1.14 | 1.61 | 0.00072899 | 0.00427521 |
| ENAH | 1.39 | 1.17 | 1.66 | 0.00018161 | 0.00144943 |
| EPCAM | 1.3 | 1.11 | 1.52 | 0.0012947 | 0.00657172 |
| ESM1 | 1.27 | 1.06 | 1.51 | 0.00794201 | 0.02638969 |
| FABP1 | 1.27 | 1.08 | 1.49 | 0.00469587 | 0.01736762 |
| FABP9 | 1.22 | 1.04 | 1.44 | 0.01707889 | 0.0472001 |
| FCAR | 1.34 | 1.13 | 1.59 | 0.00098289 | 0.00526113 |
| FCRL2 | 1.31 | 1.12 | 1.54 | 0.00068716 | 0.00412857 |
| FIS1 | 1.22 | 1.04 | 1.43 | 0.01711653 | 0.0472001 |
| FST | 1.37 | 1.16 | 1.6 | 0.00014126 | 0.00126792 |
| FSTL3 | 1.61 | 1.33 | 1.94 | 6.0393E-07 | 2.6147E-05 |
| FXYD5 | 1.3 | 1.12 | 1.51 | 0.00075382 | 0.00440329 |
| GBP2 | 1.47 | 1.24 | 1.75 | 8.7219E-06 | 0.00017831 |
| CSF3 | 1.27 | 1.1 | 1.46 | 0.00120676 | 0.00618935 |
| GLOD4 | 1.43 | 1.21 | 1.69 | 2.2248E-05 | 0.0003484 |
| GOPC | 1.26 | 1.08 | 1.47 | 0.00303534 | 0.01248741 |
| HCLS1 | 1.32 | 1.09 | 1.61 | 0.00442263 | 0.01652312 |
| ERBB3 | 1.26 | 1.07 | 1.5 | 0.00657759 | 0.02278168 |
| HEXIM1 | 1.59 | 1.35 | 1.88 | 3.2484E-08 | 2.8127E-06 |
| HPCAL1 | 1.35 | 1.15 | 1.59 | 0.00034522 | 0.00238573 |
| HSPA1A | 1.51 | 1.25 | 1.82 | 1.9036E-05 | 0.00031135 |
| ICA1 | 1.27 | 1.07 | 1.5 | 0.00551894 | 0.01971817 |
| ICAM4 | 1.24 | 1.04 | 1.48 | 0.0142839 | 0.04188426 |
| IFNG | 1.24 | 1.05 | 1.47 | 0.0115014 | 0.03534459 |
| IFNGR1 | 1.42 | 1.22 | 1.66 | 8.0543E-06 | 0.00017208 |
| IL12RB1 | 1.27 | 1.08 | 1.48 | 0.00364859 | 0.01426476 |
| IL15RA | 1.3 | 1.1 | 1.54 | 0.0017495 | 0.00817543 |
| IL16 | 1.38 | 1.13 | 1.68 | 0.00129093 | 0.00657172 |
| IL17C | 1.28 | 1.09 | 1.5 | 0.0021138 | 0.00959342 |
| IL17D | 1.25 | 1.04 | 1.49 | 0.01489962 | 0.0429279 |
| IL18R1 | 1.42 | 1.19 | 1.7 | 9.8706E-05 | 0.00100899 |
| IL1R2 | 1.28 | 1.07 | 1.54 | 0.00751362 | 0.02519373 |
| IL1RN | 1.38 | 1.15 | 1.64 | 0.00042078 | 0.00271171 |
| IL3RA | 1.31 | 1.12 | 1.53 | 0.00088024 | 0.00490799 |

| | | | | | |
|---|---|---|---|---|---|
| IL4R | 1.37 | 1.15 | 1.63 | 0.00041421 | 0.00270292 |
| IL15 | 1.63 | 1.37 | 1.93 | 4.1001E-08 | 3.0177E-06 |
| IL18 | 1.47 | 1.25 | 1.74 | 5.1041E-06 | 0.00012215 |
| IL20 | 1.22 | 1.06 | 1.41 | 0.00596256 | 0.02094721 |
| IL24 | 1.31 | 1.15 | 1.49 | 5.0962E-05 | 0.00065231 |
| IL6 | 1.38 | 1.16 | 1.65 | 0.00036719 | 0.00247941 |
| ITGB6 | 1.41 | 1.19 | 1.68 | 6.8224E-05 | 0.00080679 |
| ITM2A | 1.28 | 1.09 | 1.51 | 0.00324263 | 0.01314916 |
| JUN | 1.56 | 1.32 | 1.85 | 2.0994E-07 | 1.1886E-05 |
| KRT19 | 1.79 | 1.43 | 2.23 | 2.5561E-07 | 1.3438E-05 |
| KYNU | 1.27 | 1.07 | 1.51 | 0.00602712 | 0.02112363 |
| LAIR1 | 1.29 | 1.1 | 1.53 | 0.00222428 | 0.009947 |
| LAMA4 | 1.33 | 1.13 | 1.57 | 0.0007126 | 0.00421264 |
| LAP3 | 1.4 | 1.19 | 1.66 | 7.2912E-05 | 0.00084509 |
| LGALS9 | 1.59 | 1.33 | 1.9 | 2.5116E-07 | 1.3438E-05 |
| LHPP | 1.27 | 1.08 | 1.49 | 0.00335452 | 0.01347751 |
| LIFR | 1.4 | 1.17 | 1.66 | 0.00017634 | 0.00144206 |
| LILRB4 | 1.39 | 1.18 | 1.62 | 6.2489E-05 | 0.00076653 |
| LTBR | 1.39 | 1.18 | 1.63 | 9.1759E-05 | 0.00096478 |
| LY6D | 1.31 | 1.12 | 1.53 | 0.00054902 | 0.00336733 |
| MAPK9 | 1.64 | 1.38 | 1.94 | 1.1446E-08 | 1.404E-06 |
| MATN2 | 1.32 | 1.11 | 1.57 | 0.00159631 | 0.00765399 |
| CSF1 | 1.39 | 1.16 | 1.67 | 0.00029614 | 0.00208573 |
| MEPE | 1.38 | 1.12 | 1.71 | 0.00263729 | 0.01129013 |
| METAP1D | 1.28 | 1.12 | 1.47 | 0.00043219 | 0.00276598 |
| MILR1 | 1.29 | 1.1 | 1.52 | 0.00190495 | 0.00881786 |
| NBN | 1.4 | 1.16 | 1.68 | 0.00036663 | 0.00247941 |
| NFATC3 | 1.23 | 1.06 | 1.43 | 0.00755804 | 0.02522775 |
| NME3 | 1.21 | 1.04 | 1.42 | 0.01642196 | 0.0458693 |
| NPPC | 1.28 | 1.09 | 1.5 | 0.00223509 | 0.009947 |
| NUB1 | 1.42 | 1.21 | 1.67 | 1.8286E-05 | 0.00030245 |
| NUDC | 1.52 | 1.26 | 1.83 | 1.5381E-05 | 0.00027278 |
| OMD | 1.4 | 1.16 | 1.68 | 0.00039923 | 0.00262353 |
| TNFRSF11B | 1.36 | 1.15 | 1.62 | 0.00041499 | 0.00270292 |
| OSCAR | 1.32 | 1.1 | 1.57 | 0.00213308 | 0.00963159 |
| PADI2 | 1.33 | 1.14 | 1.55 | 0.00019443 | 0.00150631 |
| PARP1 | 1.39 | 1.18 | 1.64 | 6.5945E-05 | 0.00079567 |
| PCDH1 | 1.22 | 1.07 | 1.39 | 0.00351076 | 0.01392948 |
| PLAUR | 1.42 | 1.18 | 1.72 | 0.00022 | 0.00164386 |
| PGF | 1.27 | 1.08 | 1.49 | 0.00368628 | 0.01435505 |

| | | | | | |
|---|---|---|---|---|---|
| PRDX3 | 1.4 | 1.2 | 1.64 | 1.6057E-05 | 0.00028138 |
| PREB | 1.6 | 1.38 | 1.86 | 8.81E-10 | 2.1614E-07 |
| PRELP | 1.44 | 1.19 | 1.73 | 0.00015116 | 0.00131663 |
| PRKAB1 | 1.24 | 1.06 | 1.46 | 0.00865537 | 0.02824991 |
| PROK1 | 1.66 | 1.4 | 1.96 | 3.4321E-09 | 7.2172E-07 |
| PSIP1 | 1.69 | 1.39 | 2.05 | 1.0647E-07 | 6.9553E-06 |
| PTX3 | 1.31 | 1.08 | 1.58 | 0.0058504 | 0.02065179 |
| AGER | 1.49 | 1.26 | 1.77 | 4.8998E-06 | 0.00012215 |
| TNFRSF11A | 1.31 | 1.11 | 1.54 | 0.00170513 | 0.00799842 |
| SAMD9L | 1.32 | 1.11 | 1.57 | 0.00134313 | 0.00677086 |
| SCG3 | 1.22 | 1.04 | 1.43 | 0.01544366 | 0.0438032 |
| SCGB1A1 | 1.29 | 1.07 | 1.55 | 0.00709186 | 0.02410904 |
| SCRN1 | 1.36 | 1.16 | 1.59 | 0.00013072 | 0.00120258 |
| SERPINB8 | 1.41 | 1.18 | 1.7 | 0.00021831 | 0.00164386 |
| SH2D1A | 1.26 | 1.08 | 1.47 | 0.00372354 | 0.01442382 |
| SIGLEC1 | 1.35 | 1.15 | 1.59 | 0.00024324 | 0.00175546 |
| SIGLEC10 | 1.29 | 1.08 | 1.54 | 0.00584648 | 0.02065179 |
| SIRPB1 | 1.34 | 1.13 | 1.58 | 0.0007764 | 0.00446429 |
| SLAMF7 | 1.27 | 1.06 | 1.52 | 0.01005767 | 0.03204523 |
| SPINK4 | 1.28 | 1.08 | 1.51 | 0.00419548 | 0.01595799 |
| SRPK2 | 1.47 | 1.23 | 1.76 | 2.2736E-05 | 0.00035229 |
| TPP1 | 1.37 | 1.16 | 1.63 | 0.00023162 | 0.00170474 |
| TRAF2 | 1.25 | 1.05 | 1.48 | 0.01041474 | 0.03282763 |
| TRIM21 | 1.4 | 1.17 | 1.68 | 0.00018633 | 0.00146669 |
| TRIM5 | 1.25 | 1.06 | 1.47 | 0.00719377 | 0.02439914 |
| CKMT1B | 1.28 | 1.08 | 1.52 | 0.00472548 | 0.01743336 |
| CKMT1A | 1.28 | 1.08 | 1.52 | 0.00472548 | 0.01743336 |
| YTHDF3 | 1.33 | 1.14 | 1.56 | 0.00039032 | 0.00257646 |
| ACVRL1 | 1.32 | 1.14 | 1.53 | 0.0002853 | 0.00201906 |
| AGR2 | 1.57 | 1.29 | 1.9 | 4.466E-06 | 0.00011739 |
| AKT1S1 | 1.27 | 1.08 | 1.5 | 0.00353696 | 0.0139466 |
| AMFR | 1.4 | 1.2 | 1.63 | 1.3322E-05 | 0.00024211 |
| ANXA5 | 1.39 | 1.22 | 1.58 | 9.9215E-07 | 4.1727E-05 |
| ARID4B | 1.57 | 1.31 | 1.89 | 1.162E-06 | 4.5164E-05 |
| ASGR1 | 1.37 | 1.14 | 1.65 | 0.00071726 | 0.00422323 |
| ATP6V1F | 1.27 | 1.08 | 1.49 | 0.00342365 | 0.0136847 |
| BAG3 | 1.68 | 1.39 | 2.02 | 6.698E-08 | 4.695E-06 |
| BCAM | 1.44 | 1.21 | 1.73 | 5.0897E-05 | 0.00065231 |
| BST1 | 1.31 | 1.08 | 1.58 | 0.00508792 | 0.01840749 |
| CASP1 | 1.34 | 1.13 | 1.58 | 0.00078829 | 0.00450413 |

| | | | | | |
|---|---|---|---|---|---|
| CASP10 | 1.25 | 1.06 | 1.49 | 0.00993626 | 0.03172705 |
| CCL19 | 1.24 | 1.05 | 1.48 | 0.01243081 | 0.03734318 |
| CD300C | 1.37 | 1.16 | 1.62 | 0.00018817 | 0.00147336 |
| CD300LG | 1.27 | 1.09 | 1.49 | 0.00252925 | 0.01101494 |
| CD74 | 1.37 | 1.15 | 1.62 | 0.00027206 | 0.00193462 |
| CD99 | 1.28 | 1.1 | 1.48 | 0.00139933 | 0.00695884 |
| CD99L2 | 1.24 | 1.07 | 1.44 | 0.00532748 | 0.01912695 |
| CERT1 | 1.25 | 1.04 | 1.5 | 0.01801386 | 0.04901368 |
| CALCA | 1.82 | 1.48 | 2.22 | 6.279E-09 | 9.2426E-07 |
| CLEC11A | 1.29 | 1.09 | 1.52 | 0.00298172 | 0.01236363 |
| CLEC14A | 1.32 | 1.14 | 1.54 | 0.00023063 | 0.00170474 |
| CLPP | 1.28 | 1.1 | 1.5 | 0.00185109 | 0.00859562 |
| CLSPN | 1.2 | 1.06 | 1.35 | 0.00414055 | 0.01578987 |
| CPPED1 | 1.31 | 1.11 | 1.54 | 0.00149271 | 0.00727572 |
| CRIP2 | 1.41 | 1.2 | 1.65 | 2.3502E-05 | 0.00035598 |
| CTSS | 1.37 | 1.16 | 1.61 | 0.00014564 | 0.00129149 |
| CX3CL1 | 1.38 | 1.17 | 1.63 | 0.00015982 | 0.00136122 |
| CXCL13 | 1.33 | 1.13 | 1.56 | 0.00070303 | 0.00417285 |
| DBI | 1.34 | 1.14 | 1.58 | 0.00036054 | 0.00246551 |
| DRAXIN | 1.43 | 1.2 | 1.7 | 5.6113E-05 | 0.00069998 |
| DSG2 | 1.33 | 1.13 | 1.57 | 0.00076846 | 0.00443598 |
| EBAG9 | 1.36 | 1.15 | 1.6 | 0.00021622 | 0.0016406 |
| ECE1 | 1.23 | 1.04 | 1.45 | 0.01537227 | 0.0438032 |
| EFNA1 | 1.39 | 1.19 | 1.63 | 3.8785E-05 | 0.00055428 |
| EFNA4 | 1.25 | 1.07 | 1.47 | 0.00531656 | 0.01912695 |
| ENO1 | 1.31 | 1.1 | 1.56 | 0.00211811 | 0.00959342 |
| EZR | 1.67 | 1.4 | 2 | 2.0262E-08 | 2.1304E-06 |
| F11R | 1.26 | 1.08 | 1.48 | 0.0038166 | 0.0147455 |
| FABP5 | 1.41 | 1.19 | 1.67 | 6.4816E-05 | 0.00078851 |
| FGR | 1.37 | 1.14 | 1.64 | 0.00089278 | 0.00492843 |
| FKBP4 | 1.4 | 1.16 | 1.7 | 0.0006068 | 0.00370629 |
| FKBP7 | 1.34 | 1.15 | 1.56 | 0.00017841 | 0.00144293 |
| FOSB | 1.43 | 1.25 | 1.64 | 3.3966E-07 | 1.7241E-05 |
| GBP4 | 1.27 | 1.08 | 1.51 | 0.00476548 | 0.01752323 |
| GGT5 | 1.23 | 1.05 | 1.44 | 0.01103321 | 0.03426347 |
| GLB1 | 1.24 | 1.05 | 1.47 | 0.01135292 | 0.03503459 |
| GPKOW | 1.58 | 1.32 | 1.89 | 5.271E-07 | 2.3512E-05 |
| GRN | 1.37 | 1.17 | 1.61 | 8.6858E-05 | 0.00095415 |
| GSTP1 | 1.35 | 1.15 | 1.59 | 0.00020394 | 0.0015717 |
| HAVCR2 | 1.29 | 1.08 | 1.53 | 0.00438886 | 0.01643869 |

| | | | | | |
|---|---|---|---|---|---|
| HMOX2 | 1.43 | 1.2 | 1.69 | 5.0749E-05 | 0.00065231 |
| HNMT | 1.3 | 1.12 | 1.52 | 0.00075725 | 0.0044058 |
| IDI2 | 1.42 | 1.21 | 1.67 | 1.7736E-05 | 0.00030008 |
| IGFBP4 | 1.39 | 1.17 | 1.65 | 0.00017812 | 0.00144293 |
| ILKAP | 1.34 | 1.13 | 1.59 | 0.00069414 | 0.0041528 |
| IMPA1 | 1.37 | 1.17 | 1.6 | 8.9348E-05 | 0.00096294 |
| ING1 | 1.58 | 1.33 | 1.87 | 1.1214E-07 | 6.9553E-06 |
| IFNL1 | 1.31 | 1.11 | 1.53 | 0.00094852 | 0.00511434 |
| IL34 | 1.36 | 1.18 | 1.57 | 1.6478E-05 | 0.00028536 |
| KRT14 | 1.24 | 1.06 | 1.45 | 0.00639894 | 0.0222677 |
| LAYN | 1.31 | 1.11 | 1.53 | 0.00094644 | 0.00511434 |
| LBR | 1.28 | 1.07 | 1.54 | 0.00842061 | 0.02785425 |
| LIF | 1.2 | 1.05 | 1.37 | 0.00907651 | 0.02949366 |
| LRPAP1 | 1.25 | 1.08 | 1.45 | 0.00329858 | 0.01330277 |
| MAD1L1 | 1.53 | 1.27 | 1.84 | 9.1349E-06 | 0.0001842 |
| MASP1 | 1.27 | 1.07 | 1.51 | 0.00690051 | 0.02370811 |
| MATN3 | 1.63 | 1.37 | 1.94 | 3.6962E-08 | 2.8636E-06 |
| MAX | 1.32 | 1.12 | 1.55 | 0.0011503 | 0.0059832 |
| CCL2 | 1.44 | 1.19 | 1.73 | 0.00015796 | 0.00135974 |
| MIF | 1.33 | 1.12 | 1.58 | 0.00092194 | 0.00504497 |
| MMP13 | 1.28 | 1.09 | 1.5 | 0.00255552 | 0.0110639 |
| MMP3 | 1.35 | 1.15 | 1.59 | 0.00034128 | 0.00236962 |
| MSR1 | 1.29 | 1.09 | 1.53 | 0.0028401 | 0.01191063 |
| MUC13 | 1.29 | 1.09 | 1.53 | 0.00327852 | 0.01325818 |
| NOS1 | 1.5 | 1.26 | 1.79 | 4.871E-06 | 0.00012215 |
| NPM1 | 1.46 | 1.2 | 1.76 | 0.0001017 | 0.00102893 |
| NUDT5 | 1.51 | 1.29 | 1.78 | 4.1929E-07 | 1.991E-05 |
| PAMR1 | 0.78 | 0.67 | 0.92 | 0.00301578 | 0.01246974 |
| PARK7 | 1.22 | 1.04 | 1.44 | 0.01626491 | 0.04569074 |
| PDCD5 | 1.46 | 1.24 | 1.71 | 5.145E-06 | 0.00012215 |
| CD274 | 1.45 | 1.23 | 1.7 | 6.8255E-06 | 0.00015699 |
| PEBP1 | 1.27 | 1.08 | 1.48 | 0.00323927 | 0.01314916 |
| PFDN2 | 1.51 | 1.27 | 1.8 | 3.1618E-06 | 9.1258E-05 |
| PHOSPHO1 | 1.53 | 1.26 | 1.85 | 1.8184E-05 | 0.00030245 |
| PIK3IP1 | 1.22 | 1.04 | 1.44 | 0.01278792 | 0.03818219 |
| PLAU | 1.39 | 1.17 | 1.65 | 0.00020735 | 0.00158141 |
| PLIN1 | 1.34 | 1.14 | 1.58 | 0.00030647 | 0.00214817 |
| PPCDC | 1.27 | 1.09 | 1.5 | 0.00294996 | 0.01226652 |
| PPP3R1 | 1.42 | 1.19 | 1.7 | 0.00010594 | 0.00104442 |
| PRDX1 | 1.45 | 1.25 | 1.69 | 1.3144E-06 | 4.9612E-05 |

| | | | | | |
|---|---|---|---|---|---|
| PSME1 | 1.31 | 1.11 | 1.55 | 0.00142723 | 0.00700295 |
| PSME2 | 1.34 | 1.14 | 1.58 | 0.00045032 | 0.00285718 |
| SPINK1 | 1.34 | 1.09 | 1.63 | 0.00516546 | 0.01863615 |
| PTK7 | 1.26 | 1.06 | 1.49 | 0.00751164 | 0.02519373 |
| PTPRN2 | 1.21 | 1.04 | 1.4 | 0.01089259 | 0.03408691 |
| PVR | 1.29 | 1.09 | 1.52 | 0.00313277 | 0.01280953 |
| PXN | 1.38 | 1.14 | 1.67 | 0.00088864 | 0.00492843 |
| RAB6B | 0.8 | 0.66 | 0.96 | 0.01464725 | 0.04252135 |
| RBKS | 1.3 | 1.1 | 1.54 | 0.00170619 | 0.00799842 |
| RELT | 1.28 | 1.08 | 1.51 | 0.00481584 | 0.01763412 |
| RGMB | 1.21 | 1.04 | 1.41 | 0.01309335 | 0.039015 |
| RWDD1 | 1.36 | 1.16 | 1.59 | 0.00016924 | 0.00139958 |
| SCARB1 | 1.18 | 1.04 | 1.35 | 0.01117143 | 0.03454694 |
| SCARB2 | 1.37 | 1.18 | 1.6 | 4.5935E-05 | 0.00062608 |
| SCARF2 | 1.23 | 1.04 | 1.46 | 0.0137504 | 0.0405623 |
| SERPINB1 | 1.43 | 1.18 | 1.73 | 0.00023671 | 0.00172663 |
| SERPINB6 | 1.37 | 1.17 | 1.6 | 7.4508E-05 | 0.00084725 |
| SERPINB9 | 1.24 | 1.04 | 1.47 | 0.0154442 | 0.0438032 |
| SETMAR | 1.46 | 1.23 | 1.74 | 2.0818E-05 | 0.00033309 |
| SMPD1 | 1.24 | 1.05 | 1.46 | 0.01259184 | 0.03774987 |
| SNCG | 1.31 | 1.1 | 1.56 | 0.00254023 | 0.01103015 |
| SOD2 | 1.34 | 1.12 | 1.59 | 0.00104979 | 0.00547975 |
| SPOCK1 | 1.32 | 1.12 | 1.57 | 0.00116957 | 0.00605839 |
| SSB | 1.37 | 1.16 | 1.62 | 0.00016098 | 0.00136189 |
| STAMBP | 1.34 | 1.14 | 1.57 | 0.00042022 | 0.00271171 |
| STC1 | 1.48 | 1.25 | 1.75 | 5.0922E-06 | 0.00012215 |
| STC2 | 1.25 | 1.05 | 1.49 | 0.01144364 | 0.03524066 |
| SUSD2 | 1.26 | 1.06 | 1.49 | 0.00980189 | 0.03143439 |
| TARBP2 | 1.25 | 1.06 | 1.47 | 0.00664538 | 0.02296245 |
| TBCC | 1.27 | 1.07 | 1.51 | 0.00673427 | 0.02321508 |
| TDGF1 | 1.23 | 1.05 | 1.44 | 0.01184188 | 0.03623959 |
| TDRKH | 1.28 | 1.1 | 1.49 | 0.00130944 | 0.00662368 |
| THBS2 | 1.43 | 1.17 | 1.74 | 0.00036179 | 0.00246551 |
| THY1 | 1.21 | 1.04 | 1.41 | 0.01564008 | 0.04427345 |
| TIGAR | 1.42 | 1.2 | 1.68 | 4.8636E-05 | 0.00064498 |
| TMSB10 | 1.45 | 1.2 | 1.75 | 9.6991E-05 | 0.00099839 |
| TNFRSF1B | 1.38 | 1.17 | 1.62 | 0.00014005 | 0.00126792 |
| TNFRSF10A | 1.49 | 1.25 | 1.78 | 8.6635E-06 | 0.00017831 |
| TNFRSF10B | 1.39 | 1.18 | 1.64 | 9.0654E-05 | 0.00096478 |
| TNFRSF1A | 1.38 | 1.18 | 1.62 | 6.8725E-05 | 0.00080679 |

| | | | | | |
|---|---|---|---|---|---|
| TNFRSF21 | 1.22 | 1.04 | 1.43 | 0.01223548 | 0.03705776 |
| TNFRSF6B | 1.43 | 1.18 | 1.74 | 0.0002564 | 0.00183211 |
| TPPP3 | 1.24 | 1.09 | 1.41 | 0.00101771 | 0.00536943 |
| MAPT | 1.23 | 1.04 | 1.45 | 0.01784919 | 0.04865558 |
| TXLNA | 1.4 | 1.18 | 1.65 | 0.00011451 | 0.00108285 |
| TXNDC5 | 1.23 | 1.04 | 1.46 | 0.01814205 | 0.0492714 |
| TXNRD1 | 1.39 | 1.18 | 1.64 | 8.4227E-05 | 0.0009322 |
| ULBP2 | 1.36 | 1.13 | 1.63 | 0.00102281 | 0.00537708 |
| VCAN | 1.33 | 1.1 | 1.59 | 0.00262712 | 0.01129013 |
| VSIG4 | 1.58 | 1.28 | 1.96 | 2.37E-05 | 0.00035598 |
| VSTM1 | 1.23 | 1.04 | 1.46 | 0.01718699 | 0.0472001 |
| WARS1 | 1.42 | 1.18 | 1.7 | 0.00015416 | 0.00133481 |
| XRCC4 | 1.42 | 1.19 | 1.68 | 6.9059E-05 | 0.00080679 |
| ABL1 | 1.31 | 1.11 | 1.54 | 0.00139601 | 0.00695884 |
| ADAMTS15 | 1.23 | 1.04 | 1.45 | 0.01617383 | 0.04552473 |
| AIF1 | 1.52 | 1.27 | 1.82 | 4.0271E-06 | 0.00010778 |
| AKR1B1 | 1.27 | 1.09 | 1.48 | 0.00210351 | 0.00958629 |
| AMBP | 1.21 | 1.03 | 1.42 | 0.01725246 | 0.04729167 |
| APBB1IP | 1.35 | 1.11 | 1.63 | 0.00230554 | 0.01016096 |
| APEX1 | 1.26 | 1.04 | 1.52 | 0.01828536 | 0.04956916 |
| AREG | 1.4 | 1.16 | 1.69 | 0.00053187 | 0.00328958 |
| ATOX1 | 1.24 | 1.04 | 1.46 | 0.01440664 | 0.04191021 |
| ATP6AP2 | 1.28 | 1.1 | 1.48 | 0.00141696 | 0.0069758 |
| CALB1 | 1.41 | 1.19 | 1.68 | 9.3153E-05 | 0.00097249 |
| CAPG | 1.85 | 1.49 | 2.31 | 3.4977E-08 | 2.8603E-06 |
| CCL8 | 1.25 | 1.05 | 1.48 | 0.01046178 | 0.03283527 |
| CD300E | 1.47 | 1.22 | 1.76 | 3.4882E-05 | 0.00050838 |
| CD300LF | 1.26 | 1.05 | 1.51 | 0.01219024 | 0.03699802 |
| CD302 | 1.33 | 1.13 | 1.57 | 0.00076051 | 0.00440734 |
| CD38 | 1.29 | 1.09 | 1.54 | 0.00348435 | 0.01386207 |
| CDC37 | 1.31 | 1.12 | 1.54 | 0.00099555 | 0.00530961 |
| CDKN1A | 1.39 | 1.19 | 1.62 | 4.1447E-05 | 0.00058105 |
| CEACAM5 | 1.27 | 1.08 | 1.5 | 0.0043149 | 0.01630838 |
| CEP85 | 1.26 | 1.07 | 1.48 | 0.00612258 | 0.0214072 |
| CFC1 | 1.23 | 1.04 | 1.44 | 0.01337589 | 0.03961632 |
| CNPY4 | 1.34 | 1.16 | 1.55 | 0.00010841 | 0.00105677 |
| COX5B | 1.73 | 1.47 | 2.05 | 1.3069E-10 | 9.6191E-08 |
| CREG1 | 1.26 | 1.06 | 1.49 | 0.00739067 | 0.0248949 |
| KRT18 | 1.54 | 1.28 | 1.85 | 3.6072E-06 | 9.8943E-05 |
| DAB2 | 1.31 | 1.12 | 1.54 | 0.00078945 | 0.00450413 |

| | | | | | |
|---|---|---|---|---|---|
| DCTN2 | 1.46 | 1.26 | 1.71 | 1.1659E-06 | 4.5164E-05 |
| DDAH1 | 1.5 | 1.29 | 1.74 | 1.1882E-07 | 6.9961E-06 |
| DLL1 | 1.4 | 1.19 | 1.66 | 7.452E-05 | 0.00084725 |
| DNAJB1 | 1.31 | 1.11 | 1.55 | 0.0016508 | 0.00781342 |
| DPY30 | 1.66 | 1.38 | 2 | 1.134E-07 | 6.9553E-06 |
| DTX3 | 1.27 | 1.09 | 1.49 | 0.00243399 | 0.01066318 |
| EDA2R | 1.26 | 1.05 | 1.51 | 0.01414462 | 0.04164176 |
| ELOA | 1.51 | 1.26 | 1.8 | 6.0843E-06 | 0.00014216 |
| EPHA2 | 1.38 | 1.18 | 1.62 | 4.9971E-05 | 0.00065231 |
| EPS8L2 | 1.46 | 1.23 | 1.73 | 1.5168E-05 | 0.00027229 |
| ERP44 | 1.23 | 1.05 | 1.43 | 0.00853218 | 0.02816003 |
| F3 | 1.44 | 1.22 | 1.7 | 2.1494E-05 | 0.0003402 |
| FGF23 | 1.23 | 1.04 | 1.46 | 0.01714643 | 0.0472001 |
| FLT3 | 1.23 | 1.04 | 1.45 | 0.01331721 | 0.03952204 |
| FOLR1 | 1.24 | 1.05 | 1.47 | 0.01031001 | 0.03263727 |
| FURIN | 1.32 | 1.12 | 1.56 | 0.00094535 | 0.00511434 |
| FUS | 1.49 | 1.23 | 1.81 | 5.7726E-05 | 0.00071405 |
| FXN | 1.5 | 1.28 | 1.75 | 3.8069E-07 | 1.8679E-05 |
| GALNT2 | 1.34 | 1.13 | 1.58 | 0.00062994 | 0.00383168 |
| GFER | 1.45 | 1.23 | 1.71 | 6.9473E-06 | 0.00015733 |
| GPC1 | 1.37 | 1.15 | 1.63 | 0.00051569 | 0.00320296 |
| GRPEL1 | 1.43 | 1.19 | 1.7 | 9.1381E-05 | 0.00096478 |
| HAGH | 1.24 | 1.07 | 1.44 | 0.00422534 | 0.01603016 |
| HGS | 1.25 | 1.06 | 1.46 | 0.00622508 | 0.02171402 |
| HPGDS | 1.29 | 1.11 | 1.51 | 0.00100388 | 0.00533471 |
| HS3ST3B1 | 1.3 | 1.08 | 1.57 | 0.00537325 | 0.01924434 |
| HS6ST1 | 1.3 | 1.09 | 1.54 | 0.00280462 | 0.01186322 |
| HSPB6 | 1.47 | 1.22 | 1.79 | 7.4825E-05 | 0.00084725 |
| HTRA2 | 1.43 | 1.2 | 1.7 | 4.8611E-05 | 0.00064498 |
| IDUA | 1.3 | 1.08 | 1.57 | 0.0056396 | 0.02010046 |
| IGF1R | 1.27 | 1.08 | 1.48 | 0.00303702 | 0.01248741 |
| INPP1 | 1.25 | 1.08 | 1.46 | 0.00371105 | 0.01441337 |
| INPPL1 | 1.25 | 1.08 | 1.45 | 0.00365341 | 0.01426476 |
| ITGB5 | 1.25 | 1.07 | 1.47 | 0.00477365 | 0.01752323 |
| KLK10 | 1.28 | 1.08 | 1.5 | 0.00336022 | 0.01347751 |
| KLK11 | 1.24 | 1.05 | 1.46 | 0.00978112 | 0.03143439 |
| LAG3 | 1.22 | 1.04 | 1.43 | 0.01231063 | 0.03705776 |
| LAT2 | 1.29 | 1.1 | 1.52 | 0.00183436 | 0.00854484 |
| LPCAT2 | 1.35 | 1.13 | 1.6 | 0.00091409 | 0.00502068 |
| LRP1 | 1.29 | 1.09 | 1.52 | 0.00282155 | 0.01186665 |

| | | | | | |
|---|---|---|---|---|---|
| LTA4H | 1.24 | 1.04 | 1.47 | 0.01648833 | 0.04596747 |
| LTBP3 | 1.28 | 1.07 | 1.53 | 0.00807031 | 0.02675562 |
| LYAR | 1.3 | 1.12 | 1.5 | 0.00069683 | 0.0041528 |
| LYN | 1.48 | 1.23 | 1.78 | 2.5956E-05 | 0.00038593 |
| MANSC1 | 1.45 | 1.23 | 1.71 | 8.073E-06 | 0.00017208 |
| NAMPT | 1.27 | 1.06 | 1.51 | 0.00893157 | 0.02908687 |
| NBL1 | 1.33 | 1.11 | 1.59 | 0.00222324 | 0.009947 |
| NDUFS6 | 1.71 | 1.46 | 2 | 2.5941E-11 | 3.8184E-08 |
| NECTIN4 | 1.27 | 1.09 | 1.48 | 0.00223672 | 0.009947 |
| NFKBIE | 1.41 | 1.18 | 1.68 | 0.00011399 | 0.00108285 |
| NUCB2 | 1.42 | 1.21 | 1.67 | 1.7215E-05 | 0.00029466 |
| OGFR | 1.59 | 1.3 | 1.94 | 7.5432E-06 | 0.00016755 |
| P4HB | 1.29 | 1.09 | 1.54 | 0.00357164 | 0.01401987 |
| ADCYAP1R1 | 0.76 | 0.6 | 0.95 | 0.01733279 | 0.04742354 |
| PODXL2 | 1.41 | 1.2 | 1.66 | 3.5446E-05 | 0.00051153 |
| POLR2F | 1.81 | 1.51 | 2.18 | 2.2285E-10 | 1.0935E-07 |
| PQBP1 | 1.68 | 1.41 | 2.01 | 1.0785E-08 | 1.404E-06 |
| PRDX6 | 1.3 | 1.1 | 1.53 | 0.0015946 | 0.00765399 |
| PSMD9 | 1.27 | 1.09 | 1.49 | 0.00257091 | 0.01109787 |
| RABEPK | 1.29 | 1.13 | 1.49 | 0.00025148 | 0.00180577 |
| RAD23B | 1.46 | 1.24 | 1.73 | 8.183E-06 | 0.00017208 |
| RANGAP1 | 0.79 | 0.66 | 0.96 | 0.01673691 | 0.04657228 |
| RARRES1 | 1.25 | 1.05 | 1.47 | 0.0098835 | 0.0316272 |
| RBP2 | 1.3 | 1.1 | 1.55 | 0.00230457 | 0.01016096 |
| RRM2B | 1.38 | 1.17 | 1.62 | 0.00012751 | 0.00118049 |
| S100A4 | 1.26 | 1.07 | 1.49 | 0.00578403 | 0.02051587 |
| SEMA4C | 1.36 | 1.15 | 1.62 | 0.00044725 | 0.00284999 |
| SEPTIN9 | 1.3 | 1.11 | 1.54 | 0.00160901 | 0.00768982 |
| SF3B4 | 1.3 | 1.1 | 1.53 | 0.0019893 | 0.00912227 |
| SFTPA1 | 1.47 | 1.26 | 1.72 | 1.3582E-06 | 4.9981E-05 |
| SFTPA2 | 1.32 | 1.1 | 1.58 | 0.00263845 | 0.01129013 |
| SLAMF8 | 1.48 | 1.24 | 1.75 | 1.0088E-05 | 0.000198 |
| SPINK6 | 1.45 | 1.2 | 1.76 | 0.00014956 | 0.00131663 |
| SRP14 | 1.37 | 1.13 | 1.66 | 0.00140592 | 0.00696808 |
| ST3GAL1 | 1.24 | 1.06 | 1.47 | 0.00917389 | 0.02961395 |
| SUGT1 | 1.23 | 1.04 | 1.45 | 0.01434364 | 0.04189252 |
| TACSTD2 | 1.7 | 1.42 | 2.03 | 4.5864E-09 | 8.2952E-07 |
| TBL1X | 1.25 | 1.06 | 1.49 | 0.00858424 | 0.02816662 |
| TGFBR2 | 1.28 | 1.1 | 1.49 | 0.00141273 | 0.0069758 |
| TNFRSF12A | 1.41 | 1.17 | 1.71 | 0.00035866 | 0.00246551 |

| | | | | | |
|---|---|---|---|---|---|
| TNFRSF19 | 1.23 | 1.05 | 1.45 | 0.01109605 | 0.03438607 |
| TRIAP1 | 1.64 | 1.4 | 1.91 | 3.5752E-10 | 1.3157E-07 |
| UBAC1 | 1.22 | 1.04 | 1.43 | 0.0167992 | 0.04665741 |
| USO1 | 1.32 | 1.11 | 1.58 | 0.00157174 | 0.0075856 |
| VWA1 | 1.41 | 1.15 | 1.73 | 0.00083747 | 0.00474135 |
| WFDC12 | 1.3 | 1.11 | 1.53 | 0.00126705 | 0.00647604 |
| CCN4 | 1.42 | 1.18 | 1.7 | 0.00018028 | 0.00144943 |
| ZBTB16 | 1.22 | 1.04 | 1.44 | 0.01510201 | 0.04324933 |

| **OLINK Prognostic Model Output for Time to Clinical Failure (adjusted for time from symptom onset)** | | | | | |
|---|---|---|---|---|---|
| **Protein** | **HR** | **Lower** | **Upper** | ***P* value** | ***P* adjusted** |
| ACAN | 1.25 | 1.05 | 1.49 | 0.01146573 | 0.03707811 |
| AHCY | 1.28 | 1.08 | 1.52 | 0.00452191 | 0.01885624 |
| ALCAM | 1.32 | 1.11 | 1.56 | 0.00150067 | 0.00856199 |
| ANGPTL1 | 1.38 | 1.15 | 1.64 | 0.00046327 | 0.0035891 |
| ACTA2 | 1.32 | 1.1 | 1.58 | 0.00277751 | 0.01281659 |
| TNFSF13B | 1.33 | 1.12 | 1.57 | 0.00083293 | 0.00552284 |
| DIABLO | 1.45 | 1.26 | 1.66 | 9.8774E-08 | 1.2116E-05 |
| CA3 | 1.43 | 1.2 | 1.71 | 6.8027E-05 | 0.00105406 |
| CA4 | 1.25 | 1.05 | 1.49 | 0.01141067 | 0.0369967 |
| CCDC80 | 1.37 | 1.14 | 1.64 | 0.00068649 | 0.00472206 |
| CD14 | 1.39 | 1.14 | 1.68 | 0.00089746 | 0.00584542 |
| CD209 | 1.26 | 1.06 | 1.5 | 0.00853538 | 0.02973887 |
| CD55 | 1.32 | 1.14 | 1.54 | 0.00027609 | 0.002613 |
| CD59 | 1.29 | 1.1 | 1.52 | 0.00171267 | 0.00926857 |
| CD93 | 1.38 | 1.16 | 1.63 | 0.00019633 | 0.0021182 |
| ADGRE5 | 1.3 | 1.1 | 1.54 | 0.0022031 | 0.01077454 |
| CDH1 | 1.22 | 1.04 | 1.44 | 0.01578084 | 0.04692807 |
| CDH2 | 1.26 | 1.07 | 1.49 | 0.00626551 | 0.02341654 |
| CLEC1A | 1.22 | 1.06 | 1.42 | 0.00737599 | 0.02641715 |
| CLEC5A | 1.47 | 1.23 | 1.76 | 1.9068E-05 | 0.00046012 |
| CLTA | 1.46 | 1.26 | 1.68 | 2.2925E-07 | 1.7761E-05 |
| CLUL1 | 1.25 | 1.05 | 1.49 | 0.01325769 | 0.04082704 |

| | | | | | |
|---|---|---|---|---|---|
| COL1A1 | 1.27 | 1.05 | 1.53 | 0.01315965 | 0.04069539 |
| COL4A1 | 1.24 | 1.04 | 1.46 | 0.01495038 | 0.04491216 |
| COL6A3 | 1.31 | 1.08 | 1.6 | 0.00672651 | 0.02475356 |
| CST6 | 1.32 | 1.13 | 1.56 | 0.00065085 | 0.00451914 |
| CSTB | 1.41 | 1.19 | 1.67 | 5.1178E-05 | 0.00080142 |
| CTSB | 1.35 | 1.14 | 1.58 | 0.0003834 | 0.00323163 |
| CTSD | 1.24 | 1.06 | 1.46 | 0.00620053 | 0.02340303 |
| CTSH | 1.27 | 1.06 | 1.52 | 0.01070238 | 0.03524364 |
| CTSL | 1.4 | 1.16 | 1.68 | 0.00035554 | 0.0031152 |
| CTSZ | 1.46 | 1.24 | 1.72 | 4.0302E-06 | 0.00016479 |
| CXCL16 | 1.4 | 1.16 | 1.7 | 0.00055544 | 0.00406767 |
| CST3 | 1.42 | 1.19 | 1.69 | 0.00012504 | 0.00154669 |
| DCTPP1 | 1.35 | 1.13 | 1.62 | 0.00081586 | 0.00545595 |
| DCN | 1.52 | 1.26 | 1.83 | 1.1086E-05 | 0.00032637 |
| DEFA1B | 1.45 | 1.2 | 1.75 | 0.00011536 | 0.0014514 |
| DEFA1 | 1.45 | 1.2 | 1.75 | 0.00011536 | 0.0014514 |
| DKK3 | 1.28 | 1.07 | 1.52 | 0.00556091 | 0.02191384 |
| EIF4EBP1 | 1.39 | 1.17 | 1.64 | 0.00012118 | 0.00151173 |
| EPHB4 | 1.42 | 1.2 | 1.67 | 3.29E-05 | 0.00063851 |
| SELE | 1.25 | 1.06 | 1.48 | 0.00890699 | 0.03042015 |
| FABP6 | 1.31 | 1.14 | 1.52 | 0.00017264 | 0.00192524 |
| FCGR2A | 1.42 | 1.17 | 1.72 | 0.00038419 | 0.00323163 |
| CEP43 | 1.27 | 1.06 | 1.51 | 0.00821409 | 0.02892616 |
| FUCA1 | 1.31 | 1.07 | 1.61 | 0.00883114 | 0.03042015 |
| GAS6 | 1.33 | 1.11 | 1.58 | 0.0015529 | 0.00865087 |
| GDF15 | 1.55 | 1.28 | 1.89 | 8.2363E-06 | 0.00027554 |
| GPR37 | 1.47 | 1.23 | 1.77 | 3.3309E-05 | 0.00063851 |
| HMOX1 | 1.4 | 1.18 | 1.67 | 0.00015372 | 0.00179784 |
| HSPG2 | 1.23 | 1.04 | 1.47 | 0.01638318 | 0.0484258 |
| HYOU1 | 1.35 | 1.12 | 1.63 | 0.00160651 | 0.00879103 |
| ICAM1 | 1.31 | 1.09 | 1.57 | 0.00391086 | 0.01683271 |
| ICAM5 | 1.28 | 1.07 | 1.52 | 0.00604887 | 0.02306721 |
| IGFBP1 | 1.4 | 1.17 | 1.68 | 0.00026031 | 0.00249383 |
| IGFBP2 | 1.38 | 1.13 | 1.67 | 0.00121512 | 0.00739112 |
| IGFBP6 | 1.26 | 1.07 | 1.48 | 0.00613993 | 0.02323389 |
| IGFBP7 | 1.33 | 1.14 | 1.56 | 0.00037507 | 0.00320989 |
| IGFBPL1 | 1.26 | 1.05 | 1.5 | 0.01167949 | 0.03761973 |
| IL1RL1 | 1.72 | 1.4 | 2.11 | 1.744E-07 | 1.5101E-05 |
| IL2RA | 1.46 | 1.24 | 1.72 | 4.2236E-06 | 0.00016803 |
| IL18BP | 1.42 | 1.19 | 1.69 | 0.00011452 | 0.0014514 |

| | | | | | |
|---|---|---|---|---|---|
| KYAT1 | 1.29 | 1.1 | 1.51 | 0.00191367 | 0.00995381 |
| LACTB2 | 1.25 | 1.06 | 1.47 | 0.00885028 | 0.03042015 |
| LCN2 | 1.36 | 1.11 | 1.66 | 0.00280064 | 0.01288295 |
| LEPR | 1.39 | 1.17 | 1.64 | 0.00015389 | 0.00179784 |
| LGALS1 | 1.38 | 1.17 | 1.64 | 0.00021708 | 0.00220441 |
| LILRA5 | 1.42 | 1.19 | 1.69 | 0.00010609 | 0.00141221 |
| LTBP2 | 1.52 | 1.26 | 1.83 | 1.0721E-05 | 0.00032207 |
| MCFD2 | 1.35 | 1.14 | 1.62 | 0.00076004 | 0.00515563 |
| MB | 1.52 | 1.26 | 1.85 | 1.722E-05 | 0.00043703 |
| MPHOSPH8 | 1.45 | 1.24 | 1.71 | 6.3564E-06 | 0.00022821 |
| MSMB | 1.27 | 1.08 | 1.51 | 0.0043835 | 0.01838325 |
| NADK | 1.44 | 1.17 | 1.78 | 0.00063169 | 0.00444904 |
| NECTIN2 | 1.25 | 1.05 | 1.5 | 0.01323518 | 0.04082704 |
| NID1 | 1.35 | 1.14 | 1.6 | 0.00061203 | 0.00437334 |
| NPDC1 | 1.31 | 1.12 | 1.53 | 0.00092366 | 0.0059633 |
| NRCAM | 1.23 | 1.05 | 1.43 | 0.01148615 | 0.03707811 |
| SPP1 | 1.43 | 1.14 | 1.79 | 0.00219267 | 0.01077454 |
| OSMR | 1.39 | 1.18 | 1.64 | 0.00010841 | 0.00141221 |
| PAG1 | 1.52 | 1.26 | 1.85 | 1.8509E-05 | 0.00045409 |
| PCDH17 | 1.3 | 1.1 | 1.54 | 0.00250015 | 0.01172043 |
| PDCD6 | 1.31 | 1.09 | 1.58 | 0.00335145 | 0.01513295 |
| PDGFRA | 1.42 | 1.2 | 1.68 | 4.0957E-05 | 0.00072636 |
| PTGDS | 1.25 | 1.07 | 1.47 | 0.00563445 | 0.02199976 |
| PI3 | 1.31 | 1.1 | 1.55 | 0.00213928 | 0.01060275 |
| PLA2G2A | 1.29 | 1.06 | 1.57 | 0.01131122 | 0.03675521 |
| PLIN3 | 1.5 | 1.25 | 1.81 | 1.9915E-05 | 0.00046532 |
| PLXNB2 | 1.3 | 1.11 | 1.53 | 0.00131822 | 0.00792011 |
| PPP1R2 | 1.51 | 1.27 | 1.79 | 2.4746E-06 | 0.00011038 |
| PRKAR1A | 1.25 | 1.05 | 1.48 | 0.01366456 | 0.04164436 |
| PRTN3 | 1.29 | 1.06 | 1.56 | 0.00938374 | 0.03158592 |
| QDPR | 1.24 | 1.05 | 1.45 | 0.00898589 | 0.03047749 |
| RCOR1 | 1.44 | 1.21 | 1.71 | 4.4727E-05 | 0.00074937 |
| REG1A | 1.27 | 1.07 | 1.51 | 0.00556778 | 0.02191384 |
| REG1B | 1.29 | 1.07 | 1.55 | 0.00872018 | 0.03020259 |
| REN | 1.3 | 1.07 | 1.58 | 0.00737184 | 0.02641715 |
| RETN | 1.39 | 1.15 | 1.68 | 0.00050869 | 0.00382035 |
| RNASET2 | 1.28 | 1.08 | 1.51 | 0.0036789 | 0.01626229 |
| S100A11 | 1.46 | 1.21 | 1.77 | 0.00010709 | 0.00141221 |
| SEMA3F | 1.34 | 1.13 | 1.58 | 0.00056267 | 0.00410025 |
| SIRPA | 1.28 | 1.08 | 1.53 | 0.00538591 | 0.02166136 |

| | | | | | |
|---|---|---|---|---|---|
| SOD1 | 1.23 | 1.04 | 1.45 | 0.01437971 | 0.04355337 |
| SOST | 1.28 | 1.08 | 1.52 | 0.00483857 | 0.01989491 |
| SPARCL1 | 1.41 | 1.19 | 1.67 | 9.1118E-05 | 0.0012852 |
| ST6GAL1 | 1.28 | 1.08 | 1.52 | 0.0054165 | 0.02172502 |
| STK11 | 1.27 | 1.08 | 1.49 | 0.00393442 | 0.01688474 |
| TCN2 | 1.29 | 1.08 | 1.54 | 0.00602375 | 0.02306721 |
| TFF3 | 1.24 | 1.05 | 1.46 | 0.00960539 | 0.03206154 |
| TGM2 | 1.27 | 1.08 | 1.49 | 0.00370988 | 0.01626286 |
| THBD | 1.42 | 1.19 | 1.68 | 7.1283E-05 | 0.001093 |
| THOP1 | 1.34 | 1.12 | 1.59 | 0.0011355 | 0.0070229 |
| TIMP1 | 1.41 | 1.18 | 1.69 | 0.00019895 | 0.00212215 |
| TINAGL1 | 1.46 | 1.24 | 1.72 | 7.382E-06 | 0.00025872 |
| TNC | 1.44 | 1.19 | 1.75 | 0.00021552 | 0.00220441 |
| PLAT | 1.33 | 1.11 | 1.59 | 0.00159542 | 0.00876288 |
| TSPAN1 | 1.22 | 1.04 | 1.43 | 0.01275351 | 0.0399429 |
| TYMP | 1.42 | 1.16 | 1.74 | 0.00060747 | 0.00436193 |
| VASN | 1.33 | 1.13 | 1.56 | 0.00064176 | 0.00449526 |
| VCAM1 | 1.45 | 1.23 | 1.72 | 1.0292E-05 | 0.00032082 |
| CHI3L1 | 1.35 | 1.09 | 1.67 | 0.00544817 | 0.02179267 |
| ZBTB17 | 1.38 | 1.15 | 1.65 | 0.00040177 | 0.0032868 |
| ACTN4 | 1.26 | 1.09 | 1.46 | 0.00140888 | 0.0082955 |
| ADA | 1.24 | 1.06 | 1.46 | 0.00828292 | 0.02909893 |
| ADAM23 | 1.27 | 1.06 | 1.51 | 0.00772558 | 0.02746875 |
| ADGRE2 | 1.36 | 1.15 | 1.62 | 0.00044942 | 0.00351888 |
| AGRP | 1.29 | 1.09 | 1.54 | 0.0029664 | 0.01356071 |
| AMN | 1.26 | 1.06 | 1.49 | 0.00887338 | 0.03042015 |
| ANGPTL4 | 1.59 | 1.31 | 1.93 | 3.385E-06 | 0.0001437 |
| TNFSF13 | 1.39 | 1.19 | 1.62 | 2.0673E-05 | 0.00046817 |
| ATP5IF1 | 1.76 | 1.46 | 2.12 | 3.4675E-09 | 8.507E-07 |
| B4GALT1 | 1.33 | 1.08 | 1.64 | 0.00626349 | 0.02341654 |
| BACH1 | 1.43 | 1.21 | 1.7 | 4.039E-05 | 0.00072636 |
| BSG | 1.26 | 1.09 | 1.46 | 0.00197745 | 0.01014217 |
| BTN2A1 | 1.21 | 1.04 | 1.42 | 0.01342476 | 0.04125522 |
| BTN3A2 | 1.35 | 1.14 | 1.58 | 0.00032566 | 0.00295907 |
| C1QA | 1.27 | 1.06 | 1.53 | 0.00941998 | 0.03158592 |
| CCL23 | 1.26 | 1.05 | 1.52 | 0.01189616 | 0.03809273 |
| CCL7 | 1.41 | 1.16 | 1.7 | 0.00041704 | 0.00335458 |
| CD22 | 0.8 | 0.67 | 0.95 | 0.01081377 | 0.03545182 |
| CD4 | 1.37 | 1.15 | 1.63 | 0.00034439 | 0.00303559 |
| CD40 | 1.33 | 1.14 | 1.56 | 0.00023854 | 0.00234089 |

| | | | | | |
|---|---|---|---|---|---|
| CD48 | 1.22 | 1.04 | 1.44 | 0.01485175 | 0.04470711 |
| CDSN | 1.3 | 1.11 | 1.52 | 0.00135394 | 0.00803626 |
| CHRDL1 | 1.37 | 1.15 | 1.63 | 0.00047778 | 0.00366299 |
| CKAP4 | 1.64 | 1.38 | 1.96 | 4.4933E-08 | 6.6142E-06 |
| CLEC4D | 1.26 | 1.05 | 1.51 | 0.01249179 | 0.03945906 |
| CLEC7A | 1.25 | 1.05 | 1.48 | 0.01248579 | 0.03945906 |
| COLEC12 | 1.3 | 1.09 | 1.55 | 0.00323686 | 0.01469006 |
| CRELD2 | 1.3 | 1.12 | 1.51 | 0.0007069 | 0.00483981 |
| CRIM1 | 1.46 | 1.24 | 1.73 | 5.9336E-06 | 0.00022821 |
| CTSC | 1.25 | 1.05 | 1.49 | 0.01124945 | 0.03671661 |
| CTSO | 1.31 | 1.11 | 1.56 | 0.00181317 | 0.00962398 |
| CXADR | 1.32 | 1.13 | 1.56 | 0.00071358 | 0.00486291 |
| CXCL10 | 1.43 | 1.17 | 1.75 | 0.00040415 | 0.0032868 |
| CXCL14 | 1.27 | 1.09 | 1.47 | 0.00182114 | 0.00962398 |
| LAMP3 | 1.27 | 1.06 | 1.53 | 0.0098551 | 0.03267277 |
| DECR1 | 1.41 | 1.21 | 1.64 | 9.0563E-06 | 0.00029624 |
| DFFA | 1.45 | 1.22 | 1.72 | 3.0079E-05 | 0.00059977 |
| DNPH1 | 1.25 | 1.06 | 1.48 | 0.00857585 | 0.02977277 |
| EGLN1 | 1.29 | 1.07 | 1.56 | 0.00685268 | 0.02515497 |
| EIF4G1 | 1.33 | 1.12 | 1.59 | 0.00156527 | 0.00865087 |
| ENAH | 1.38 | 1.15 | 1.66 | 0.00041973 | 0.00335787 |
| EPCAM | 1.29 | 1.1 | 1.51 | 0.00212369 | 0.01060275 |
| ESM1 | 1.24 | 1.04 | 1.48 | 0.01669002 | 0.04903397 |
| FABP1 | 1.25 | 1.06 | 1.47 | 0.00950356 | 0.03179371 |
| FCAR | 1.35 | 1.13 | 1.6 | 0.00088282 | 0.00577563 |
| FCRL2 | 1.29 | 1.09 | 1.51 | 0.0022852 | 0.01099285 |
| FST | 1.35 | 1.15 | 1.59 | 0.00032777 | 0.00296003 |
| FSTL3 | 1.57 | 1.3 | 1.9 | 3.4167E-06 | 0.0001437 |
| FXYD5 | 1.28 | 1.1 | 1.5 | 0.00176081 | 0.00941321 |
| GBP2 | 1.44 | 1.21 | 1.72 | 4.3692E-05 | 0.00074937 |
| CSF3 | 1.22 | 1.05 | 1.43 | 0.00940916 | 0.03158592 |
| GLOD4 | 1.41 | 1.19 | 1.67 | 8.0101E-05 | 0.00117908 |
| GOPC | 1.24 | 1.06 | 1.45 | 0.00714082 | 0.02602538 |
| HCLS1 | 1.36 | 1.12 | 1.66 | 0.00194774 | 0.01005991 |
| HEXIM1 | 1.57 | 1.33 | 1.85 | 1.2219E-07 | 1.3164E-05 |
| HPCAL1 | 1.31 | 1.11 | 1.55 | 0.00153027 | 0.00863051 |
| HSPA1A | 1.47 | 1.21 | 1.78 | 7.9877E-05 | 0.00117908 |
| ICA1 | 1.25 | 1.06 | 1.49 | 0.00854588 | 0.02973887 |
| ICAM4 | 1.24 | 1.04 | 1.48 | 0.01598425 | 0.04743714 |
| IFNGR1 | 1.41 | 1.21 | 1.65 | 1.5397E-05 | 0.00040472 |

| | | | | | |
|---|---|---|---|---|---|
| IKBKG | 1.23 | 1.04 | 1.46 | 0.01618824 | 0.04794587 |
| IL12RB1 | 1.26 | 1.07 | 1.48 | 0.00608911 | 0.02313356 |
| IL15RA | 1.3 | 1.1 | 1.54 | 0.00213786 | 0.01060275 |
| IL16 | 1.4 | 1.14 | 1.7 | 0.00101366 | 0.0063765 |
| IL17C | 1.24 | 1.06 | 1.46 | 0.00820997 | 0.02892616 |
| IL18R1 | 1.39 | 1.17 | 1.67 | 0.0002609 | 0.00249383 |
| IL1R2 | 1.29 | 1.07 | 1.55 | 0.00672601 | 0.02475356 |
| IL1RN | 1.32 | 1.1 | 1.58 | 0.00324339 | 0.01469006 |
| IL3RA | 1.3 | 1.11 | 1.52 | 0.00117407 | 0.00717106 |
| IL4R | 1.33 | 1.11 | 1.58 | 0.00153854 | 0.00864401 |
| IL15 | 1.58 | 1.32 | 1.88 | 5.3656E-07 | 3.434E-05 |
| IL18 | 1.47 | 1.24 | 1.74 | 7.7226E-06 | 0.00026436 |
| IL20 | 1.22 | 1.05 | 1.4 | 0.00723029 | 0.026279 |
| IL24 | 1.28 | 1.12 | 1.46 | 0.00026061 | 0.00249383 |
| IL6 | 1.34 | 1.12 | 1.61 | 0.00144522 | 0.00838168 |
| ITGB6 | 1.52 | 1.26 | 1.84 | 1.2507E-05 | 0.00035406 |
| ITM2A | 1.26 | 1.06 | 1.48 | 0.00765266 | 0.02727534 |
| JUN | 1.54 | 1.29 | 1.83 | 1.8601E-06 | 9.4416E-05 |
| KRT19 | 1.71 | 1.37 | 2.14 | 2.3469E-06 | 0.00011038 |
| KYNU | 1.25 | 1.05 | 1.49 | 0.01361253 | 0.04160018 |
| LAIR1 | 1.29 | 1.09 | 1.53 | 0.00358166 | 0.01592811 |
| LAMA4 | 1.31 | 1.11 | 1.56 | 0.00176285 | 0.00941321 |
| LAP3 | 1.38 | 1.15 | 1.65 | 0.00045676 | 0.00355743 |
| LGALS9 | 1.56 | 1.3 | 1.86 | 1.3973E-06 | 7.6179E-05 |
| LHPP | 1.25 | 1.06 | 1.47 | 0.00725307 | 0.02629264 |
| LIFR | 1.36 | 1.14 | 1.63 | 0.00066875 | 0.00462162 |
| LILRB4 | 1.37 | 1.16 | 1.61 | 0.00016752 | 0.00191157 |
| LSP1 | 1.31 | 1.07 | 1.59 | 0.00894895 | 0.03047749 |
| LTA | 0.78 | 0.64 | 0.94 | 0.01099344 | 0.03596075 |
| LTBR | 1.37 | 1.16 | 1.62 | 0.00019294 | 0.00210371 |
| LY6D | 1.32 | 1.13 | 1.54 | 0.00054125 | 0.00399221 |
| MAPK9 | 1.59 | 1.34 | 1.89 | 1.5831E-07 | 1.4565E-05 |
| MATN2 | 1.33 | 1.11 | 1.59 | 0.00171974 | 0.00927273 |
| CSF1 | 1.34 | 1.11 | 1.61 | 0.00194439 | 0.01005991 |
| MEPE | 1.37 | 1.11 | 1.69 | 0.00372322 | 0.01626286 |
| METAP1D | 1.27 | 1.1 | 1.46 | 0.00098918 | 0.00630337 |
| MILR1 | 1.27 | 1.08 | 1.5 | 0.00433859 | 0.01829916 |
| NBN | 1.38 | 1.15 | 1.66 | 0.0005975 | 0.00431136 |
| NFATC3 | 1.21 | 1.04 | 1.41 | 0.01286138 | 0.04019524 |
| NPPC | 1.3 | 1.11 | 1.53 | 0.00151906 | 0.00860024 |

| | | | | | |
|---|---|---|---|---|---|
| NUB1 | 1.38 | 1.17 | 1.63 | 0.00014206 | 0.00171407 |
| NUDC | 1.47 | 1.21 | 1.79 | 9.2435E-05 | 0.0012852 |
| OMD | 1.36 | 1.13 | 1.64 | 0.00136844 | 0.00808973 |
| TNFRSF11B | 1.32 | 1.11 | 1.58 | 0.00188779 | 0.00988907 |
| OSCAR | 1.28 | 1.07 | 1.54 | 0.00626774 | 0.02341654 |
| PADI2 | 1.34 | 1.15 | 1.56 | 0.00016944 | 0.00191854 |
| PARP1 | 1.38 | 1.17 | 1.62 | 0.0001389 | 0.0017039 |
| PCDH1 | 1.21 | 1.06 | 1.37 | 0.00465103 | 0.01928539 |
| PLAUR | 1.46 | 1.2 | 1.77 | 0.00016305 | 0.00187507 |
| PGF | 1.27 | 1.08 | 1.5 | 0.00419688 | 0.01783788 |
| PRDX3 | 1.37 | 1.17 | 1.6 | 0.00010213 | 0.00137922 |
| PRDX5 | 1.27 | 1.05 | 1.54 | 0.01575614 | 0.04692807 |
| PREB | 1.58 | 1.36 | 1.85 | 5.3924E-09 | 1.1339E-06 |
| PRELP | 1.4 | 1.15 | 1.7 | 0.00062333 | 0.00441127 |
| PRKAB1 | 1.23 | 1.04 | 1.45 | 0.01458962 | 0.04409841 |
| PROK1 | 1.68 | 1.42 | 1.99 | 1.6555E-09 | 6.0923E-07 |
| PSIP1 | 1.67 | 1.37 | 2.04 | 2.9237E-07 | 2.1518E-05 |
| PTX3 | 1.29 | 1.06 | 1.56 | 0.00985055 | 0.03267277 |
| AGER | 1.46 | 1.22 | 1.76 | 4.9311E-05 | 0.0007805 |
| TNFRSF11A | 1.33 | 1.12 | 1.57 | 0.00103188 | 0.00646354 |
| SAMD9L | 1.29 | 1.08 | 1.54 | 0.00552269 | 0.02191384 |
| SCGB1A1 | 1.32 | 1.09 | 1.59 | 0.00443118 | 0.01853041 |
| SCRN1 | 1.33 | 1.13 | 1.56 | 0.00044188 | 0.00349413 |
| SERPINB8 | 1.39 | 1.15 | 1.67 | 0.00052481 | 0.00392143 |
| SH2D1A | 1.26 | 1.07 | 1.48 | 0.00475569 | 0.019664 |
| SIGLEC1 | 1.32 | 1.11 | 1.56 | 0.00126542 | 0.0076654 |
| SIGLEC10 | 1.3 | 1.08 | 1.56 | 0.005025 | 0.02048975 |
| SIRPB1 | 1.31 | 1.1 | 1.56 | 0.00222092 | 0.01078944 |
| SPINK4 | 1.27 | 1.07 | 1.5 | 0.00707776 | 0.02591658 |
| SRPK2 | 1.48 | 1.23 | 1.77 | 2.3732E-05 | 0.00052139 |
| TPP1 | 1.33 | 1.12 | 1.58 | 0.00134615 | 0.00803626 |
| TRAF2 | 1.27 | 1.07 | 1.5 | 0.00626545 | 0.02341654 |
| TRIM21 | 1.39 | 1.16 | 1.68 | 0.00047607 | 0.00366299 |
| TRIM5 | 1.23 | 1.05 | 1.45 | 0.01268097 | 0.0399429 |
| CKMT1B | 1.27 | 1.07 | 1.51 | 0.00535499 | 0.02159603 |
| CKMT1A | 1.27 | 1.07 | 1.51 | 0.00535499 | 0.02159603 |
| YTHDF3 | 1.34 | 1.14 | 1.57 | 0.00033679 | 0.00300137 |
| ACVRL1 | 1.32 | 1.14 | 1.54 | 0.00031997 | 0.00292549 |
| AGR2 | 1.52 | 1.25 | 1.85 | 2.6618E-05 | 0.00056785 |
| AKT1S1 | 1.3 | 1.1 | 1.52 | 0.00182411 | 0.00962398 |

| | | | | | |
|---|---|---|---|---|---|
| AMFR | 1.39 | 1.19 | 1.63 | 3.0152E-05 | 0.00059977 |
| ANXA5 | 1.36 | 1.19 | 1.56 | 6.0634E-06 | 0.00022821 |
| ARID4B | 1.53 | 1.27 | 1.85 | 6.3376E-06 | 0.00022821 |
| ASGR1 | 1.34 | 1.11 | 1.62 | 0.00221054 | 0.01077454 |
| ATP6V1F | 1.27 | 1.08 | 1.49 | 0.00364656 | 0.01616789 |
| BAG3 | 1.64 | 1.35 | 1.98 | 3.3402E-07 | 2.2349E-05 |
| BCAM | 1.39 | 1.16 | 1.67 | 0.00036511 | 0.00316142 |
| BST1 | 1.3 | 1.07 | 1.57 | 0.00726977 | 0.02629264 |
| CASP1 | 1.31 | 1.1 | 1.56 | 0.00209318 | 0.01058817 |
| CASP10 | 1.26 | 1.06 | 1.49 | 0.00965682 | 0.03216026 |
| CD300C | 1.35 | 1.14 | 1.6 | 0.0006152 | 0.00437473 |
| CD300LG | 1.28 | 1.09 | 1.5 | 0.00220645 | 0.01077454 |
| CD74 | 1.33 | 1.12 | 1.57 | 0.00114208 | 0.00703404 |
| CD99 | 1.29 | 1.11 | 1.51 | 0.00100512 | 0.00634992 |
| CD99L2 | 1.26 | 1.08 | 1.47 | 0.00371323 | 0.01626286 |
| CERT1 | 1.26 | 1.05 | 1.51 | 0.01474898 | 0.04448872 |
| CALCA | 1.76 | 1.43 | 2.16 | 5.7917E-08 | 7.7503E-06 |
| CLEC11A | 1.27 | 1.08 | 1.5 | 0.00498036 | 0.02038879 |
| CLEC14A | 1.32 | 1.13 | 1.53 | 0.00037177 | 0.00320029 |
| CLPP | 1.28 | 1.09 | 1.49 | 0.00234878 | 0.01122536 |
| CLSPN | 1.2 | 1.06 | 1.36 | 0.00429154 | 0.01815272 |
| CPPED1 | 1.3 | 1.1 | 1.53 | 0.00210171 | 0.01059492 |
| CRIP2 | 1.41 | 1.2 | 1.66 | 3.3608E-05 | 0.00063851 |
| CTSS | 1.32 | 1.12 | 1.56 | 0.00084649 | 0.00556263 |
| CX3CL1 | 1.36 | 1.15 | 1.61 | 0.0004034 | 0.0032868 |
| CXCL13 | 1.31 | 1.11 | 1.55 | 0.00114798 | 0.00704094 |
| DBI | 1.33 | 1.13 | 1.57 | 0.00064436 | 0.00449526 |
| DRAXIN | 1.41 | 1.19 | 1.68 | 9.9946E-05 | 0.00136223 |
| DSG2 | 1.31 | 1.11 | 1.54 | 0.00145054 | 0.00838168 |
| EBAG9 | 1.34 | 1.14 | 1.58 | 0.00040012 | 0.0032868 |
| ECE1 | 1.24 | 1.05 | 1.46 | 0.01274963 | 0.0399429 |
| EFNA1 | 1.38 | 1.18 | 1.63 | 9.2548E-05 | 0.0012852 |
| EFNA4 | 1.26 | 1.07 | 1.48 | 0.00530071 | 0.02147762 |
| ENO1 | 1.3 | 1.09 | 1.54 | 0.00356792 | 0.01591508 |
| EZR | 1.64 | 1.36 | 1.96 | 1.296E-07 | 1.3164E-05 |
| F11R | 1.26 | 1.07 | 1.48 | 0.00481692 | 0.01986135 |
| FABP5 | 1.38 | 1.16 | 1.64 | 0.00022083 | 0.00222641 |
| FGR | 1.34 | 1.12 | 1.62 | 0.001634 | 0.00890832 |
| FKBP4 | 1.36 | 1.12 | 1.66 | 0.00197085 | 0.01014217 |
| FKBP7 | 1.35 | 1.16 | 1.58 | 0.00011104 | 0.00143382 |

| | | | | | |
|---|---|---|---|---|---|
| FOSB | 1.42 | 1.23 | 1.63 | 8.1922E-07 | 5.0245E-05 |
| GBP4 | 1.26 | 1.06 | 1.49 | 0.00901307 | 0.0304994 |
| GGT5 | 1.22 | 1.04 | 1.44 | 0.01500232 | 0.04497641 |
| GPKOW | 1.55 | 1.29 | 1.87 | 2.4619E-06 | 0.00011038 |
| GRN | 1.34 | 1.13 | 1.58 | 0.00054242 | 0.00399221 |
| GSTP1 | 1.31 | 1.11 | 1.54 | 0.0012794 | 0.00771834 |
| HAVCR2 | 1.31 | 1.09 | 1.56 | 0.00380005 | 0.016452 |
| HMOX2 | 1.44 | 1.21 | 1.71 | 4.8419E-05 | 0.0007747 |
| HNMT | 1.28 | 1.09 | 1.49 | 0.00227543 | 0.01098174 |
| IDI2 | 1.37 | 1.16 | 1.61 | 0.00022734 | 0.00226116 |
| IGFBP4 | 1.39 | 1.16 | 1.65 | 0.00027692 | 0.002613 |
| ILKAP | 1.33 | 1.12 | 1.58 | 0.0009941 | 0.0063074 |
| IMPA1 | 1.33 | 1.13 | 1.57 | 0.00048189 | 0.00367534 |
| ING1 | 1.57 | 1.32 | 1.87 | 3.2486E-07 | 2.2349E-05 |
| IFNL1 | 1.3 | 1.09 | 1.55 | 0.00340037 | 0.01530687 |
| IL34 | 1.34 | 1.15 | 1.55 | 0.00011413 | 0.0014514 |
| KRT14 | 1.22 | 1.04 | 1.42 | 0.01234997 | 0.03917922 |
| LAYN | 1.3 | 1.1 | 1.53 | 0.00156915 | 0.00865087 |
| LBR | 1.28 | 1.06 | 1.55 | 0.00889712 | 0.03042015 |
| LRPAP1 | 1.25 | 1.07 | 1.45 | 0.00420499 | 0.01783788 |
| MAD1L1 | 1.49 | 1.23 | 1.8 | 4.5058E-05 | 0.00074937 |
| MASP1 | 1.27 | 1.07 | 1.51 | 0.00733634 | 0.02641715 |
| MATN3 | 1.6 | 1.34 | 1.9 | 2.1836E-07 | 1.7761E-05 |
| MAX | 1.3 | 1.1 | 1.54 | 0.00185486 | 0.00975124 |
| CCL2 | 1.39 | 1.14 | 1.68 | 0.00084152 | 0.00555481 |
| MIF | 1.32 | 1.11 | 1.57 | 0.00145086 | 0.00838168 |
| MMP13 | 1.25 | 1.07 | 1.47 | 0.00559651 | 0.02196815 |
| MMP3 | 1.36 | 1.15 | 1.61 | 0.00030756 | 0.00282959 |
| MSR1 | 1.29 | 1.08 | 1.53 | 0.0041177 | 0.01756884 |
| MUC13 | 1.28 | 1.08 | 1.52 | 0.00400154 | 0.01712286 |
| NOS1 | 1.46 | 1.22 | 1.75 | 2.7037E-05 | 0.00056856 |
| NPM1 | 1.44 | 1.19 | 1.75 | 0.000162 | 0.00187507 |
| NUDT5 | 1.49 | 1.26 | 1.75 | 2.1238E-06 | 0.00010421 |
| PAMR1 | 0.8 | 0.68 | 0.94 | 0.00735276 | 0.02641715 |
| PDCD5 | 1.45 | 1.23 | 1.71 | 1.0226E-05 | 0.00032082 |
| CD274 | 1.42 | 1.2 | 1.67 | 3.937E-05 | 0.00072442 |
| PEBP1 | 1.25 | 1.07 | 1.46 | 0.00604595 | 0.02306721 |
| PFDN2 | 1.48 | 1.24 | 1.77 | 1.5842E-05 | 0.00040912 |
| PHOSPHO1 | 1.48 | 1.22 | 1.8 | 8.3958E-05 | 0.00122363 |
| PIK3IP1 | 1.25 | 1.06 | 1.46 | 0.00774911 | 0.02748598 |

| | | | | | |
|---|---|---|---|---|---|
| PLAU | 1.37 | 1.15 | 1.64 | 0.00048453 | 0.0036764 |
| PLIN1 | 1.32 | 1.12 | 1.56 | 0.00081913 | 0.00545595 |
| PPCDC | 1.26 | 1.07 | 1.49 | 0.00523753 | 0.02129738 |
| PPP3R1 | 1.43 | 1.2 | 1.72 | 9.4956E-05 | 0.00130631 |
| PRDX1 | 1.42 | 1.22 | 1.66 | 1.0461E-05 | 0.00032082 |
| PSME1 | 1.3 | 1.1 | 1.54 | 0.00240708 | 0.01135647 |
| PSME2 | 1.33 | 1.12 | 1.58 | 0.00091015 | 0.00590194 |
| SPINK1 | 1.31 | 1.07 | 1.61 | 0.00897645 | 0.03047749 |
| PTK7 | 1.29 | 1.09 | 1.54 | 0.0037642 | 0.01634486 |
| PTPRN2 | 1.21 | 1.05 | 1.4 | 0.01046602 | 0.03462018 |
| PVR | 1.25 | 1.05 | 1.48 | 0.01186261 | 0.03809273 |
| PXN | 1.38 | 1.14 | 1.68 | 0.00093368 | 0.00598246 |
| RAB6B | 0.79 | 0.66 | 0.95 | 0.013104 | 0.04069427 |
| RBKS | 1.28 | 1.08 | 1.52 | 0.00437597 | 0.01838325 |
| RELT | 1.29 | 1.09 | 1.54 | 0.00375485 | 0.01634486 |
| RGMB | 1.21 | 1.03 | 1.41 | 0.01647446 | 0.04859802 |
| RWDD1 | 1.34 | 1.14 | 1.58 | 0.00036485 | 0.00316142 |
| SCARB1 | 1.18 | 1.04 | 1.35 | 0.01291614 | 0.04028084 |
| SCARB2 | 1.35 | 1.16 | 1.58 | 0.00014049 | 0.00170906 |
| SCARF2 | 1.25 | 1.06 | 1.49 | 0.00835515 | 0.02921325 |
| SERPINB1 | 1.42 | 1.17 | 1.72 | 0.00033847 | 0.00300137 |
| SERPINB6 | 1.34 | 1.14 | 1.57 | 0.00029916 | 0.00277619 |
| SETMAR | 1.47 | 1.22 | 1.75 | 2.9565E-05 | 0.00059977 |
| SNCG | 1.32 | 1.1 | 1.57 | 0.00245866 | 0.01156278 |
| SOD2 | 1.31 | 1.1 | 1.57 | 0.00224797 | 0.01088488 |
| SPOCK1 | 1.32 | 1.11 | 1.57 | 0.00134914 | 0.00803626 |
| SSB | 1.35 | 1.14 | 1.6 | 0.00053658 | 0.00398915 |
| STAMBP | 1.33 | 1.13 | 1.57 | 0.00049481 | 0.00373517 |
| STC1 | 1.44 | 1.21 | 1.71 | 3.3834E-05 | 0.00063851 |
| SUSD2 | 1.25 | 1.05 | 1.48 | 0.01078393 | 0.0354329 |
| TARBP2 | 1.23 | 1.05 | 1.45 | 0.01128979 | 0.03675521 |
| TBCC | 1.28 | 1.08 | 1.52 | 0.00531104 | 0.02147762 |
| TDGF1 | 1.22 | 1.04 | 1.43 | 0.01672218 | 0.04903397 |
| TDRKH | 1.26 | 1.08 | 1.47 | 0.00302863 | 0.0138023 |
| THBS2 | 1.4 | 1.15 | 1.7 | 0.00081329 | 0.00545595 |
| THY1 | 1.22 | 1.04 | 1.42 | 0.01208899 | 0.03843412 |
| TIGAR | 1.39 | 1.17 | 1.65 | 0.00018826 | 0.00206809 |
| TMSB10 | 1.44 | 1.19 | 1.74 | 0.00021715 | 0.00220441 |
| TNFRSF1B | 1.36 | 1.15 | 1.61 | 0.00038308 | 0.00323163 |
| TNFRSF10A | 1.47 | 1.23 | 1.76 | 2.8531E-05 | 0.00059152 |

| | | | | | |
|---|---|---|---|---|---|
| TNFRSF10B | 1.43 | 1.21 | 1.7 | 4.4282E-05 | 0.00074937 |
| TNFRSF1A | 1.39 | 1.18 | 1.64 | 7.5607E-05 | 0.00113564 |
| TNFRSF21 | 1.23 | 1.05 | 1.44 | 0.01049631 | 0.03464253 |
| TNFRSF6B | 1.42 | 1.17 | 1.73 | 0.00039931 | 0.0032868 |
| TPPP3 | 1.21 | 1.06 | 1.39 | 0.00552649 | 0.02191384 |
| MAPT | 1.24 | 1.05 | 1.47 | 0.0130345 | 0.04056404 |
| TXLNA | 1.38 | 1.17 | 1.64 | 0.00021018 | 0.00220029 |
| TXNRD1 | 1.36 | 1.14 | 1.61 | 0.00044389 | 0.00349413 |
| ULBP2 | 1.36 | 1.13 | 1.64 | 0.00108178 | 0.0067189 |
| VCAN | 1.3 | 1.08 | 1.56 | 0.00555086 | 0.02191384 |
| VSIG4 | 1.57 | 1.26 | 1.94 | 4.7404E-05 | 0.0007668 |
| VSTM1 | 1.24 | 1.04 | 1.48 | 0.0139429 | 0.04240486 |
| WARS1 | 1.41 | 1.17 | 1.69 | 0.00029987 | 0.00277619 |
| XRCC4 | 1.38 | 1.16 | 1.64 | 0.00027916 | 0.00261738 |
| ABL1 | 1.31 | 1.11 | 1.55 | 0.00156417 | 0.00865087 |
| AIF1 | 1.48 | 1.24 | 1.78 | 1.94E-05 | 0.0004606 |
| AKR1B1 | 1.24 | 1.06 | 1.45 | 0.00635948 | 0.02369912 |
| AMBP | 1.22 | 1.04 | 1.44 | 0.0136218 | 0.04160018 |
| APBB1IP | 1.36 | 1.12 | 1.66 | 0.0019007 | 0.00992136 |
| APEX1 | 1.26 | 1.04 | 1.53 | 0.01666005 | 0.04903397 |
| AREG | 1.39 | 1.15 | 1.68 | 0.00080264 | 0.00541963 |
| ATOX1 | 1.25 | 1.05 | 1.48 | 0.01192304 | 0.03809273 |
| ATP6AP2 | 1.24 | 1.06 | 1.45 | 0.00592173 | 0.02281881 |
| CALB1 | 1.42 | 1.2 | 1.7 | 7.2963E-05 | 0.00110724 |
| CAPG | 1.81 | 1.45 | 2.26 | 1.3414E-07 | 1.3164E-05 |
| CD300E | 1.42 | 1.18 | 1.72 | 0.00020351 | 0.00215517 |
| CD300LF | 1.26 | 1.04 | 1.51 | 0.01514101 | 0.04529993 |
| CD302 | 1.35 | 1.15 | 1.6 | 0.00033352 | 0.00299357 |
| CD38 | 1.3 | 1.09 | 1.56 | 0.00383046 | 0.01653503 |
| CDC37 | 1.3 | 1.11 | 1.53 | 0.00149105 | 0.00856199 |
| CDKN1A | 1.36 | 1.16 | 1.6 | 0.00015084 | 0.00179065 |
| CEACAM5 | 1.27 | 1.07 | 1.5 | 0.0059102 | 0.02281881 |
| CEP85 | 1.26 | 1.07 | 1.48 | 0.00599491 | 0.02304048 |
| CNPY4 | 1.33 | 1.14 | 1.54 | 0.00022426 | 0.00224562 |
| COX5B | 1.72 | 1.45 | 2.04 | 6.4439E-10 | 4.7427E-07 |
| KRT18 | 1.49 | 1.24 | 1.79 | 2.5263E-05 | 0.00054687 |
| DAB2 | 1.3 | 1.1 | 1.52 | 0.00149892 | 0.00856199 |
| DCTN2 | 1.42 | 1.21 | 1.67 | 1.2153E-05 | 0.00035077 |
| DDAH1 | 1.46 | 1.25 | 1.7 | 1.4814E-06 | 7.7878E-05 |
| DLL1 | 1.38 | 1.16 | 1.63 | 0.00023219 | 0.00229387 |

| | | | | | |
|---|---|---|---|---|---|
| DNAJB1 | 1.31 | 1.11 | 1.55 | 0.00176498 | 0.00941321 |
| DPY30 | 1.61 | 1.33 | 1.95 | 9.1907E-07 | 5.2034E-05 |
| DTX3 | 1.28 | 1.1 | 1.5 | 0.00202389 | 0.01030854 |
| EDA2R | 1.3 | 1.08 | 1.58 | 0.00648108 | 0.02403061 |
| ELOA | 1.48 | 1.24 | 1.77 | 2.0583E-05 | 0.00046817 |
| EPHA2 | 1.38 | 1.17 | 1.61 | 9.0066E-05 | 0.0012852 |
| EPS8L2 | 1.48 | 1.24 | 1.77 | 1.2827E-05 | 0.00035435 |
| ERP44 | 1.22 | 1.04 | 1.42 | 0.0120852 | 0.03843412 |
| F3 | 1.43 | 1.2 | 1.69 | 3.6194E-05 | 0.0006744 |
| FOLR1 | 1.3 | 1.09 | 1.54 | 0.00370344 | 0.01626286 |
| FURIN | 1.28 | 1.07 | 1.52 | 0.00583717 | 0.02264097 |
| FUS | 1.47 | 1.21 | 1.79 | 0.0001082 | 0.00141221 |
| FXN | 1.49 | 1.27 | 1.75 | 9.0033E-07 | 5.2034E-05 |
| GALNT2 | 1.32 | 1.11 | 1.56 | 0.00151838 | 0.00860024 |
| GFER | 1.43 | 1.2 | 1.7 | 4.5308E-05 | 0.00074937 |
| GPC1 | 1.37 | 1.15 | 1.64 | 0.00058627 | 0.00425116 |
| GRPEL1 | 1.4 | 1.17 | 1.68 | 0.00024295 | 0.00236832 |
| HAGH | 1.23 | 1.06 | 1.43 | 0.00554096 | 0.02191384 |
| WFDC2 | 1.32 | 1.06 | 1.65 | 0.01274028 | 0.0399429 |
| HGS | 1.24 | 1.06 | 1.45 | 0.00831828 | 0.02915359 |
| HPGDS | 1.27 | 1.09 | 1.49 | 0.00273819 | 0.01271489 |
| HS3ST3B1 | 1.29 | 1.07 | 1.55 | 0.00670775 | 0.02475356 |
| HS6ST1 | 1.28 | 1.08 | 1.53 | 0.00459916 | 0.01912421 |
| HSPB6 | 1.42 | 1.17 | 1.73 | 0.0003916 | 0.00327518 |
| HTRA2 | 1.41 | 1.18 | 1.69 | 0.00017196 | 0.00192524 |
| IDUA | 1.28 | 1.06 | 1.54 | 0.01192986 | 0.03809273 |
| IGF1R | 1.29 | 1.1 | 1.51 | 0.00200206 | 0.01023273 |
| INPP1 | 1.24 | 1.06 | 1.44 | 0.00584482 | 0.02264097 |
| INPPL1 | 1.24 | 1.06 | 1.44 | 0.0060977 | 0.02313356 |
| LAT2 | 1.28 | 1.09 | 1.51 | 0.00291675 | 0.01337527 |
| LPCAT2 | 1.32 | 1.1 | 1.58 | 0.00237507 | 0.01129542 |
| LRP1 | 1.26 | 1.07 | 1.5 | 0.00639123 | 0.02375731 |
| LTBP3 | 1.28 | 1.06 | 1.53 | 0.00920694 | 0.03108398 |
| LYAR | 1.3 | 1.11 | 1.51 | 0.00093476 | 0.00598246 |
| LYN | 1.48 | 1.22 | 1.78 | 4.3664E-05 | 0.00074937 |
| MANSC1 | 1.44 | 1.22 | 1.71 | 1.4975E-05 | 0.00040078 |
| NAMPT | 1.25 | 1.04 | 1.49 | 0.01557563 | 0.04650572 |
| NBL1 | 1.35 | 1.12 | 1.63 | 0.00145199 | 0.00838168 |
| NCS1 | 1.28 | 1.07 | 1.52 | 0.00583723 | 0.02264097 |
| NDUFS6 | 1.68 | 1.43 | 1.97 | 1.7748E-10 | 2.6125E-07 |

| | | | | | |
|---|---|---|---|---|---|
| NECTIN4 | 1.28 | 1.09 | 1.5 | 0.0021852 | 0.01077454 |
| NFKBIE | 1.4 | 1.17 | 1.67 | 0.00019714 | 0.0021182 |
| NUCB2 | 1.39 | 1.18 | 1.64 | 8.8462E-05 | 0.00127663 |
| OGFR | 1.58 | 1.29 | 1.95 | 1.2999E-05 | 0.00035435 |
| P4HB | 1.27 | 1.06 | 1.51 | 0.00791838 | 0.02801889 |
| ADCYAP1R1 | 0.75 | 0.59 | 0.94 | 0.01410598 | 0.04281238 |
| PODXL2 | 1.41 | 1.2 | 1.66 | 4.0547E-05 | 0.00072636 |
| POLR2F | 1.81 | 1.49 | 2.19 | 1.1205E-09 | 5.4977E-07 |
| PQBP1 | 1.67 | 1.39 | 2.01 | 4.0337E-08 | 6.6142E-06 |
| PRDX6 | 1.29 | 1.1 | 1.53 | 0.00234777 | 0.01122536 |
| PRKRA | 1.28 | 1.07 | 1.53 | 0.00748285 | 0.02673486 |
| PSMD9 | 1.28 | 1.09 | 1.5 | 0.00263471 | 0.01231205 |
| RABEPK | 1.3 | 1.13 | 1.49 | 0.00021226 | 0.00220029 |
| RAD23B | 1.44 | 1.22 | 1.71 | 2.2618E-05 | 0.00050445 |
| RANGAP1 | 0.79 | 0.65 | 0.95 | 0.01313565 | 0.04069539 |
| RBP2 | 1.28 | 1.07 | 1.52 | 0.00562361 | 0.02199976 |
| RRM2B | 1.35 | 1.14 | 1.59 | 0.00043582 | 0.00346771 |
| S100A4 | 1.27 | 1.08 | 1.5 | 0.00498639 | 0.02038879 |
| SEMA4C | 1.33 | 1.12 | 1.59 | 0.0010603 | 0.00661342 |
| SEPTIN9 | 1.3 | 1.1 | 1.53 | 0.00237879 | 0.01129542 |
| SF3B4 | 1.29 | 1.09 | 1.52 | 0.00266472 | 0.01241286 |
| SFTPA1 | 1.42 | 1.21 | 1.67 | 1.7606E-05 | 0.00043925 |
| SFTPA2 | 1.31 | 1.09 | 1.57 | 0.00354989 | 0.01588277 |
| SLAMF8 | 1.44 | 1.21 | 1.72 | 4.5844E-05 | 0.00074981 |
| SPINK6 | 1.46 | 1.2 | 1.79 | 0.00017412 | 0.00192711 |
| SRP14 | 1.35 | 1.12 | 1.64 | 0.00211828 | 0.01060275 |
| TACSTD2 | 1.66 | 1.39 | 1.99 | 4.136E-08 | 6.6142E-06 |
| TBL1X | 1.24 | 1.04 | 1.47 | 0.01358109 | 0.04160018 |
| TGFBR2 | 1.27 | 1.09 | 1.49 | 0.00212606 | 0.01060275 |
| TNFRSF12A | 1.42 | 1.17 | 1.73 | 0.00040668 | 0.00328918 |
| TNFRSF19 | 1.25 | 1.06 | 1.48 | 0.00714283 | 0.02602538 |
| TRIAP1 | 1.63 | 1.38 | 1.91 | 3.1554E-09 | 8.507E-07 |
| UBAC1 | 1.25 | 1.06 | 1.48 | 0.00882645 | 0.03042015 |
| USO1 | 1.32 | 1.11 | 1.57 | 0.00204249 | 0.01036739 |
| VWA1 | 1.37 | 1.11 | 1.68 | 0.00277282 | 0.01281659 |
| WFDC12 | 1.3 | 1.11 | 1.53 | 0.00143947 | 0.00838168 |
| CCN4 | 1.43 | 1.19 | 1.72 | 0.00014342 | 0.00171633 |

**Supplemental Table 3. Hazard modeling of gene expression signatures for length of hospital stay and clinical failure.**

Each row represents a gene set significantly associated with the clinical endpoints "time to hospital discharge" and "clinical failure." The outputs include hazard ratios with lower and upper 95% confidence intervals, unadjusted and Benjamini-Hochberg adjusted *P* values.

BCR, B cell receptor; CD, cluster of differentiation; DC, dendritic cell; gMDSC, granulocytic myeloid-derived suppressor cell; HLA, human leukocyte antigen; HR, hazard ratio; IFN, interferon; IRF, interferon regulatory factor; ITK, IL-2 inducible T cell kinase; LPS, lipopolysaccharide; MHC, major histocompatibility complex; mMDSC, monocytic myeloid-derived suppressor cell; NK, natural killer; PKC, protein kinase C; TLR, toll-like receptor.

| **Immune Pathways Prognostic Model Output for Time to Hospital Discharge** | | | | | |
|---|---|---|---|---|---|
| **Gene set** | **HR** | **Lower** | **Upper** | ***P* value** | ***P* adjusted** |
| Activated LPS dendritic cell surface signature | 0.68 | 0.59 | 0.79 | 9.366E-08 | 3.6951E-07 |
| Antigen presentation lipids and proteins | 1.55 | 1.36 | 1.78 | 8.7718E-11 | 1.4912E-09 |
| Antigen processing and presentation | 1.6 | 1.39 | 1.84 | 7.8087E-11 | 1.4912E-09 |
| Antiviral IFN signature | 0.68 | 0.59 | 0.79 | 1.5708E-07 | 5.3409E-07 |
| B cell surface signature | 0.75 | 0.65 | 0.88 | 0.00028738 | 0.00058626 |
| BCR signaling | 0.8 | 0.7 | 0.93 | 0.0031229 | 0.00513767 |
| CD1 and other DC receptors | 1.7 | 1.48 | 1.96 | 1.4024E-13 | 7.1523E-12 |
| CD28 costimulation | 1.24 | 1.08 | 1.44 | 0.00284866 | 0.00484272 |
| CD4 T cell surface signature Th1 stimulated | 1.18 | 1 | 1.38 | 0.04657939 | 0.05794022 |
| CD4 T cell surface signature Th2 stimulated | 1.16 | 1 | 1.34 | 0.04372198 | 0.05574552 |
| Chemokines and inflammatory molecules in myeloid cells | 0.67 | 0.58 | 0.77 | 5.2175E-08 | 2.6609E-07 |
| Chemokines and receptors | 1.2 | 1.03 | 1.4 | 0.0206199 | 0.02842202 |

| | | | | | |
|---|---|---|---|---|---|
| Complement activation | 0.7 | 0.61 | 0.81 | 7.1753E-07 | 2.1526E-06 |
| Complement and other receptors in DCs | 0.68 | 0.59 | 0.78 | 1.0144E-07 | 3.6951E-07 |
| DC surface signature | 0.65 | 0.56 | 0.76 | 1.9393E-08 | 1.4129E-07 |
| Inflammasome receptors and signaling | 0.73 | 0.63 | 0.84 | 2.0198E-05 | 5.4217E-05 |
| Innate antiviral response | 0.66 | 0.57 | 0.76 | 8.6517E-09 | 8.8247E-08 |
| Interferon alpha response | 0.81 | 0.7 | 0.94 | 0.00621086 | 0.00931629 |
| Lymphocyte generic cluster | 1.3 | 1.12 | 1.5 | 0.00056003 | 0.00102006 |
| Memory B cell surface signature | 0.8 | 0.69 | 0.94 | 0.00481602 | 0.00744294 |
| MHC TLR7 TLR8 cluster | 1.52 | 1.32 | 1.75 | 8.493E-09 | 8.8247E-08 |
| Monocyte surface signature | 0.72 | 0.62 | 0.83 | 3.9616E-06 | 1.1224E-05 |
| Myeloid cell enriched receptors and transporters | 0.78 | 0.68 | 0.9 | 0.00067971 | 0.00119535 |
| Myeloid | 0.75 | 0.65 | 0.87 | 0.00013047 | 0.00028931 |
| Naive B cell surface signature | 0.76 | 0.65 | 0.89 | 0.00044229 | 0.00083544 |
| NK cell surface signature | 1.1 | 0.94 | 1.3 | 0.22174238 | 0.24061408 |
| Plasma cell surface signature | 1.04 | 0.91 | 1.19 | 0.52470843 | 0.5352026 |
| Plasma cells B cells immunoglobulins | 1.14 | 0.98 | 1.32 | 0.09143498 | 0.10598146 |
| Proinflammatory cytokines and chemokines | 0.65 | 0.55 | 0.76 | 3.9746E-08 | 2.2522E-07 |
| Proinflammatory DC | 0.97 | 0.84 | 1.11 | 0.61497311 | 0.61497311 |
| Regulation of antigen presentation and immune response | 1.51 | 1.29 | 1.77 | 2.754E-07 | 8.7782E-07 |
| Resting DC surface signature | 1.53 | 1.31 | 1.78 | 6.1893E-08 | 2.8696E-07 |
| Signaling in T cells | 1.12 | 0.97 | 1.3 | 0.12724782 | 0.14421419 |
| T cell differentiation via ITK and PKC | 1.21 | 1.04 | 1.42 | 0.01367008 | 0.01991927 |
| T cell surface signature | 1.25 | 1.07 | 1.46 | 0.0040111 | 0.00639269 |
| T cell surface activation | 1.34 | 1.15 | 1.56 | 0.00013951 | 0.00029646 |
| TLR and inflammatory signaling | 0.68 | 0.59 | 0.78 | 9.6339E-08 | 3.6951E-07 |
| Type I interferon response | 0.66 | 0.57 | 0.76 | 1.1244E-08 | 9.5573E-08 |
| Viral sensing immunity IRF2 targets network | 0.67 | 0.58 | 0.77 | 3.2184E-08 | 2.0517E-07 |
| Cell cycle and transcription | 1.08 | 0.93 | 1.25 | 0.32089481 | 0.34095074 |

| | | | | | |
|---|---|---|---|---|---|
| Mitotic cell cycle DNA replication | 1.14 | 0.99 | 1.31 | 0.06206631 | 0.07361353 |
| Cell division E2F transcription network | 1.17 | 1.02 | 1.34 | 0.02839473 | 0.03713157 |
| Mitotic cell division | 1.1 | 0.95 | 1.27 | 0.19877143 | 0.22037702 |
| Blood coagulation | 0.76 | 0.66 | 0.89 | 0.00037178 | 0.00072927 |
| Cell movement Adhesion platelet activation | 0.86 | 0.75 | 1 | 0.05284953 | 0.06417443 |
| Platelet activation | 0.75 | 0.65 | 0.87 | 0.00012621 | 0.00028931 |
| Leukocyte activation and migration | 0.75 | 0.64 | 0.87 | 0.00011074 | 0.00026895 |
| DNA repair | 1.18 | 1.02 | 1.35 | 0.02486518 | 0.03337169 |
| Adhesion and migration chemotaxis | 1.39 | 1.18 | 1.64 | 5.8915E-05 | 0.00015023 |
| Chaperonin-mediated protein folding | 1.19 | 1.03 | 1.36 | 0.0159285 | 0.02256538 |
| Heme biosynthesis | 1.07 | 0.91 | 1.26 | 0.42415967 | 0.44147231 |

| **Immune Pathways Prognostic Model Output for Time to Hospital Discharge Adjusted for Time From Symptom Onset** | | | | | |
|---|---|---|---|---|---|
| **Gene set** | **HR** | **Lower** | **Upper** | ***P* value** | ***P* adjusted** |
| Activated LPS dendritic cell surface signature | 0.7 | 0.61 | 0.8 | 6.3115E-07 | 3.0277E-06 |
| Antigen presentation lipids and proteins | 1.49 | 1.3 | 1.72 | 2.606E-08 | 4.4301E-07 |
| Antigen processing and presentation | 1.54 | 1.34 | 1.78 | 4.5328E-09 | 1.1559E-07 |
| Antiviral IFN signature | 0.71 | 0.61 | 0.84 | 2.5596E-05 | 7.6787E-05 |
| B cell surface signature | 0.74 | 0.64 | 0.87 | 0.00016541 | 0.00040172 |
| BCR signaling | 0.82 | 0.71 | 0.94 | 0.00543195 | 0.00814792 |
| CD1 and other DC receptors | 1.65 | 1.43 | 1.9 | 1.3243E-11 | 6.7541E-10 |
| CD28 costimulation | 1.24 | 1.07 | 1.43 | 0.00361138 | 0.00575564 |
| CD4 T cell surface signature Th1 stimulated | 1.18 | 1.01 | 1.39 | 0.03707997 | 0.04727696 |
| CD4 T cell surface signature Th2 stimulated | 1.16 | 1.01 | 1.34 | 0.03875941 | 0.04821292 |
| Chemokines and inflammatory molecules in myeloid cells | 0.68 | 0.59 | 0.79 | 4.5912E-07 | 2.831E-06 |
| Chemokines and receptors | 1.22 | 1.04 | 1.42 | 0.01220287 | 0.01682017 |

| | | | | | |
|---|---|---|---|---|---|
| Complement activation | 0.73 | 0.63 | 0.85 | 3.7405E-05 | 0.0001004 |
| Complement and other receptors in DCs | 0.69 | 0.6 | 0.8 | 4.9959E-07 | 2.831E-06 |
| DC surface signature | 0.66 | 0.57 | 0.77 | 4.9292E-08 | 6.2847E-07 |
| Inflammasome receptors and signaling | 0.74 | 0.64 | 0.85 | 5.3214E-05 | 0.0001357 |
| Innate antiviral response | 0.69 | 0.59 | 0.8 | 1.8566E-06 | 7.2838E-06 |
| Interferon alpha response | 0.8 | 0.69 | 0.93 | 0.00350296 | 0.00575564 |
| Lymphocyte generic cluster | 1.3 | 1.12 | 1.5 | 0.00050827 | 0.00099699 |
| Memory B cell surface signature | 0.8 | 0.68 | 0.93 | 0.00423477 | 0.00654464 |
| MHC TLR7 TLR8 cluster | 1.47 | 1.27 | 1.7 | 1.5941E-07 | 1.355E-06 |
| Monocyte surface signature | 0.73 | 0.63 | 0.83 | 6.7114E-06 | 2.1392E-05 |
| Myeloid cell enriched receptors and transporters | 0.8 | 0.69 | 0.92 | 0.00202753 | 0.0034468 |
| Myeloid | 0.78 | 0.67 | 0.9 | 0.00088137 | 0.00160536 |
| Naive B cell surface signature | 0.75 | 0.64 | 0.87 | 0.00021555 | 0.0004997 |
| NK cell surface signature | 1.15 | 0.97 | 1.35 | 0.10032799 | 0.1112332 |
| Plasma cell surface signature | 1.08 | 0.94 | 1.23 | 0.30227561 | 0.31461339 |
| Plasma cells B cells immunoglobulins | 1.15 | 0.99 | 1.34 | 0.06361913 | 0.07210168 |
| Proinflammatory cytokines and chemokines | 0.67 | 0.57 | 0.78 | 4.4551E-07 | 2.831E-06 |
| Proinflammatory DC | 1 | 0.87 | 1.15 | 0.99640992 | 0.99640992 |
| Regulation of antigen presentation and immune response | 1.49 | 1.27 | 1.75 | 7.421E-07 | 3.1539E-06 |
| Resting DC surface signature | 1.52 | 1.3 | 1.77 | 1.1172E-07 | 1.1396E-06 |
| Signaling in T cells | 1.16 | 1 | 1.35 | 0.04992091 | 0.05987974 |
| T cell differentiation via ITK and PKC | 1.23 | 1.06 | 1.44 | 0.00823717 | 0.01166932 |
| T cell surface signature | 1.27 | 1.09 | 1.48 | 0.00197432 | 0.0034468 |
| T cell surface activation | 1.32 | 1.14 | 1.53 | 0.00027798 | 0.00060432 |
| TLR and inflammatory signaling | 0.7 | 0.61 | 0.8 | 6.5304E-07 | 3.0277E-06 |
| Type I interferon response | 0.69 | 0.59 | 0.81 | 2.4518E-06 | 8.9315E-06 |
| Viral sensing immunity IRF2 targets network | 0.7 | 0.6 | 0.82 | 5.2939E-06 | 1.7999E-05 |
| Cell cycle and transcription | 1.08 | 0.93 | 1.25 | 0.29649141 | 0.31461339 |

| | | | | | |
|---|---|---|---|---|---|
| Mitotic cell cycle DNA replication | 1.15 | 1 | 1.33 | 0.05048684 | 0.05987974 |
| Cell division E2F transcription network | 1.18 | 1.03 | 1.36 | 0.01743083 | 0.02339401 |
| Mitotic cell division | 1.1 | 0.95 | 1.28 | 0.19081956 | 0.20705953 |
| Blood coagulation | 0.77 | 0.67 | 0.9 | 0.00063048 | 0.0011909 |
| Cell movement Adhesion platelet activation | 0.87 | 0.75 | 1 | 0.05432611 | 0.0629689 |
| Platelet activation | 0.77 | 0.66 | 0.89 | 0.00045014 | 0.00091828 |
| Leukocyte activation and migration | 0.76 | 0.66 | 0.88 | 0.00028439 | 0.00060432 |
| DNA repair | 1.18 | 1.02 | 1.36 | 0.0233412 | 0.03052311 |
| Adhesion and migration chemotaxis | 1.41 | 1.2 | 1.65 | 3.4456E-05 | 9.7625E-05 |
| Chaperonin mediated protein folding | 1.21 | 1.05 | 1.4 | 0.00732316 | 0.01067089 |
| Heme biosynthesis | 1.02 | 0.85 | 1.21 | 0.8587 | 0.875874 |

| **Cell Types Prognostic Model Output for Time to Hospital Discharge** | | | | | |
|---|---|---|---|---|---|
| **Gene set** | **HR** | **Lower** | **Upper** | ***P* value** | ***P* adjusted** |
| Immature neutrophils | 0.88 | 0.75 | 1.04 | 0.12745628 | 0.16387235 |
| Eosinophils | 1.19 | 1.02 | 1.38 | 0.02283934 | 0.03745765 |
| CD14 classical monocytes | 0.96 | 0.82 | 1.11 | 0.57230484 | 0.60596983 |
| Nonclassical monocytes | 0.84 | 0.72 | 0.98 | 0.02289079 | 0.03745765 |
| CD8 T cells | 1.14 | 0.97 | 1.32 | 0.1047436 | 0.14502961 |
| CD4 T cells | 1.32 | 1.15 | 1.51 | 6.8857E-05 | 0.00017706 |
| NK cells | 1.04 | 0.89 | 1.22 | 0.63022792 | 0.63022792 |
| B cells | 1.51 | 1.32 | 1.74 | 4.1019E-09 | 1.8459E-08 |
| Plasmablasts | 1.05 | 0.92 | 1.21 | 0.47595872 | 0.53545356 |
| Megakaryocytes | 0.89 | 0.77 | 1.02 | 0.09861727 | 0.14502961 |
| Myeloid DC | 1.43 | 1.24 | 1.64 | 3.9492E-07 | 1.4217E-06 |
| Plasmacytoid DC | 1.35 | 1.17 | 1.55 | 3.2982E-05 | 9.8947E-05 |
| Neutrophils | 0.66 | 0.58 | 0.76 | 3.8838E-09 | 1.8459E-08 |
| Activated T cells | 1.25 | 1.08 | 1.45 | 0.00360684 | 0.00733982 |
| HLA DR high monocytes | 0.81 | 0.7 | 0.93 | 0.00366991 | 0.00733982 |
| HLA DR low monocytes | 0.67 | 0.59 | 0.76 | 1.0406E-09 | 1.3209E-08 |
| gMDSC | 0.64 | 0.56 | 0.74 | 1.4677E-09 | 1.3209E-08 |
| mMDSC | 1.1 | 0.95 | 1.27 | 0.21885618 | 0.26262742 |

| Cell Types Prognostic Model Output for Time to Hospital Discharge Adjusted for Time From Symptom Onset | | | | | |
|---|---|---|---|---|---|
| **Gene set** | **HR** | **Lower** | **Upper** | ***P* value** | ***P* adjusted** |
| Immature neutrophils | 0.84 | 0.72 | 1 | 0.04738676 | 0.07108014 |
| Eosinophils | 1.18 | 1.02 | 1.37 | 0.02872391 | 0.04700277 |
| CD14 classical monocytes | 0.95 | 0.82 | 1.11 | 0.52174818 | 0.52174818 |
| Nonclassical monocytes | 0.84 | 0.72 | 0.98 | 0.02410387 | 0.04338696 |
| CD8 T cells | 1.16 | 0.99 | 1.35 | 0.0655575 | 0.09077192 |
| CD4 T cells | 1.28 | 1.12 | 1.47 | 0.00042378 | 0.00108971 |
| NK cells | 1.07 | 0.91 | 1.26 | 0.40676107 | 0.43068819 |
| B cells | 1.47 | 1.28 | 1.69 | 5.4137E-08 | 3.2482E-07 |
| Plasmablasts | 1.08 | 0.93 | 1.24 | 0.31542122 | 0.35484887 |
| Megakaryocytes | 0.88 | 0.76 | 1.02 | 0.08116188 | 0.10435099 |
| Myeloid DC | 1.39 | 1.21 | 1.6 | 4.6453E-06 | 1.6723E-05 |
| Plasmacytoid DC | 1.32 | 1.15 | 1.52 | 8.0233E-05 | 0.0002407 |
| Neutrophils | 0.68 | 0.59 | 0.78 | 5.0033E-08 | 3.2482E-07 |
| Activated T cells | 1.26 | 1.08 | 1.46 | 0.0029061 | 0.00653873 |
| HLA DR high monocytes | 0.82 | 0.71 | 0.95 | 0.00632718 | 0.01265436 |
| HLA DR low monocytes | 0.69 | 0.61 | 0.79 | 8.5537E-08 | 3.8492E-07 |
| gMDSC | 0.66 | 0.57 | 0.77 | 3.2774E-08 | 3.2482E-07 |
| mMDSC | 1.09 | 0.94 | 1.27 | 0.24778518 | 0.29734222 |

| Immune Pathways Prognostic Model Output for Time to Clinical Failure | | | | | |
|---|---|---|---|---|---|
| **Gene set** | **HR** | **Lower** | **Upper** | ***P* value** | ***P* adjusted** |
| Activated LPS dendritic cell surface signature | 1.23 | 1.04 | 1.45 | 0.01840557 | 0.02681955 |
| Antigen presentation lipids and proteins | 0.78 | 0.66 | 0.92 | 0.00319117 | 0.00813747 |
| Antigen processing and presentation | 0.79 | 0.67 | 0.93 | 0.00562506 | 0.01254642 |
| Antiviral IFN signature | 1.38 | 1.19 | 1.61 | 3.6717E-05 | 0.00052748 |
| B cell surface signature | 1.35 | 1.16 | 1.58 | 0.00016774 | 0.00171094 |
| BCR signaling | 1.15 | 0.97 | 1.37 | 0.1071296 | 0.13659024 |
| CD1 and other DC receptors | 0.74 | 0.62 | 0.87 | 0.00036409 | 0.00232104 |
| CD28 costimulation | 0.81 | 0.69 | 0.96 | 0.01231365 | 0.02325912 |
| CD4 T cell surface signature Th1 stimulated | 0.84 | 0.72 | 0.99 | 0.03669806 | 0.05058382 |
| CD4 T cell surface signature Th2 stimulated | 0.85 | 0.72 | 1 | 0.05471822 | 0.0715546 |

| | | | | | |
|---|---|---|---|---|---|
| Chemokines and inflammatory molecules in myeloid cells | 1.22 | 1.05 | 1.43 | 0.00973274 | 0.01909115 |
| Chemokines and receptors | 0.81 | 0.69 | 0.96 | 0.01454874 | 0.0243394 |
| Complement activation | 1.38 | 1.16 | 1.64 | 0.00022144 | 0.00177118 |
| Complement and other receptors in DCs | 1.27 | 1.07 | 1.52 | 0.00747611 | 0.01525126 |
| DC surface signature | 1.31 | 1.11 | 1.55 | 0.00171712 | 0.00486517 |
| Inflammasome receptors and signaling | 1.33 | 1.12 | 1.58 | 0.0012889 | 0.00410836 |
| Innate antiviral response | 1.21 | 1.04 | 1.42 | 0.01610066 | 0.0251363 |
| Interferon alpha response | 1.29 | 1.1 | 1.52 | 0.00144006 | 0.00432018 |
| Lymphocyte generic cluster | 0.74 | 0.63 | 0.88 | 0.00060936 | 0.00310773 |
| Memory B cell surface signature | 1.27 | 1.08 | 1.48 | 0.00317238 | 0.00813747 |
| MHC TLR7 TLR8 cluster | 0.79 | 0.66 | 0.93 | 0.00549548 | 0.01254642 |
| Monocyte surface signature | 1.4 | 1.16 | 1.7 | 0.00051831 | 0.00293709 |
| Myeloid cell enriched receptors and transporters | 1.21 | 1 | 1.47 | 0.0459916 | 0.06172557 |
| Myeloid | 1.32 | 1.12 | 1.56 | 0.00090252 | 0.00360299 |
| Naive B cell surface signature | 1.34 | 1.15 | 1.57 | 0.0002431 | 0.00177118 |
| NK cell surface signature | 0.82 | 0.7 | 0.97 | 0.01772003 | 0.02658005 |
| Plasma cell surface signature | 1.01 | 0.86 | 1.19 | 0.92858364 | 0.94715532 |
| Plasma cells B cells immunoglobulins | 0.92 | 0.79 | 1.07 | 0.27340127 | 0.3031188 |
| Proinflammatory cytokines and chemokines | 1.42 | 1.2 | 1.67 | 4.1371E-05 | 0.00052748 |
| Proinflammatory DC | 1.07 | 0.91 | 1.26 | 0.42991991 | 0.46650884 |
| Regulation of antigen presentation and immune response | 0.79 | 0.67 | 0.93 | 0.00565819 | 0.01254642 |
| Resting DC surface signature | 0.76 | 0.65 | 0.9 | 0.00112514 | 0.00388511 |
| Signaling in T cells | 0.83 | 0.7 | 0.97 | 0.02257645 | 0.0319833 |
| T cell differentiation via ITK and PKC | 0.76 | 0.65 | 0.9 | 0.00091841 | 0.00360299 |
| T cell surface signature | 0.77 | 0.65 | 0.9 | 0.00114268 | 0.00388511 |
| T cell surface activation | 0.75 | 0.63 | 0.88 | 0.00068214 | 0.00316264 |
| TLR and inflammatory signaling | 1.49 | 1.23 | 1.79 | 3.1129E-05 | 0.00052748 |

| | | | | | |
|---|---|---|---|---|---|
| Type I interferon response | 1.22 | 1.04 | 1.43 | 0.01479454 | 0.0243394 |
| Viral sensing immunity IRF2 targets network | 1.22 | 1.04 | 1.42 | 0.01332416 | 0.024269 |
| Cell cycle and transcription | 1 | 0.85 | 1.18 | 0.95521458 | 0.95521458 |
| Mitotic cell cycle DNA replication | 0.91 | 0.77 | 1.06 | 0.23516906 | 0.27258232 |
| Cell division E2F transcription network | 0.89 | 0.76 | 1.05 | 0.16723349 | 0.20306924 |
| Mitotic cell division | 0.99 | 0.84 | 1.16 | 0.86071996 | 0.89585139 |
| Blood coagulation | 1.26 | 1.05 | 1.51 | 0.01416788 | 0.0243394 |
| Cell movement adhesion platelet activation | 1.11 | 0.94 | 1.31 | 0.20841522 | 0.24719014 |
| Platelet activation | 1.25 | 1.04 | 1.49 | 0.01626466 | 0.0251363 |
| Leukocyte activation and migration | 1.27 | 1.07 | 1.5 | 0.00619633 | 0.0131672 |
| DNA repair | 0.88 | 0.74 | 1.03 | 0.1142495 | 0.14211524 |
| Adhesion and migration chemotaxis | 0.69 | 0.58 | 0.81 | 5.9325E-06 | 0.00030256 |
| Chaperonin mediated protein folding | 0.91 | 0.77 | 1.07 | 0.24443936 | 0.27703128 |
| Heme biosynthesis | 0.96 | 0.82 | 1.13 | 0.65245237 | 0.69323064 |

| **Immune Pathways Prognostic Model Output for Time to Clinical Failure (adjusted for time from symptom onset)** | | | | | |
|---|---|---|---|---|---|
| **Gene set** | **HR** | **Lower** | **Upper** | ***P* value** | ***P* adjusted** |
| Activated LPS dendritic cell surface signature | 1.49 | 1.24 | 1.79 | 1.8847E-05 | 8.7379E-05 |
| Antigen presentation lipids and proteins | 0.64 | 0.53 | 0.76 | 8.6684E-07 | 5.5261E-06 |
| Antigen processing and presentation | 0.64 | 0.54 | 0.75 | 2.0132E-07 | 2.1159E-06 |
| Antiviral IFN signature | 1.33 | 1.12 | 1.58 | 0.00130283 | 0.00276851 |
| B cell surface signature | 1.37 | 1.16 | 1.61 | 0.00018165 | 0.000579 |
| BCR signaling | 1.14 | 0.96 | 1.35 | 0.13326429 | 0.16991196 |
| CD1 and other DC receptors | 0.56 | 0.47 | 0.67 | 1.3466E-10 | 6.8679E-09 |
| CD28 costimulation | 0.82 | 0.69 | 0.96 | 0.01653504 | 0.02480256 |
| CD4 T cell surface signature Th1 stimulated | 0.84 | 0.72 | 0.99 | 0.03736687 | 0.05150568 |

| | | | | | |
|---|---|---|---|---|---|
| CD4 T cell surface signature Th2 stimulated | 0.85 | 0.72 | 1 | 0.04687474 | 0.06291084 |
| Chemokines and inflammatory molecules in myeloid cells | 1.42 | 1.22 | 1.66 | 7.1832E-06 | 3.6634E-05 |
| Chemokines and receptors | 0.81 | 0.68 | 0.96 | 0.01310655 | 0.02088856 |
| Complement activation | 1.34 | 1.11 | 1.61 | 0.00199058 | 0.00375998 |
| Complement and other receptors in DCs | 1.62 | 1.35 | 1.95 | 2.9042E-07 | 2.1159E-06 |
| DC surface signature | 1.68 | 1.41 | 2 | 1.0266E-08 | 1.7452E-07 |
| Inflammasome receptors and signaling | 1.34 | 1.12 | 1.59 | 0.00106493 | 0.00244646 |
| Innate antiviral response | 1.39 | 1.17 | 1.65 | 0.00021468 | 0.00059657 |
| Interferon alpha response | 1.32 | 1.12 | 1.55 | 0.0011033 | 0.00244646 |
| Lymphocyte generic cluster | 0.73 | 0.61 | 0.86 | 0.00022225 | 0.00059657 |
| Memory B cell surface signature | 1.26 | 1.08 | 1.47 | 0.00407459 | 0.00716565 |
| MHC TLR7 TLR8 cluster | 0.63 | 0.53 | 0.75 | 2.4135E-07 | 2.1159E-06 |
| Monocyte surface signature | 1.43 | 1.18 | 1.73 | 0.00026597 | 0.00067821 |
| Myeloid cell enriched receptors and transporters | 1.2 | 0.99 | 1.45 | 0.06235855 | 0.0815458 |
| Myeloid | 1.28 | 1.09 | 1.51 | 0.0030925 | 0.00563277 |
| Naive B cell surface signature | 1.36 | 1.16 | 1.6 | 0.00017795 | 0.000579 |
| NK cell surface signature | 0.8 | 0.68 | 0.95 | 0.00885436 | 0.01505241 |
| Plasma cell surface signature | 0.98 | 0.83 | 1.16 | 0.85388384 | 0.88939861 |
| Plasma cells B cells immunoglobulins | 0.91 | 0.78 | 1.06 | 0.23012519 | 0.26080855 |
| Proinflammatory cytokines and chemokines | 1.43 | 1.19 | 1.72 | 0.00017513 | 0.000579 |
| Proinflammatory DC | 1.03 | 0.87 | 1.21 | 0.76019326 | 0.82489056 |
| Regulation of antigen presentation and immune response | 0.64 | 0.54 | 0.76 | 2.5273E-07 | 2.1159E-06 |
| Resting DC surface signature | 0.6 | 0.51 | 0.71 | 3.1584E-09 | 8.0538E-08 |
| Signaling in T cells | 0.82 | 0.69 | 0.96 | 0.01436529 | 0.0222009 |
| T cell differentiation via ITK and PKC | 0.75 | 0.63 | 0.89 | 0.00143216 | 0.0029216 |
| T cell surface signature | 0.76 | 0.64 | 0.9 | 0.00160753 | 0.00315323 |
| T cell surface activation | 0.74 | 0.63 | 0.87 | 0.00041452 | 0.00100668 |

| | | | | | |
|---|---|---|---|---|---|
| TLR and inflammatory signaling | 1.5 | 1.24 | 1.81 | 2.7894E-05 | 0.00011855 |
| Type I interferon response | 1.4 | 1.17 | 1.66 | 0.00016448 | 0.000579 |
| Viral sensing immunity IRF2 targets network | 1.36 | 1.16 | 1.6 | 0.00020261 | 0.00059657 |
| Cell cycle and transcription | 1 | 0.85 | 1.18 | 0.97131324 | 0.97131324 |
| Mitotic cell cycle DNA replication | 0.91 | 0.78 | 1.07 | 0.26748722 | 0.29656191 |
| Cell division E2F transcription network | 0.89 | 0.76 | 1.05 | 0.17630031 | 0.20910036 |
| Mitotic cell division | 0.99 | 0.84 | 1.16 | 0.85452024 | 0.88939861 |
| Blood coagulation | 1.25 | 1.04 | 1.5 | 0.01732548 | 0.0252457 |
| Cell movement adhesion platelet activation | 1.13 | 0.95 | 1.33 | 0.16238551 | 0.19718241 |
| Platelet activation | 1.23 | 1.03 | 1.47 | 0.02555383 | 0.03620126 |
| Leukocyte activation and migration | 1.25 | 1.06 | 1.48 | 0.00951958 | 0.01566124 |
| DNA repair | 0.89 | 0.75 | 1.04 | 0.1425417 | 0.17730796 |
| Adhesion and migration chemotaxis | 0.68 | 0.57 | 0.8 | 2.3504E-06 | 1.3319E-05 |
| Chaperonin mediated protein folding | 0.9 | 0.76 | 1.06 | 0.20284133 | 0.23511154 |
| Heme biosynthesis | 0.99 | 0.84 | 1.17 | 0.93105601 | 0.94967713 |

| **Cell Types Prognostic Model Output for Time to Clinical Failure** | | | | | |
|---|---|---|---|---|---|
| **Gene set** | **HR** | **Lower** | **Upper** | ***P* value** | ***P* adjusted** |
| Immature neutrophils | 1.21 | 1.03 | 1.43 | 0.0207708 | 0.03738744 |
| Eosinophils | 0.79 | 0.68 | 0.93 | 0.00397908 | 0.00795816 |
| CD14 classical monocytes | 1.04 | 0.87 | 1.24 | 0.6520298 | 0.73353353 |
| Nonclassical monocytes | 1.14 | 0.95 | 1.37 | 0.15084149 | 0.19393906 |
| CD8 T cells | 0.83 | 0.7 | 0.97 | 0.02317214 | 0.03791804 |
| CD4 T cells | 0.77 | 0.65 | 0.91 | 0.00245333 | 0.00552 |
| NK cells | 0.87 | 0.74 | 1.03 | 0.10521653 | 0.14568443 |
| B cells | 0.68 | 0.57 | 0.8 | 4.5236E-06 | 2.0356E-05 |
| Plasmablasts | 1 | 0.85 | 1.16 | 0.9573181 | 0.9573181 |
| Megakaryocytes | 1.07 | 0.91 | 1.26 | 0.43973958 | 0.5276875 |
| Myeloid DC | 0.67 | 0.56 | 0.79 | 4.169E-06 | 2.0356E-05 |
| Plasmacytoid DC | 0.71 | 0.6 | 0.84 | 5.8442E-05 | 0.00017532 |
| Neutrophils | 1.61 | 1.32 | 1.96 | 2.4924E-06 | 2.0356E-05 |
| Activated T cells | 0.73 | 0.62 | 0.86 | 0.00020401 | 0.0005246 |

| | | | | | |
|---|---|---|---|---|---|
| HLA DR high monocytes | 1.2 | 1 | 1.43 | 0.04623741 | 0.06935611 |
| HLA DR low monocytes | 1.55 | 1.28 | 1.87 | 6.2733E-06 | 2.2584E-05 |
| gMDSC | 1.6 | 1.33 | 1.94 | 1.1976E-06 | 2.0356E-05 |
| mMDSC | 0.99 | 0.84 | 1.16 | 0.88517525 | 0.93724438 |

| **Cell Types Prognostic Model Output for Time to Clinical Failure Adjusted for Time From Symptom Onset** | | | | | |
|---|---|---|---|---|---|
| **Gene set** | **HR** | **Lower** | **Upper** | ***P* value** | ***P* adjusted** |
| Immature neutrophils | 1.26 | 1.07 | 1.49 | 0.0061253 | 0.01102554 |
| Eosinophils | 0.8 | 0.68 | 0.93 | 0.00469062 | 0.00938125 |
| CD14 classical monocytes | 1.05 | 0.88 | 1.26 | 0.58531115 | 0.65847505 |
| Non classical monocytes | 1.14 | 0.95 | 1.36 | 0.1631433 | 0.20975568 |
| CD8 T cells | 0.83 | 0.7 | 0.98 | 0.02366135 | 0.03871858 |
| CD4 T cells | 0.78 | 0.66 | 0.92 | 0.00384206 | 0.00864464 |
| NK cells | 0.86 | 0.73 | 1.01 | 0.0741421 | 0.10265829 |
| B cells | 0.69 | 0.58 | 0.82 | 1.6104E-05 | 5.9099E-05 |
| Plasmablasts | 0.98 | 0.84 | 1.14 | 0.77544517 | 0.82105959 |
| Megakaryocytes | 1.08 | 0.92 | 1.28 | 0.34492588 | 0.41391106 |
| Myeloid DC | 0.68 | 0.57 | 0.81 | 1.6416E-05 | 5.9099E-05 |
| Plasmacytoid DC | 0.71 | 0.59 | 0.85 | 0.00017223 | 0.0005167 |
| Neutrophils | 1.54 | 1.28 | 1.86 | 5.5963E-06 | 3.812E-05 |
| Activated T cells | 0.73 | 0.61 | 0.87 | 0.00057829 | 0.00148702 |
| HLA DR high monocytes | 1.19 | 0.99 | 1.42 | 0.0608285 | 0.09124276 |
| HLA DR low monocytes | 1.54 | 1.28 | 1.86 | 6.3533E-06 | 3.812E-05 |
| gMDSC | 1.55 | 1.29 | 1.85 | 2.2638E-06 | 3.812E-05 |
| mMDSC | 1 | 0.85 | 1.17 | 0.96191505 | 0.96191505 |

**Supplemental Table 4. Cell type specific gene sets used for RNA-seq analysis.**

CD, cluster of differentiation; gMDSC, granulocytic myeloid-derived suppressor cell; HLA, human leukocyte antigen; mMDSC, monocytic myeloid-derived suppressor cell; NK, natural killer.

| Immature neutrophils | Eosinophils | CD14+ Classical Monocytes | Nonclassical Monocytes | CD8 T-cells | CD4 T-cells | NK cells | B cells | Plasmablasts |
|---|---|---|---|---|---|---|---|---|
| AZU1 | AKAP12 | ANXA2 | CDKN1C | CCL5 | EEF1A1 | CCL4 | BANK1 | IGKC |
| BPI | AREG | CD36 | FCGR3A | CD2 | FOXP3 | CCL5 | BIRC3 | JCHAIN |
| CAMP | ATP10D | CLU | MS4A7 | CD3D | IKZF2 | CD7 | CD22 | IGHA1 |
| CEACAM8 | BHLHE40 | CPVL | LST1 | CD3E | IL2RA | CTSW | CD74 | IGLC2 |
| CRISP3 | CAMK1D | CST3 | TCF7L2 | CD3G | IL32 | CX3CR1 | CD79A | IGHGP |
| DEFA1B | CCR3 | CTSB | CSF1R | ARL4C | IL7R | FGFBP2 | FAM129C | IGHG1 |
| DEFA3 | CLC | CYP1B1 | AIF1 | CD8A | LDHB | GNLY | HLA-DPA1 | IGLC3 |
| DEFA4 | CPA3 | FCN1 | COTL1 | CD8B | RPL13 | GZMA | HLA-DPB1 | IGHG2 |
| ELANE | CSF2RB | HLA-DRA | LILRB2 | DUSP2 | RPL32 | GZMB | HLA-DQA1 | IGHA2 |
| LCN2 | FCER1A | HLA-DRB1 | SMIM25 | FGFBP2 | RPS12 | GZMH | HLA-DQB1 | IGHM |
| LTF | GATA2 | IFI27 | PECAM1 | GNLY | RPS14 | IL2RB | HLA-DRA | IGHG3 |
| MMP8 | GCSAML | LGALS1 | SIGLEC10 | GZMA | RPS18 | KLRB1 | HLA-DRB1 | HSP90B1 |
| MPO | HDC | LTA4H | HLA-DPA1 | GZMH | RPS20 | KLRD1 | IGHD | MZB1 |
| MS4A3 | MS4A2 | LYZ | WARS | GZMK | RPS27 | KLRF1 | IGHM | IGLV1-51 |
| OLFM4 | MS4A3 | MAFB | HMOX1 | IL32 | RPS6 | ARL4C | IGKC | IGHG4 |
| PGLYRP1 | RUNX1 | MPEG1 | LRRC25 | NKG7 | RPS8 | NKG7 | LINC00926 | SEC11C |
| RETN | SLC24A3 | MS4A6A | NR4A1 | RUNX3 | RTKN2 | PRF1 | MEF2C | AC244205.1 |
| RNASE2 | SLC45A3 | RNASE2 | LYN | TRAC | TCF7 | RUNX3 | MS4A1 | ITM2C |
| RNASE3 | SMPD3 | TMEM176B | CST3 | TRBC2 | TRAC | SH2D1B | PAX5 | XBP1 |
| TCN1 | SYNE1 | VCAN | SAT1 | TRGC2 | TRBC2 | SPON2 | RALGPS2 | IGLL5 |

| Megakaryo-cytes | Myeloid dendritic cells | Plasmacytoid dendritic cells | Neutrophils | Activated T cells | HLA-DR High Monocytes | HLA-DR Low Monocytes | gMDSC | mMDSC |
|---|---|---|---|---|---|---|---|---|
| AC068234.1 | ANXA2 | APP | ALDH1A2 | ANK3 | AC245128.3 | APLP2 | SAMD9L | ANKLE1 |
| CAVIN2 | CD1C | BCL11A | ALOX5AP | ARHGAP15 | ATP2B1-AS1 | C5AR1 | IL1RN | GFI1 |
| CLU | CD74 | C12orf75 | ALPL | BTBD9 | C5AR1 | CD14 | CSF3R | LRR1 |
| F13A1 | CLEC10A | C1orf186 | CD177 | CD247 | CCL2 | CD163 | CSF1 | GMNN |
| GNG11 | CPVL | CCDC50 | CXCL8 | DOCK10 | CCL3 | CD36 | GDA | CCNF |
| GPX1 | CST3 | CD74 | CXCR2 | FHIT | CD300E | CLU | HP | BUB1B |
| ITGA2B | FCER1A | FAM129C | EPHB1 | INPP4B | CDKN1A | CTSB | IFIT1 | F13A1 |
| MMD | HLA-DMA | GZMB | FAM129A | KLF12 | CLEC7A | CYP1B1 | IFIT3 | MKI67 |
| MPIG6B | HLA-DMB | HLA-DPA1 | FCGR3B | LINC-PINT | CST3 | FGR | NUDT4 | MPO |
| MTURN | HLA-DPA1 | IRF8 | FFAR2 | LINC01619 | CXCL16 | FOS | IFIT2 | STMN1 |
| MYL9 | HLA-DPB1 | ITM2C | G0S2 | LRBA | DUSP6 | FPR1 | NDST1 | MYB |
| NRGN | HLA-DQA1 | JCHAIN | GBP2 | MAML2 | EMP1 | IFI27 | IL18RAP | NUSAP1 |
| PF4 | HLA-DQB1 | LILRA4 | HIST1H2AC | PDE3B | FOSB | LGALS9 | PGLYRP1 | OIP5 |
| PPBP | HLA-DRA | MPEG1 | IFIT2 | PRKCA | FTH1 | LILRB2 | CD177 | UBASH3A |
| PRKAR2B | HLA-DRB1 | NAPSB | IFIT3 | PRKCH | HBEGF | MNDA | ZBP1 | PKNOX2 |
| RUFY1 | HLA-DRB5 | PLD4 | IFITM2 | RABGAP1L | HLA-DRA | MS4A4A | RSAD2 | GEN1 |
| SPARC | NAPSB | SERPINF1 | IL1R2 | SKAP1 | IL1B | PLAC8 | IL1R2 | WDHD1 |
| TREML1 | PEA15 | TCF4 | LIMK2 | SMYD3 | KLF4 | SERPINA1 | MCEMP1 | TIPIN |
| TUBB1 | S100A10 | TSPAN13 | MME | TSHZ2 | LGALS3 | SERPINB1 | TYROBP | NUP210 |
| VCL | SAMHD1 | UGCG | MMP9 | ZBTB20 | MAP3K8 | SIGLEC1 | SLPI | ZGRF1 |
| | | | PTGS2 | | MARCKS | SPI1 | CRISPLD2 | SPNS3 |
| | | | RGS2 | | MYADM | TMEM176A | S100A6 | PRSS57 |
| | | | S100A11 | | PHLDA1 | TMEM176B | CMPK2 | KNL1 |
| | | | S100A12 | | PPIF | VCAN | PADI4 | CDC25A |

| | | | | | | | | |
|---|---|---|---|---|---|---|---|---|
| | | | S100P | | RASGEF1B | | RAF1 | NEIL3 |
| | | | SAT1 | | SGK1 | | S100A9 | CCNE2 |
| | | | SEC14L1 | | THBD | | S100A8 | PRTN3 |
| | | | TREM1 | | ZFP36L1 | | SLC27A4 | KIF20B |
| | | | VNN2 | | | | PFKFB4 | TOX |
| | | | | | | | C5AR1 | ASF1B |
| | | | | | | | FPR1 | PIF1 |
| | | | | | | | FPR2 | CDCA3 |
| | | | | | | | ADIPOR1 | CDCA5 |
| | | | | | | | RASGRP4 | ARHGAP19 |
| | | | | | | | HCAR2 | THY1 |
| | | | | | | | CD300LF | PKMYT1 |
| | | | | | | | IP6K1 | SKA1 |
| | | | | | | | PAG1 | RAD51AP1 |
| | | | | | | | CCR1 | CYP11A1 |
| | | | | | | | NBEAL2 | STC2 |
| | | | | | | | ISG15 | MYL10 |
| | | | | | | | BST1 | CEP72 |
| | | | | | | | ABTB1 | PROM1 |
| | | | | | | | SELL | GRIA3 |
| | | | | | | | TREML2 | ELANE |
| | | | | | | | TARM1 | CNTLN |
| | | | | | | | DMXL2 | CEP78 |
| | | | | | | | LCN2 | PLK4 |
| | | | | | | | FBXL5 | BARD1 |
| | | | | | | | MXD1 | STIL |
| | | | | | | | RTP4 | FANCM |
| | | | | | | | TINAGL1 | SNAI3 |
| | | | | | | | SLC2A3 | ARHGEF39 |

| | | | | | | | | |
|---|---|---|---|---|---|---|---|---|
| | | | | | | | PYGL | NKG7 |
| | | | | | | | LYST | KLHL23 |
| | | | | | | | SLC2A6 | CDC7 |
| | | | | | | | TIMP2 | CDC6 |
| | | | | | | | FFAR2 | NDC80 |
| | | | | | | | UPP1 | ARHGAP33 |
| | | | | | | | CD33 | HAL |
| | | | | | | | CERS6 | MXD3 |
| | | | | | | | SH2D3C | MS4A3 |
| | | | | | | | MMP8 | NCAPG2 |
| | | | | | | | MMP9 | KIF11 |
| | | | | | | | DUSP6 | EFCAB11 |
| | | | | | | | RHOB | CHTF18 |
| | | | | | | | IL17RA | DMKN |
| | | | | | | | FGR | RND3 |
| | | | | | | | MMP13 | KIF15 |
| | | | | | | | OAS2 | CHAF1A |
| | | | | | | | IL1B | BCL7A |
| | | | | | | | PECAM1 | MIS18BP1 |
| | | | | | | | IRF7 | NUF2 |
| | | | | | | | CD44 | MUC13 |
| | | | | | | | TLR2 | CTSG |
| | | | | | | | LILRA6 | TCF19 |
| | | | | | | | SLFN5 | NDC1 |
| | | | | | | | PTAFR | EGFL7 |
| | | | | | | | CXCR4 | GPX3 |
| | | | | | | | SAMHD1 | H2AFX |
| | | | | | | | THBS1 | HAUS5 |
| | | | | | | | STX11 | CIT |

| | | | | | | | | |
|---|---|---|---|---|---|---|---|---|
| | | | | | | | KCTD20 | TGFBR3 |
| | | | | | | | CXCR2 | TAL1 |
| | | | | | | | CCL3 | CHDH |
| | | | | | | | EGLN3 | MCM4 |
| | | | | | | | GABBR1 | BIRC5 |
| | | | | | | | SEMA4D | MCM5 |
| | | | | | | | GSR | MTFR2 |
| | | | | | | | OSM | PLPPR3 |
| | | | | | | | ASPRV1 | DNAH8 |
| | | | | | | | KLF2 | SLC43A3 |
| | | | | | | | LRG1 | PTGER3 |
| | | | | | | | PI16 | TYMS |
| | | | | | | | SMAP2 | NSL1 |
| | | | | | | | AGRN | AURKB |
| | | | | | | | YPEL3 | HIRIP3 |
| | | | | | | | MSRB1 | CDC45 |
| | | | | | | | GRINA | POC1A |
| | | | | | | | | RAD54L |
| | | | | | | | | IL12RB1 |
| | | | | | | | | E2F8 |
| | | | | | | | | GINS1 |
| | | | | | | | | CENPU |
| | | | | | | | | POLQ |
| | | | | | | | | RRM1 |
| | | | | | | | | RRM2 |
| | | | | | | | | SHCBP1 |
| | | | | | | | | TICRR |
| | | | | | | | | TACC3 |
| | | | | | | | | ADGRL2 |

| | | | | | | | | |
|---|---|---|---|---|---|---|---|---|
| | | | | | | | | SPC24 |
| | | | | | | | | SPC25 |

**Supplemental Table 5. Sample counts per category corresponding to box and line plots.**

Sample numbers for each category plotted in Figure 4A, 6C, S9.

Abs, absolute; BL, baseline; CRP, C reactive protein; IL-6, interleukin 6; sIL6R, soluble interleukin 6 receptor; Wk, week.

| **FIG 4A** | | | | | | | | | | | | | |
|---|---|---|---|---|---|---|---|---|---|---|---|---|---|
| **Tocilizumab** | | | | | | | **Placebo** | | | | | | |
| **Serum IL-6** | **BL** | **Day 3** | **Wk 1** | **Wk 2** | **Wk 4** | **Day 60** | **Serum IL-6** | **BL** | **Day 3** | **Wk 1** | **Wk 2** | **Wk 4** | **Day 60** |
| Survivors | 150 | 148 | 127 | 105 | 74 | 32 | Survivors | 65 | 75 | 66 | 52 | 27 | 18 |
| Dead | 35 | 35 | 28 | 14 | 0 | 0 | Dead | 19 | 15 | 13 | 6 | 0 | 0 |
| **Tocilizumab** | | | | | | | **Placebo** | | | | | | |
| **sIL6R** | **BL** | **Day 3** | **Wk 1** | **Wk 2** | **Wk 4** | **Day 60** | **sIL6R** | **BL** | **Day 3** | **Wk 1** | **Wk 2** | **Wk 4** | **Day 60** |
| Survivors | 227 | 211 | 184 | 143 | 151 | 138 | Survivors | 109 | 101 | 92 | 83 | 81 | 66 |
| Dead | 56 | 48 | 31 | 17 | 0 | 0 | Dead | 27 | 20 | 15 | 8 | 0 | 0 |
| **Tocilizumab** | | | | | | | **Placebo** | | | | | | |
| **CRP** | **BL** | **Day 3** | **Wk 1** | **Wk 2** | **Wk 4** | **Day 60** | **CRP** | **BL** | **Day 3** | **Wk 1** | **Wk 2** | **Wk 4** | **Day 60** |
| Survivors | 214 | 192 | 172 | 135 | 146 | 138 | Survivors | 106 | 96 | 90 | 81 | 77 | 66 |
| Dead | 52 | 45 | 30 | 16 | 0 | 0 | Dead | 24 | 18 | 13 | 7 | 0 | 0 |

| **FIG 6C** | | | | | | | | | | | | | | |
|---|---|---|---|---|---|---|---|---|---|---|---|---|---|---|
| **Tocilizumab** | | | | | | | **Placebo** | | | | | | | |
| | **BL** | **Day 3** | **Day 7** | **Day 14** | **Day 28** | **Day 60** | | **BL** | **Day 3** | **Day 7** | **Day 14** | **Day 28** | **Day 60** | **CTRL** |
| Survivors | 216 | 200 | 183 | 57 | 143 | 101 | Survivors | 105 | 94 | 86 | 40 | 76 | 45 | 19 |
| Dead | 55 | 48 | 30 | 5 | 0 | 0 | Dead | 28 | 20 | 15 | 1 | 0 | 0 | 0 |

| FIG S9 | | | | | | | | | | | | | |
|---|---|---|---|---|---|---|---|---|---|---|---|---|---|
| **Tocilizumab** | | | | | | | **Placebo** | | | | | | |
| **Lymphocyte Pct** | **BL** | **Day 3** | **Wk 1** | **Wk 2** | **Wk 4** | **Day 60** | **Lymphocyt e Pct** | **BL** | **Day 3** | **Wk 1** | **Wk 2** | **Wk 4** | **Day 60** |
| Survivors | 225 | 188 | 190 | 147 | 102 | 137 | Survivors | 109 | 99 | 102 | 80 | 53 | 69 |
| Dead | 51 | 44 | 36 | 17 | 0 | 0 | Dead | 26 | 20 | 14 | 8 | 0 | 0 |
| **Tocilizumab** | | | | | | | **Placebo** | | | | | | |
| **Neutrophils Pct** | **BL** | **Day 3** | **Wk 1** | **Wk 2** | **Wk 4** | **Day 60** | **Lymphocyte Pct** | **BL** | **Day 3** | **Wk 1** | **Wk 2** | **Wk 4** | **Day 60** |
| Better | 224 | 189 | 190 | 147 | 102 | 133 | Better | 111 | 99 | 103 | 79 | 53 | 69 |
| Worse | 52 | 44 | 36 | 18 | 0 | 0 | Worse | 26 | 20 | 14 | 8 | 0 | 0 |
| **Tocilizumab** | | | | | | | **Placebo** | | | | | | |
| **Neutrophil By Lymphocyte** | **BL** | **Day 3** | **Wk 1** | **Wk 2** | **Wk 4** | **Day 60** | **Lymphocyte Pct** | **BL** | **Day 3** | **Wk 1** | **Wk 2** | **Wk 4** | **Day 60** |
| Survivors | 222 | 184 | 122 | 137 | 101 | 133 | Survivors | 108 | 98 | 60 | 77 | 52 | 69 |
| Dead | 51 | 44 | 24 | 17 | 0 | 0 | Dead | 26 | 20 | 9 | 8 | 0 | 0 |
| **Tocilizumab** | | | | | | | **Placebo** | | | | | | |
| **Lymphocyte Abs** | **BL** | **Day 3** | **Wk 1** | **Wk 2** | **Wk 4** | **Day 60** | **Lymphocyte Abs** | **BL** | **Day 3** | **Wk 1** | **Wk 2** | **Wk 4** | **Day 60** |
| Survivors | 227 | 190 | 190 | 148 | 103 | 137 | Survivors | 110 | 99 | 102 | 80 | 54 | 70 |
| Dead | 56 | 44 | 36 | 17 | 0 | 0 | Dead | 28 | 20 | 14 | 8 | 0 | 0 |
| **Tocilizumab** | | | | | | | **Placebo** | | | | | | |
| **Neutrophil Abs** | **BL** | **Day 3** | **Wk 1** | **Wk 2** | **Wk 4** | **Day 60** | **Lymphocyte Abs** | **BL** | **Day 3** | **Wk 1** | **Wk 2** | **Wk 4** | **Day 60** |
| Survivors | 229 | 190 | 190 | 148 | 103 | 133 | Survivors | 112 | 99 | 103 | 79 | 53 | 70 |
| Dead | 56 | 44 | 36 | 18 | 0 | 0 | Dead | 28 | 20 | 14 | 8 | 0 | 0 |
